# Generation of fate patterns via intercellular forces

**DOI:** 10.1101/2021.04.30.442205

**Authors:** Hayden Nunley, Xufeng Xue, Jianping Fu, David K. Lubensky

## Abstract

Studies of fate patterning during development typically emphasize cell-cell communication via diffusible signals. Recent experiments on monolayer stem cell colonies, however, suggest that mechanical forces between cells may also play a role. These findings inspire a model of mechanical patterning: fate affects cell contractility, and pressure in the cell layer biases fate. Cells at the colony boundary, more contractile than cells at the center, seed a pattern that propagates via force transmission. In agreement with previous observations, our model implies that the width of the outer fate domain depends only weakly on colony diameter. We further predict and confirm experimentally that this same width varies non-monotonically with substrate stiffness. This finding supports the idea that mechanical stress can mediate patterning in a manner similar to a morphogen; we argue that a similar dependence on substrate stiffness can be achieved by a chemical signal only if strong constraints on the signaling pathway’s mechanobiology are met.

---

Embryonic development depends on fate specification events in which initially equivalent cells differentiate in a spatially controlled manner [1–6]. A key example is neural induction: in the vertebrate ectoderm, a strip of cells assumes the neural plate (NP) fate as cells on either side differentiate into neural plate border (NPB) (Fig. 1(a)) [7–11]. *In vivo*, this NP-NPB pattern is thought to be induced primarily by the secretion of diffusible chemicals from adjacent tissues [8, 9]. Here, we show that *in vitro* experiments in the absence of such exogenous chemical gradients can be explained by a model where intercellular forces mediate the formation of the same pattern.

**FIG. 1.**
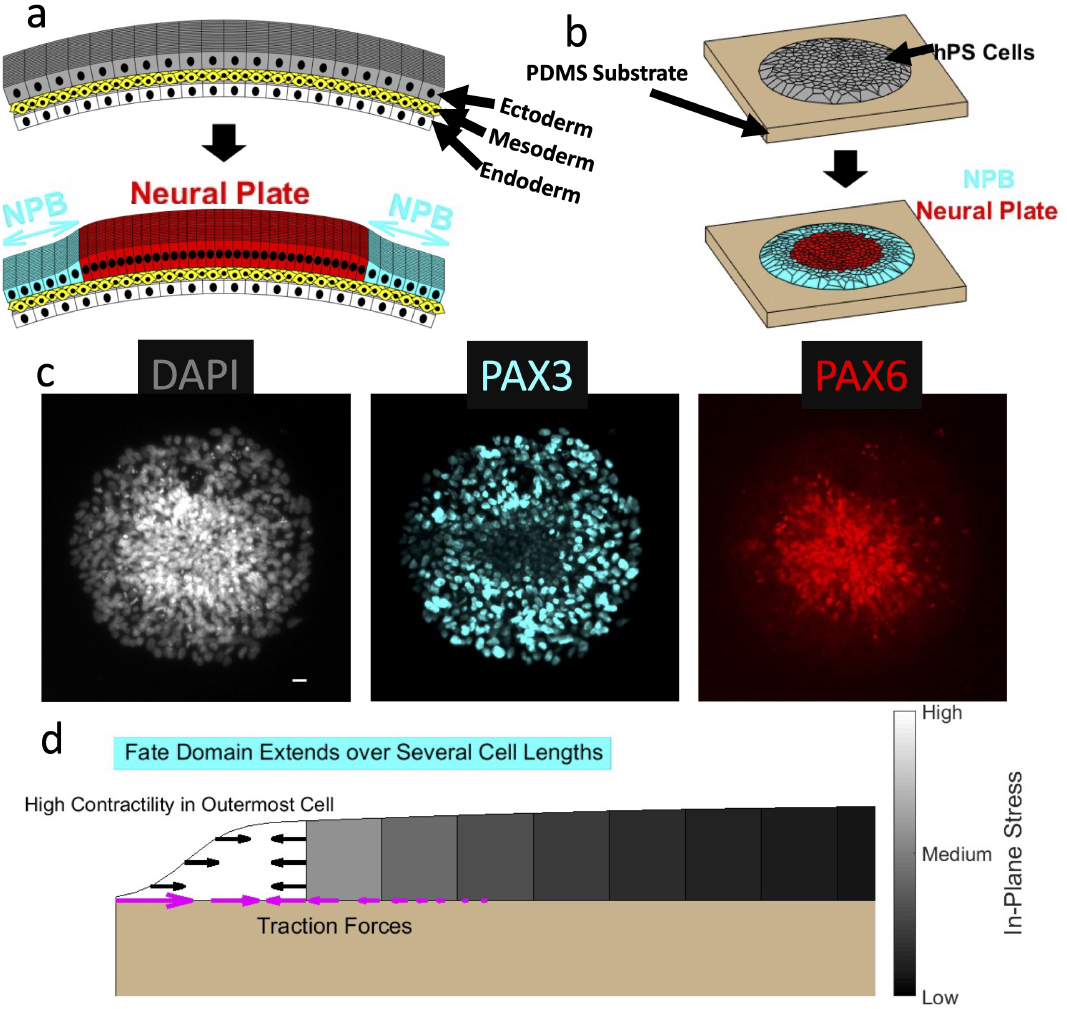
(a) Schematic of neural induction *in vivo*. A strip of cells in the embryonic ectoderm differentiates into NP (red). Cells on either side form the NPB (cyan). (b) Schematic of neural induction *in vitro*. hPS cells bind to circular regions of elastic substrate (tan). We stain for NP (red) and NPB (cyan). Black lines indicate Voronoi tessellations of nuclear positions before and after neural induction initiation. (c) Representative immunofluorescence images of colony at day 9. DAPI counterstains nuclei. PAX3 and PAX6 staining label nuclei of NPB cells and NP cells, respectively. Scale bar, 20 µm. (d) Schematic of cell-cell communication via forces. Edge cells generate contractile forces (black arrows), perceived by other cells, which in turn respond by forming a fate pattern. Importantly, cell-substrate traction forces (magenta) shape the spatial pattern of intercellular forces. Cyan: size of NPB domain, extending beyond outermost cell.

Recent experiments have probed the NP-NPB specification process *in vitro* (Fig. 1(b)) [11]: Human pluripotent stem (hPS) cell colonies bind to micro-patterned circular regions on elastic substrates. Treated with a spatially uniform neural induction medium, hPS cells near the colony center differentiate into NP; they are surrounded by a ring of NPB cells extending to the colony border (Fig. 1(b)). Since motion of individual cells is limited, patterned differentiation in these hPS cell colonies must rely on cells’ sensing their own position relative to the colony boundary [3, 11]. This differentiation does not depend on endogenous expression of BMP4 or NOGGIN, chemical signals relevant for establishing graded BMP signaling *in vivo* during neural induction [10, 11], but is affected by mechanical stretching. These findings suggest that mechanical stresses between stationary cells may act like a morphogen to generate NP-NPB patterning *in vitro*.

Inspired by these results, here we build a phenomenological model of NP-NPB pattern formation based on a coupling between mechanical pressure and cell fate. Our model is distinct from mechanical patterning models that require chemical advection [12, 13] or cell motion [14–16]. Although it shares some mathematical features with models of strain-activated contractility [15, 17], our model differs by explicitly considering cell fate and by operating in a parameter regime where it produces a single stationary NPB domain rather than traveling waves. In our model, intercellular forces act like a morphogen to determine the NPB domain size. The substrate’s mechanical properties, rather than biasing individual cells’ fates directly [18], control the intercellular propagation of information via forces [19, 20].

To model intercellular force transmission, we employ existing formalisms [21–23] that treat cell layers as thin, actively contractile elastic sheets bound to passive elastic substrates. Cell layers “leak” stress into the substrate as they deform it [21, 23]. In one dimension on thin substrates, force balance couples the cell layer’s in-plane stress *σ* to the substrate displacement *u* as:

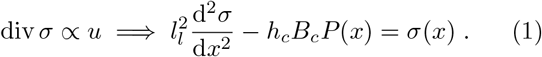

(In two dimensions, the scalars *σ* and *u* become the tensor 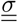 and vector 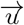, respectively.) −*h*_*c*_*B*_*c*_*P* (*x*) is an active stress, with the target strain *P* (*x*) specifying the strain at which cells are stress-free [23]. *h*_*c*_ and *B*_*c*_ are cell height and elastic modulus, respectively, and *l*_*l*_ is a length scale set by a balance of the cell layer and substrate stiffnesses [20–25]; if the compliance of cell-substrate adhesions can be neglected, 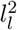 is inversely proportional to substrate stiffness [21–23].

To describe our system, we couple *σ* to a phenomenological fate variable *w*. Similar approaches to fate specification driven by chemical signals have had considerable success [26, 27]. Our model of mechanics-guided neural induction relies on two experimental observations: NPB cells are more contractile than NP cells, and extensile stress biases cells to the NPB fate [11]. These findings imply that our equations have couplings whose signs are inconsistent with a Turing-like, linear instability [5, 28]; instead, we propose that NPB domain formation is driven by forces from highly contractile cells at the colony boundary [29, 30]. Our model reproduces the observation that the NPB domain width is insensitive to colony diameter and correctly predicts that this width depends non-monotonically on substrate stiffness.

## Evidence for fate-stress feedback

To study the patterning of NP and NPB, we seed hPS cells on an elastic substrate or on a micropost array with circular micropatterned regions and induce neural differentiation as described [11]. Since BMP-SMAD and WNT signaling are known to play a role in neural induction [7], we initiate differentiation by supplying a medium that contains dual SMAD inhibitors [11, 31], supplemented with a *β*-catenin stabilizer that actives WNT [32, 33]. We label NP and NPB cells by immunostaining nuclei for PAX6 and PAX3, respectively (Fig. 1(c)) [11].

Based on our hypothesis that coupling fate to mechanical stimulus could generate patterning, we test experimentally whether cell fate regulates active stress and whether mechanical stress regulates fate. To quantify how cell fate influences active stress generation, we measure contractile forces exerted by cells on the substrate using a micropost force sensor (Fig. 2(a-d)) [11, 23, 34– 36]. Averaging radial post deflections at a given distance from the colony center, we determine which regions of the colony extend and which contract (Fig. 2(c)). Fits of these deflections to our mechanical model reveal three distinct regions: a non-contractile central region surrounded by two concentric contractile regions (Fig. 2(c); [37]). The non-contractile region coincides with the region that expresses PAX6, and the two contractile regions together correspond to the region that expresses PAX3 [11]. We propose an explanation for the unexpected presence of two mechanically distinct NPB regions below and in Section 3 of [37].

**FIG. 2.**
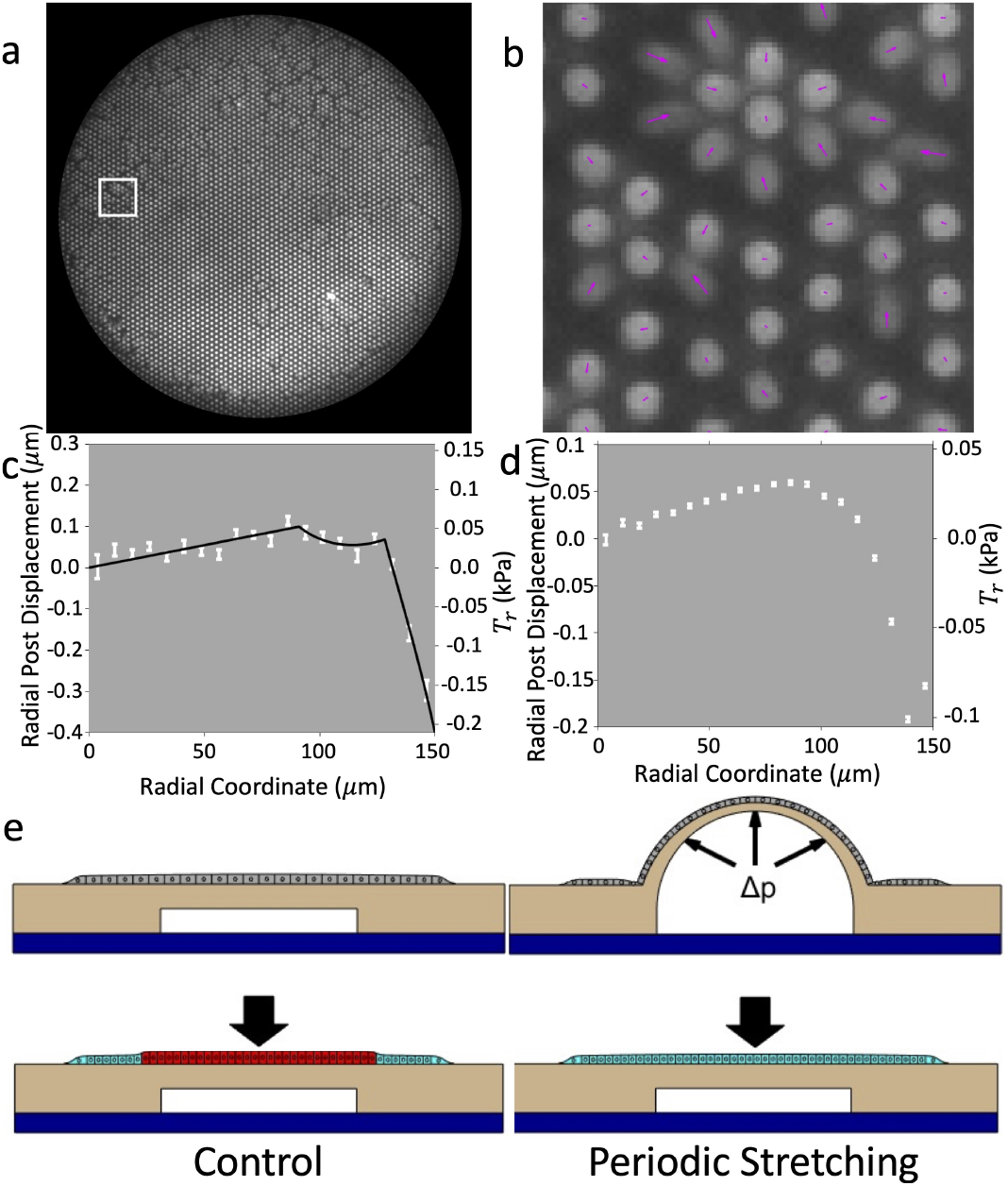
(a) Two days after neural induction initiation, we measure cell-substrate forces via micropost displacements. Colony diameter, 300 µm. (b) Posts from white box in (a). Cells displace posts (magenta arrows). Post diameter, 2.2 µm; post spacing, 4.0 µm. (c) Concentrically averaged radial post displacements (from (a)) and radial traction stress *Tr* versus radial coordinate for a representative colony. Black line: three-domain fit to Eq. 4 [37]. We interpret the outer-most region as a ring of spread cells at the colony edge, the intermediate region as a contractile ring of bulk cells, and the inner region as non-contractile cells. (d) Post displacement profiles averaged over *n* = 42 colonies. Since the elbow-like feature’s position (see (c)) shifts from sample to sample, this feature smooths out on average. (e) Schematic of control experiments (left) in which hPS cell colonies differentiate into NP (red) and NPB (cyan) domains. Schematic of experiment (right) in which a microfluidic chamber (below colony) stretches cell layer. Stretching biases cells near the colony center to the NPB fate [11].

To test whether mechanical stress itself biases fate, we stretch the colony central region during differentiation. Stretching biases the central region to be PAX3+ (*i*.*e*., positive for PAX3 expression) rather than PAX6+ (Fig. 2(e)); similarly, when constrained to larger areas, single cells more often show prominent nuclear p-SMAD 1/5 (another marker of NPB fate) [11]. Micropost and stretching experiments thus reveal that NPB cells are more contractile than NP cells and that stretching biases cells to the NPB fate.

## Phenomenological model of mechanical fate patterning

Because we are interested in a binary fate decision, it is natural to map fate to a single variable *w* evolving in a bistable potential with minima corresponding to NP (*w* ≈ 0) and NPB (*w* ≈ 1) (Fig. 3(a)) [27]. In the spirit of including all allowed couplings at the lowest order, we make two additional simplifying assumptions: (1) active stress is proportional to *w*, 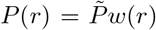; and (2) mechanical pressure 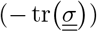 linearly biases *w*. Thus,

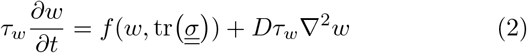

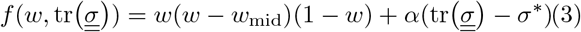

**FIG. 3.**
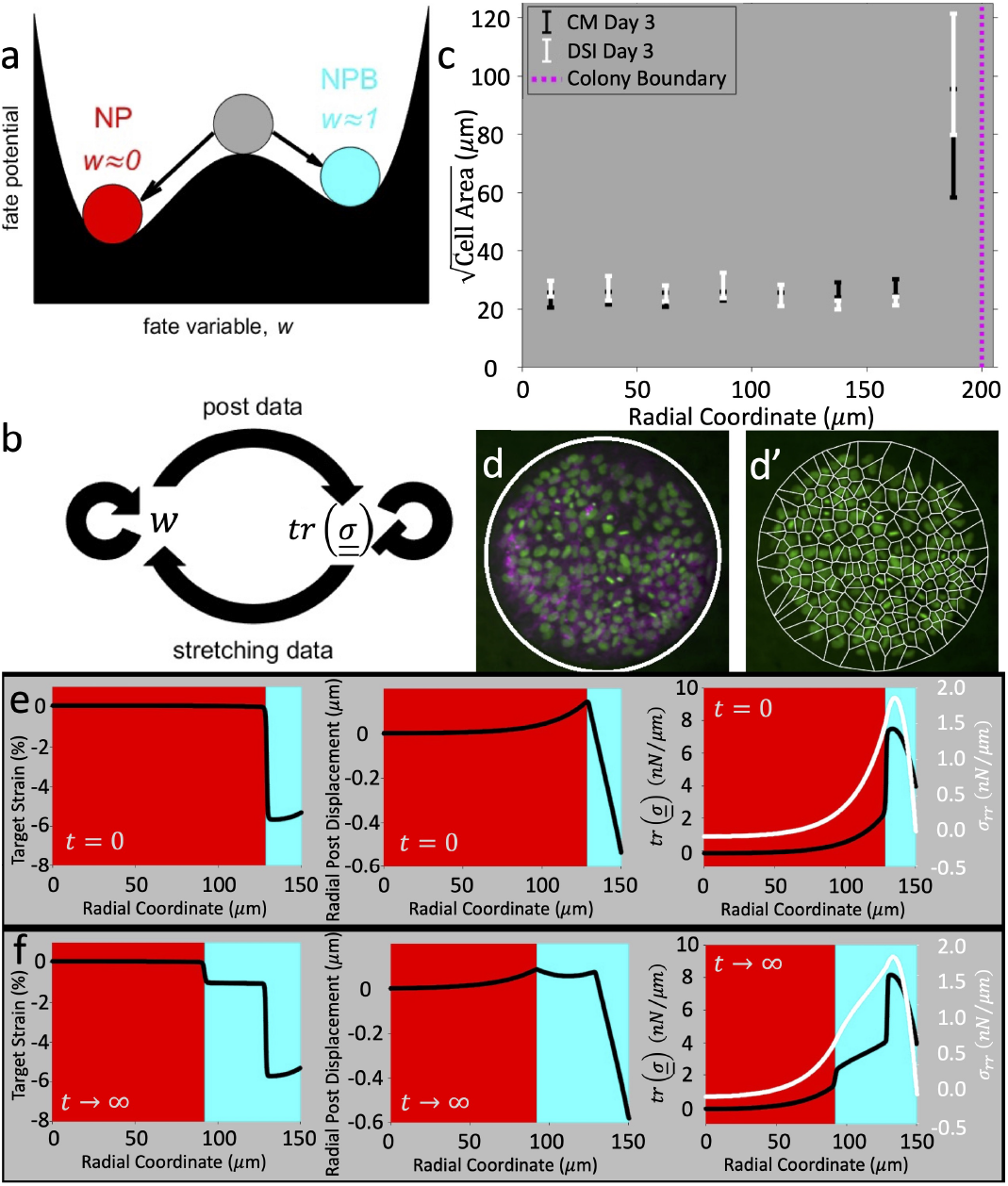
(a) Schematic of differentiation from unstable cell state (gray) to stable cell state (red or cyan). (b) Feedback diagram for fate *w* and tr 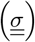. (c) Square root of cell area versus radial coordinate. In neural induction medium (DSI) and in conditioned medium (CM), cell area is large near colony boundary (magenta line) relative to colony bulk. (d) Colony, immuno-stained for DAPI (green) and E-Cad (magenta), at day 3 in DSI. White circle: micropatterned colony edge. (d’) Same colony as in (d) with tessellation over nuclei. Colony diameter, 400 µm. (e, f) Initial condition (e) and fixed point (f) for the model (Eqs. 2–4). All parameters, except for *α* and *w*_mid_ (Table S6 [37]) are from a fit to micropost data in Fig. 2(a-c). We plot target strain (left), post displacement (center), and stresses (left). Initially, outermost cell, which has large area, is contractile (cyan background, e). Highly contractile cells at colony boundary generate stress in neighbors. The fate boundary (between cyan and red, e-f) moves inward until stress at fate boundary reaches coexistence stress.

Here *α* > 0 and 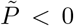 are constant. The signs of all couplings are determined by the micropost data and the stretching data, together with the assumption of bistability (Fig. 3(b)). *τ*_*w*_ is the time scale of fate evolution, and *w*_mid_ determines the asymmetry of the potential in the absence of stress feedback (Fig. 3(a)). *σ*^*^ is a constant that sets the width of the NPB domain. To regularize spatial variation of *w*, we introduce a diffusion term in Eq. (2); 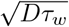 determines the width of the NP-NPB boundary. We impose no-flux boundary conditions on *w* at the free boundary to avoid artificially driving patterning in a stress-independent manner.

To calculate the in-plane mechanical stress, we generalize Eq. (1) for a contractile cell layer on a thin substrate to more than one dimension [37]:

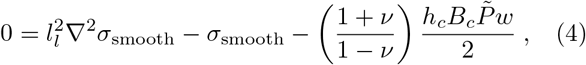

where *ν* is the Poisson’s ratio of the cell layer and 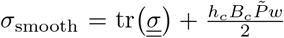. We assume that the normal component of 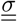 vanishes at the free boundary.

Since the fate-stress couplings are of common sign in our model (Fig. 3(b)), it does not exhibit any Turing-like, linear instability [5, 28]. Instead, we hypothesize that formation of an NPB domain is driven by mechanical signals from cells at the edge of the colony, which are known in related systems to sense the colony boundary and thereby to develop distinct properties [29, 30, 38–40]. Indeed, immunostaining of nuclei and adherens junctions during differentiation reveals that boundary cells spread to a larger area per cell than bulk (*i*.*e*., non-boundary) cells (Fig. 3(c-d’)). The region of boundary cells coincides with the outermost region of highest cell contractility whose existence we inferred from micropost measurements (Fig. 2(c), see Section 3 of [37]). We introduce boundary inhomogeneity by making the factor relating contractility to cell fate depend on position:

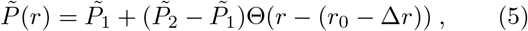

where 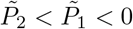 are constants, *r*_0_ is the colony radius, Δ*r* is the radial extent of the boundary cells, and Θ is the Heaviside step function.

With this choice of 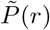, a ring of highly contractile cells at the colony boundary generates in-plane stresses in neighboring cells (Fig. 3(e)), which are then driven to increase their own *w*, upregulating their contractility. The fate boundary moves inward until the stress at the fate boundary equals a coexistence stress proportional to *σ*^*^ that must fall within a particular range in order for a stable pattern of both NP and NPB to exist (Fig. 3(f)). We can analytically determine the position of the fate boundary in the limit that the fate boundary is much sharper than *l*_*l*_, *i*.*e*., 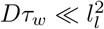 [37, 41–43].

Our phenomenological model, based on experimentally motivated assumptions, reproduces measured traction forces (compare example in Fig. 2(c) to Fig. 3(f)). To further test the model, we next compare its predictions for the NPB domain width as a function of key parameters with experiments.

## NPB domain width does not depend on colony diameter, depends non-monotonically on substrate stiffness

We previously reported that, as the colony diameter varies from 300 µm to 800 µm, the width of the NPB domain does not change significantly (Fig. 4(a, b)). Our model recapitulates this approximate independence of NPB domain width on colony diameter. (See [37] for the extension of our model to allow for a finite substrate thickness that we use to generate the model predictions in Fig. 4).

**FIG. 4.**
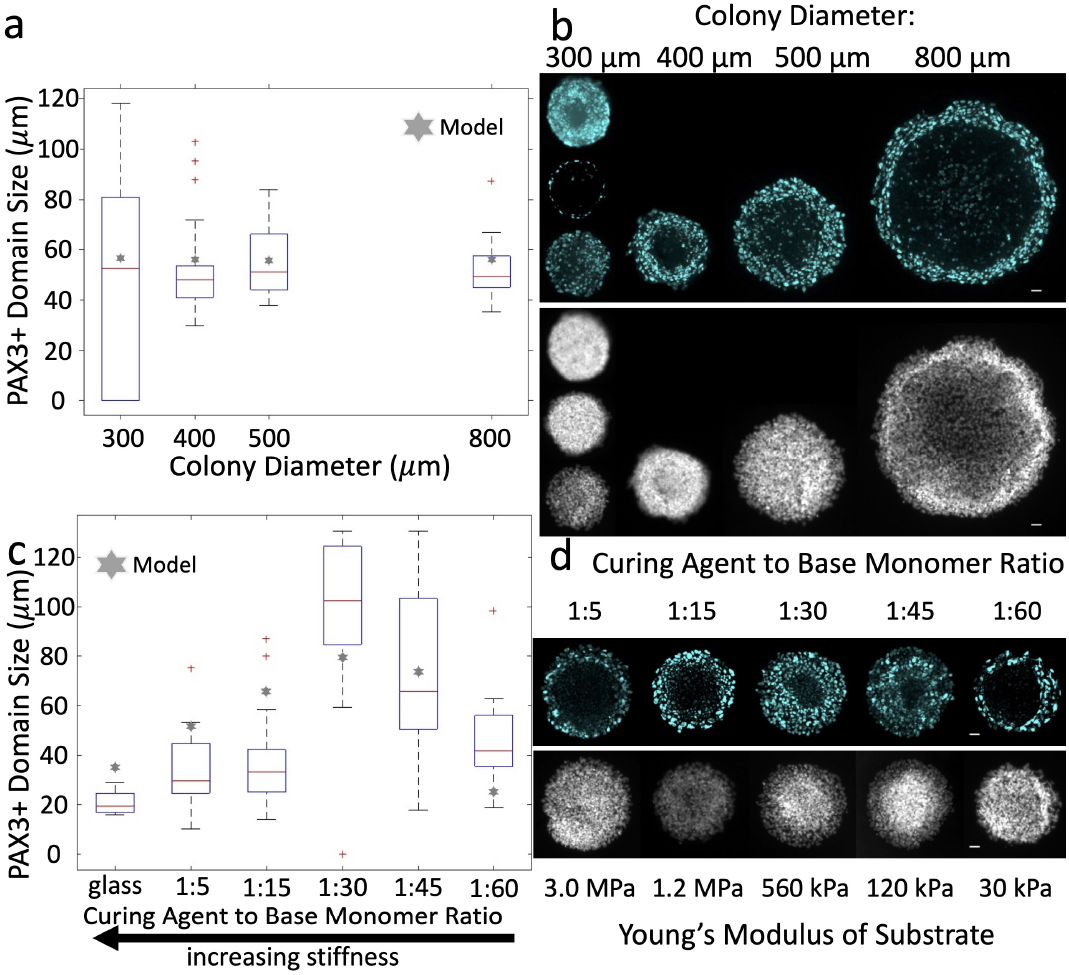
(a) NPB domain size versus colony diameter. The median width of the NPB domain (red lines; blue boxes give 25th and 75th percentile) is approximately independent of colony diameter. Red points: outliers. (b) Representative examples of colonies immuno-stained for PAX3 (cyan) and DAPI (gray). (c) NPB domain size versus substrate stiffness, showing non-monotonic dependence (colony diameter=400 µm; see Methods [37] for glass colony conditions). By changing the curing agent to base monomer ratio used in preparing the PDMS substrate, we alter its stiffness over orders of magnitude (with stiffer substrates corresponding to larger ratios). The substrate thickness also varies more mildly with the same ratio, an effect that we account for in our model calculations (stars), which employ a modification of Eqs. 2–4 that includes finite substrate thickness [37]. (d) Examples of cell colonies immunostained for PAX3 (cyan) and DAPI (gray). Scale bars, 40 µm.

We next consider how NPB domain width varies with substrate stiffness. Because the length scale *l*_*l*_ depends inversely on substrate stiffness, we might expect that the NPB domain grows as the substrate becomes softer. Strikingly, however, our model instead predicts a non-monotonic dependence of NPB domain size on stiffness (Fig. 4(c)). This behavior arises because, although the stress exerted by the contractile boundary cells decays exponentially away from the boundary with a lengthscale that decreases as the substrate becomes stiffer, the prefactor in front of this exponential decay instead *increases* along with substrate stiffness; for example, for the simplifed case of a delta function stress source in one dimension, we would have 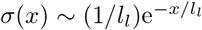. (A similar effect is at the origin of a previously reported prediction that the force between cells in a monolayer can vary nonmonotonically with substrate stiffness [20].)

Experiments confirm our model’s prediction of a non-monotonic dependence on substrate stiffness (Fig. 4(c, d)). As the substrate stiffness decreases from 3 MPa to 30 kPa at a constant colony diameter of 400 µm, the NPB domain width increases from ≈ 40 *µ*m to ≈ 100 *µ*m before decreasing to ≈ 25 µm.

## Discussion

In this paper, we have argued that mechanical stress may act like a morphogen, patterning cell fates in a position-dependent manner, in some hPS cell systems [44–47]. In particular, we find experimentally that the width of the NPB fate domain in our colonies depends non-monotonically on the substrate stiffness, as predicted by a model with a mechanical morphogen. In agreement with this picture, micropost deflection measurements indicate that the magnitude of the in-plane mechanical stresses in our colonies is comparable to stresses that have been shown to induce mechanotransduction responses in other systems [48].

Nonetheless, it is important to ask whether the observed non-monotonic dependence on substrate stiffness could also be produced by a conventional morphogen (most likely a member of the WNT family [7]). Morphogen concentrations of course usually obey a screened diffusion equation like Eq. (1) (with *σ* replaced by the morphogen concentration). Thus, if cells responded autonomously to changes in substrate stiffness by varying the diffusion length in an appropriate way, this would lead to behavior like that observed in Fig. 4(c). One obvious way for cell-autonomous changes to affect the diffusion length would be for rates of ligand internalization and degradation to depend on the substrate stiffness sensed by the cells. Importantly, however, this dependence would have to be very strong to reproduce our experimental results; we estimate that the ligand degradation rate would need to vary by at least a factor of 100 over the range of stiffnesses tested. Thus, although we cannot categorically rule out a non-mechanical origin for our observations, we can say that any signaling by a diffusible morphogen that mimicked the predictions of the mechanical model would have to obey strong and specific constraints. Direct tests of such an alternative mechanism await further investigation.

Finally, it is worth emphasizing that, whereas chemical morphogens do not necessarily play a role in our *in vitro* experiments, BMP proteins almost certainly are an important factor in the NP-NPB fate decision *in vivo* [8–10, 45]. To fully understand the functional importance of the mechanical effects uncovered here, it will be necessary to tease out how mechanical factors interact with diffusible morphogens *in vivo* to promote robust patterning and morphogenesis.

This work was supported by the National Science Foundation (CMMI 1917304 to J.F. and DGE 1256260 to H.N.), by pilot project funding from the NSF-Simons Center for Quantitative Biology (Simons Foundation/SFARI 597491-RWC and the National Science Foundation 1764421 to J.F. and D.K.L.), and by the Michigan-Cambridge Collaboration Initiative (to J.F.).

## Supporting information

Full Supplemental Materials

Expressions for Model 3, Section 3 of Supplemental Materials

## References

[1] C. J. Chan, C. P. Heisenberg, and T. Hiiragi, Curr. Biol. 27, R1024 (2017).

[2] W. Driever and C. Nusslein-Volhard, Cell 54, 95 (1988).

[3] T. Gregor, D. W. Tank, E. F. Wieschaus, and W. Bialek, Cell 130, 153 (2007).

[4] E. Hannezo and C. P. Heisenberg, Cell 178, 12 (2019).

[5] A. M. Turing, Bull. Math. Biol. 52, 153 (1990).

[6] C. H. Waddington and International Union of Biological Sciences., Towards a theoretical biology. An IUBS symposium (Aldine Pub. Co., Chicago, 1968).

[7] G. Britton, I. Heemskerk, R. Hodge, A. A. Qutub, and A. Warmflash, Development 146 (2019).

[8] I. Martyn, T. Y. Kanno, A. Ruzo, E. D. Siggia, and A. H. Brivanlou, Nature 558, 132 (2018).

[9] H. Spemann and H. Mangold, Int. J. Dev. Biol. 45, 13 (2001).

[10] P. A. Wilson, G. Lagna, A. Suzuki, and A. Hemmati-Brivanlou, Development 124, 3177 (1997).

[11] X. Xue et al., Nat. Mater. 17, 633 (2018).

[12] J. S. Bois, F. Jülicher, and S. W. Grill, Phys. Rev. Lett. 106, 028103 (2011).

[13] P. Recho, A. Hallou, and E. Hannezo, Proc. Natl. Acad. Sci. U S A 116, 5344 (2019).

[14] J. D. Murray and G. F. Oster, J. Math. Biol. 19, 265 (1984).

[15] J. D. Murray and G. F. Oster, IMA J. Math. Appl. Med. Biol. 1, 51 (1984).

[16] A. E. Shyer, A. R. Rodrigues, G. G. Schroeder, E. Kassianidou, S. Kumar, and R. M. Harland, Science 357, 811 (2017).

[17] G. M. Odell, G. Oster, P. Alberch, and B. Burnside, Dev. Biol. 85, 446 (1981).

[18] A. J. Engler, S. Sen, H. L. Sweeney, and D. E. Discher, Cell 126, 677 (2006).

[19] V. Maruthamuthu, B. Sabass, U. S. Schwarz, and M. L. Gardel, Proc. Natl. Acad. Sci. U S A 108, 4708 (2011).

[20] A. F. Mertz et al., Proc. Natl. Acad. Sci. U S A 110, 842 (2013).

[21] S. Banerjee and M. C. Marchetti, EPL (Europhysics Letters) 96, 28003 (2011).

[22] S. Banerjee and M. C. Marchetti, Phys. Rev. Lett. 109, 108101 (2012).

[23] C. M. Edwards and U. S. Schwarz, Phys. Rev. Lett. 107, 128101 (2011).

[24] P. W. Oakes, S. Banerjee, M. C. Marchetti, and M. L. Gardel, Biophys. J. 107, 825 (2014).

[25] E. N. Schaumann, M. F. Staddon, M. L. Gardel, and S. Banerjee, Mol. Biol. Cell 29, 2835 (2018).

[26] F. Corson, L. Couturier, H. Rouault, K. Mazouni, and F. Schweisguth, Science 356 (2017).

[27] F. Corson and E. D. Siggia, Proc. Natl. Acad. Sci. U S A 109, 5568 (2012).

[28] M. Cross and H. Greenside, Pattern formation and dynamics in nonequilibrium systems (Cambridge University Press, Cambridge, UK ; New York, 2009).

[29] E. Narva, A. Stubb, C. Guzman, M. Blomqvist, D. Balboa, M. Lerche, M. Saari, T. Otonkoski, and J. Ivaska, Stem Cell Reports 9, 67 (2017).

[30] K. A. Rosowski, A. F. Mertz, S. Norcross, E. R. Dufresne, and V. Horsley, Sci. Rep. 5, 14218 (2015).

[31] S. M. Chambers, C. A. Fasano, E. P. Papapetrou, M. Tomishima, M. Sadelain, and L. Studer, Nat. Biotechnol. 27, 275 (2009).

[32] Y. Mica, G. Lee, S. M. Chambers, M. J. Tomishima, and L. Studer, Cell Rep. 3, 1140 (2013).

[33] J. Tchieu, B. Zimmer, F. Fattahi, S. Amin, N. Zeltner, S. Chen, and L. Studer, Cell Stem Cell 21, 399 (2017).

[34] J. Fu, Y. K. Wang, M. T. Yang, R. A. Desai, X. Yu, Z. Liu, and C. S. Chen, Nat. Methods 7, 733 (2010).

[35] A. Saez, A. Buguin, P. Silberzan, and B. Ladoux, Biophys. J. 89, L52 (2005).

[36] Y. Sun et al., Nat. Mater. 13, 599 (2014).

[37] See Supplemental Material at [URL will be inserted by publisher] for estimation of in-plane stress in colonies, estimation of cellular mechanical properties, simulations of tissue patterning on substrates of finite thickness, and calculations based on the domain-wall approximation.

[38] J. D. Amack and M. L. Manning, Science 338, 212 (2012).

[39] M. L. Manning, R. A. Foty, M. S. Steinberg, and E. M. Schoetz, Proc. Natl. Acad. Sci. U S A 107, 12517 (2010).

[40] I. Martyn, A. H. Brivanlou, and E. D. Siggia, Development 146 (2019).

[41] C. B. Muratov and S. Y. Shvartsman, Physica D: Non-linear Phenomena 186, 93 (2003).

[42] C. B. Muratov and V. V. Osipov, Physical Review E 53, 3101 (1996).

[43] W. van Saarloos, Physics Reports 301, 9 (1998).

[44] P. Xia, D. Gutl, V. Zheden, and C. P. Heisenberg, Cell 176, 1379 (2019).

[45] J. M. Muncie, N. M. E. Ayad, J. N. Lakins, X. Xue, J. Fu, and V. M. Weaver, Dev. Cell 55, 679 (2020).

[46] N. Desprat, W. Supatto, P. A. Pouille, E. Beaurepaire, and E. Farge, Dev Cell 15, 470 (2008).

[47] T. Brunet et al., Nat. Commun. 4, 2821 (2013).

[48] T. Das, K. Safferling, S. Rausch, N. Grabe, H. Boehm, and J. P. Spatz, Nat. Cell Biol. 17, 276 (2015).

[49] R. J. LeVeque, Finite difference methods for ordinary and partial differential equations : steady-state and time-dependent problems (Society for Industrial and Applied Mathematics, Philadelphia, PA, 2007).

[50] U. M. Ascher, S. J. Ruuth, and T. R. W. Brian, SIAM Journal on Numerical Analysis 32, 797 (1995).

[51] X. Trepat, M. R. Wasserman, T. E. Angelini, E. Millet, D. A. Weitz, J. P. Butler, and J. J. Fredberg, Nature Physics 5, 426 (2009).

[52] A. F. Mertz, S. Banerjee, Y. Che, G. K. German, Y. Xu, C. Hyland, M. C. Marchetti, V. Horsley, and E. R. Dufresne, Phys. Rev. Lett. 108, 198101 (2012).

[53] D. T. Tambe et al., Nat. Mater. 10, 469 (2011).

[54] M. Uroz, S. Wistorf, X. Serra-Picamal, V. Conte, M. Sales-Pardo, P. Roca-Cusachs, R. Guimera, and X. Trepat, Nat. Cell Biol. 20, 646 (2018).

[55] M. Vishwakarma, J. Di Russo, D. Probst, U. S. Schwarz, T. Das, and J. P. Spatz, Nature Communications 9, 3469 (2018).

[56] S. Banerjee and M. Cristina Marchetti, New Journal of Physics 15, 035015 (2013).

[57] C. Perez-Gonzalez, R. Alert, C. Blanch-Mercader, M. Gomez-Gonzalez, T. Kolodziej, E. Bazellieres, J. Casademunt, and X. Trepat, Nat. Phys. 15, 79 (2019).

[58] A.E. Raftery, Sociological Methodology 25, 111 (1995).

[59] E. J. Wagenmakers, Psychon. Bull. Rev. 14, 779 (2007).

[60] L. Wasserman, Journal of Mathematical Psychology 44, 92 (2000).

[61] M. Ghibaudo, A. Saez, L. Trichet, A. Xayaphoummine, J. Browaeys, P. Silberzan, A. Buguin, and B. Ladoux, Soft Matter 4, 1836 (2008).

[62] B. S. Kerner and V. V. Osipov, Autosolitons : a new approach to problems of self-organization and turbulence (Kluwer Academic, Dordrecht ; Boston, 1994), Fundamental theories of physics, 61.

[63] A. S. Mikhailov, Foundations of synergetics (Springer, Berlin ; New York, 1994), 2nd edn., Springer series in synergetics, 51-.

