## Supplementary material for "Generation of fate patterns via intercellular forces": Full Supplemental Materials

**This PDF file includes:**

Materials and Methods

Supplemental Text

Figs. S1 to S6

Tables S1 to S9

Captions for Figs. S1 to S6

Captions for Tables S1 to S9

#### Materials and methods

##### Cell culture

H1 human embryonic stem cells (WiCell) were maintained on mitotically inactive mouse embryonic fibroblasts (MEFs; Thermo Fisher Scientific) in KSR medium consisting of DMEM/F12 (GIBCO) with 20% Knockout Serum Replacement (GIBCO), 0.1 mM β-mercaptoethanol (GIBCO), 2 mM Glutamax (GIBCO), 1% non-essential amino acids (GIBCO), and 4 ng/ml human recombinant basic fibroblast (bFGF, Thermo Fisher Scientific). The cells were passaged using the STEMPRO EZPassage Disposable Stem Cell Passaging Tool (Invitrogen) as described previously [11]. The cell line was tested negative for mycoplasma contamination using LookOut Mycoplasma PCR Detection Kit (Sigma-Aldrich).

To induce neuroectoderm differentiation, cells were seeded on micropatterned PDMS substrate in KSR medium on day 0 at a density of 20,000 cells/$cm^2$. 10 μM ROCK inhibitor Y-27632 (R&D Systems) was added to prevent cell dissociation-induced apoptosis. On day 1, cell culture medium was replaced with fresh KSR medium without Y-27632. Cells were cultured in neural induction medium consisting of KSR medium supplemented with TGF- β inhibitor SB 431542 (10 μM; Stemcell Technologies) and BMP4 inhibitor LDN 193189 (500 nM; Stemcell Technologies) from day 2 to day 9. On day 3, CHIR99021 (3 μM; Stemcell Technologies) was added to the neural induction medium and was withdrawn on day 4.

##### Microcontact printing

PDMS stamps were fabricated using soft lithography from silicon molds, as described previously [34]. To generate micropatterned adhesive islands, a thin layer of PDMS was first spin-coated onto a glass coverslip at 1000 r.p.m. for 30 s and cured at 110 °C for 24 h before use. Base-to-curing-agent ratios of 5:1, 15:1, 30:1, 45:1 and 60:1 were used. PDMS stamps were soaked in a vitronectin solution (20 μg/ml, Stemcell Technologies) for 1 h. PDMS spin-coated coverslips were activated using ultraviolet ozone and then placed in contact with Vitronectin-coated PDMS stamps. Coverslips were then immersed in 0.2% pluronics solutions (BASF) for 30 min and washed with PBS before cell seeding. PDMS stamps containing circular patterns with diameters of 300, 400, 500 and 800 μm were used.

To determine the Young's modulus of a PDMS substrate of a given base-to-curing-agent ratios, an Instron tensile testing machine (Illinois Tool Works Inc) was used. PDMS membrane thicknesses were measured via a stylus profilometer (Dektak XT, Bruker Corporation). See Table S5 for measured properties of the substrates.

**Immunocytochemistry**

hESCs were fixed in 4% paraformaldehyde for 30 min and permeabilized in 0.1% sodium dodecyl sulfate (SDS, Sigma Aldrich) for another 30 min. Cells were then incubated in 10% goat serum (Thermo Fisher Scientific) for 1 h to block nonspecific binding, followed by primary and secondary antibodies for 1 h. 4,6-diamidino-2-phenylindole (DAPI, Thermo Fisher Scientific) was used for counterstaining cell nuclei. All primary antibodies are listed in Table S7.

**Traction force measurements**

Traction force was measured using PDMS micropost arrays (PMAs) as described previously [34]. Vitronectin was first coated onto the top surface of PMAs using microcontact printing. Then PMAs were incubated in 5μg/ml DiI (Invitrogen) for 1 h to label vitronectin-coated PMAs. PMAs were then immersed in 0.2% pluronics solutions (BASF) for 30 min and washed with PBS before cell seeding. Live cell imaging was performed using Zeiss Axio Observer Z1 inverted epifluorescence microscope enclosed in the XL S1 incubator.

**Determination of $PAX3^+$ domain width from colonies stained for DAPI and PAX3**

We threshold the DAPI image to identify the region with nuclei. We, then, morphologically close the thresholded DAPI image. In this closed image, we identify all pixels that delineate the boundary of the colony. Then, for each pixel within the colony, we compute its shortest distance to the colony boundary. To obtain a PAX3 intensity profile as a function of distance from the colony boundary, we average over pixels that are approximately equidistant from the colony boundary (between some distance $|\vec{r}|$ and $|\vec{r}| - \Delta r$ from the colony boundary). We choose the averaging window's width to be approximately ten microns.

Then, for each colony, we use its PAX3 intensity profile as a function of distance from the colony boundary to calculate the width of the $PAX3^+$ domain. For each experimental condition indexed by $i$ (fixed substrate mechanical properties and fixed colony diameter), we compute the average maximal PAX3 intensity (called $\overline{PAX3^{max}}_i$) and the average minimal PAX3 (called $\overline{PAX3^{min}}_i$). From these averages, we compute a midpoint (called $\overline{PAX3^{mid}}_i \equiv \frac{\overline{PAX3^{max}}_i + \overline{PAX3^{min}}_i}{2}$). Then, for each experimental condition $i$, we loop through all colonies, calculating the distance from the colony boundary to the position where the PAX3 intensity profile crosses $\overline{PAX3^{mid}}_i$. That distance to the crossing point for each colony is its PAX3$^+$ domain width. Notably, this analysis method can identify which colonies are entirely PAX3$^-$, entirely PAX3$^+$, as well as patterned colonies (that have both PAX3$^-$ and PAX3$^+$ regions).

**Determination of in-plane nuclear density from colonies stained for DAPI and E-Cadherin**

We threshold and morphologically close the E-Cadherin image to identify the region that contains cells. (There is faint E-Cadherin even at the very boundary of the colony where cells lack adherens junctions on one side.) To the DAPI image, we apply a gaussian blur (with kernel smaller than the nuclear size) on the image (imgaussfilt in MATLAB 2016B, Mathworks, Natick, MA). Then, we look for regional maxima (imregionalmax in MATLAB 2016B, Mathworks, Natick, MA) in this blurred DAPI image; the regional maxima correspond to the approximate centers of the nuclei. We make minimal manual corrections in cases of misidentified nuclei.

Given the thresholded and morphologically closed E-Cadherin image as well as the location of individual nuclei, we would like to compute the in-plane nuclear density in the image region that actually contains cells. To do so, we count how many nuclear centers fall between $|\vec{r}|$ and $|\vec{r}| - \Delta r$ from the colony center as well as the total thresholded area (from the E-Cadherin image) that is between $|\vec{r}|$ and $|\vec{r}| - \Delta r$ from the colony center. We divide the total thresholded area (between $|\vec{r}|$ and $|\vec{r}| - \Delta r$) by the number of nuclei in the corresponding region. Repeating this process for concentric annuli, we generate plots as in Fig. 3C. See Fig. 3D for example image.

**Numerical methods for case of cell layer on substrate composed of microposts**

For the case of a cell layer bound to a substrate composed of microposts, we solve the equations for both fate and stress (Eqs. S1, S2, S19 for stripe geometry and Eqs. S1, S2, S29, S30 for disc geometry) via finite-differences [49]. We treat non-linear terms explicitly and linear terms implicitly in a first-order solver [50]. The spatial discretization is 0.05 μm, and the time discretization is on the order of $0.05\tau_w$ (see Eqs. S1, S2). For a discussion of the numerical methods for the case of a cell layer bound to a finite-thickness substrate, see SI Section 4.

**Experiments and corresponding modeling for cell layer on glass substrate**

The experimental data on glass in Fig. 4C are the results of experiments in Fig. S3 of [11]. The colony diameter for these experiments is 500 μm instead of 400 μm; however, we know that the NPB domain width is insensitive to colony diameter (especially when $l_l$ is much smaller than the colony diameter; see Fig. 4A and Section 6B). Thus, we include these glass data points in Fig. 4C to broaden the range of experimentally explored substrate stiffnesses. To model these experiments on glass, we assume that the glass stiffness is infinite relative to both the cell-substrate adhesions and to the cell layer itself; thus, the Eq. S67 becomes a simple screened diffusion equation in which $l_a$ is the relevant length scale.

**Supplemental text**

In this supplemental text, to build an understanding of potential mechanical-stress-guided neural induction *in vitro*, we present both analyses of experimental data (Sections 2, 3) and biophysical models (Sections 1, 4-6) inspired by these data.

Section 1 summarizes the main equations used throughout this study and their different limiting cases (several of which we consider in more depth in subsequent sections). In Section 2, by integrating over forces between the cell layer and the substrate, we estimate mechanical stresses in the cell layer. In Section 3, we fit traction force measurements to existing mechanical models of cell layers bound to elastic substrates; this fitting allows us to estimate the cell layer's mechanical properties, like its contractility.

Based on the results of these data analyses, in Sections 4-6 we analyze a simplified biophysical model: mechanical stress biases cell fate, and cell fate biases active stress generation. Section 4 considers this model for a cell layer bound to a continuous substrate of finite thickness and develops computational methods to solve it. Sections 5 and 6 present an analytically tractable model of fate patterning for the simpler case of a cell layer bound to a substrate composed of discrete microposts (or to a sufficiently thin substrate relative to the in-plane extent of the cell layer [22]); this model is also briefly described in the main text. In Section 5, we derive a differential equation for the cell layer's mechanical pressure. Using the differential equation from Section 5, we develop an analytical approximation for the size of the outer fate domain in Section 6. This analytical approximation reveals why the size of the outer fate domain depends only weakly on colony diameter but non-monotonically on the stiffness of the cell layer's substrate.

#### 1. Definition of relevant equations and variables

In this section, as a reference, we assemble in one place both the equation for fate evolution and the several variants of the mechanical force balance equation that we use in this manuscript. The biophysical motivation for the fate equation is explained in the main text and draws on previous experimental findings [11]; equations for the mechanical stress in cell layers are based on earlier theoretical studies [21-23]. We model the patterning of fate, represented as a scalar $w$, in a thin epithelial layer coupled to an elastic substrate. The fate value $w \approx 1$ is the neural plate border (NPB) fate, and the fate value $w \approx 0$ is the neural plate (NP) fate. The fate evolves under the influence of a bistable driving, a linear bias from the trace of the in-plane stress tensor $\underline{\underline{\sigma}}$, and a diffusive term (which regularizes spatial variation):

$$\tau_w \frac{\partial w}{\partial t} = f\left(w, \mathrm{tr}\left(\underline{\underline{\sigma}}\right)\right) + D\tau_w \nabla^2 w. \qquad \text{[Eq. S1]}$$

$$f\left(w, \mathrm{tr}\left(\underline{\underline{\sigma}}\right)\right) = w(w - w_{mid})(1 - w) + \alpha\left(\mathrm{tr}\left(\underline{\underline{\sigma}}\right) - \sigma^*\right). \qquad \text{[Eq. S2]}$$

Eqs. S1,S2 are identical to Eqs. 2,3 of the main text. Here $w_{mid}$ is the midpoint of the bistable driving and represents an unstable intermediate (undifferentiated) fate value. $w_{mid}$ satisfies $0 < w_{mid} < 1$. $\alpha\left(\mathrm{tr}\left(\underline{\underline{\sigma}}\right) - \sigma^*\right)$ is the linear bias from the trace of the in-plane stress onto fate; $\alpha$ is a positive scalar and is the strength of the bias. $\sigma^*$ is a scalar stress value, on which the position of the fate boundary depends particularly strongly. $\tau_w$ is the characteristic timescale on which the fate variable evolves, and $\sqrt{D\tau_w}$ is a short length scale that regularizes spatial variation in $w$.

To introduce the effect of fate on stress in the layer, we assume that cell layer contractility is proportional to the fate variable $w$:

$$P = \tilde{P}w. \qquad \text{[Eq. S3]}$$

$P$ is called the target strain because $\mathrm{tr}\left(\underline{\underline{\sigma}}\right) = 0$ if and only if $P$ equals the trace of the strain. The constant of proportionality between fate $w$ and target strain $P$ is $\tilde{P}$, which is not necessarily spatially uniform (see Eq. 5 of main text).

To close this system of equations, we need to specify how to calculate the trace of the in-plane stress $\underline{\underline{\sigma}}$ if we are given a target strain field $P$. The specific equation for calculating in-plane stress $\underline{\underline{\sigma}}$ depends on the geometry of the cell layer and on the properties of the underlying substrate. We consider four distinct cases:

1. A one-dimensional cell layer on a substrate composed of microposts [21],
2. A contractile cell layer in a stripe geometry (infinite in one direction – *y* – and finite in the other, which is *x*) on a substrate composed of microposts [21,23],
3. An axisymmetric contractile cell layer in a disc geometry on a substrate composed of microposts [23],
4. A one-dimensional cell layer on a substrate of finite thickness [22].

For each case, $h_c$ is the height of the cell layer, and $B_c$ is the cell layer's Young's modulus. In cases 1-3, $k$ is the spring constant of an individual micropost and is typically of order $\mathrm{nN}/\mu\mathrm{m}$, and $N$ is the planar number density of posts. For cases 2 and 3, $\nu$ is the cell layer's Poisson's ratio. In all cases, the stress depends on the target strain $P$ as well as on elements of the strain tensor. The elements of the strain tensor are defined in terms of derivatives of elements of the displacement vector $\vec{u}$:

$$e_{ij} = \frac{1}{2}\left(\frac{\partial u_i}{\partial x_j} + \frac{\partial u_j}{\partial x_i}\right). \quad \text{[Eq. S4]}$$

For case 1, a detailed derivation is given in [21]. The constitutive relation for case 1 is:

$$\sigma(x) = h_c B_c \left(\frac{\partial u}{\partial x} - P\right). \quad \text{[Eq. S5]}$$

where $x$ is the spatial coordinate along the cell layer. To calculate the spatial variation of the in-plane stress, we need to enforce force balance between the cell layer and the substrate:

$$\frac{\partial \sigma}{\partial \mathrm{x}} = kNu(x). \quad \text{[Eq. S6]}$$

Taking the derivative of both sides of Eq. S6 with respect to $x$ and substituting $\frac{\partial u}{\partial x}$ from Eq. S5, we find that:

$$\frac{h_c B_c}{kN}\frac{\partial^2 \sigma}{\partial x^2} = \sigma(x) + h_c B_c P. \quad \text{[Eq. S7]}$$

We note that $\frac{h_c B_c}{kN}$ has units of length squared and represents a mechanical length scale that originates from the interplay of cell layer stiffness and substrate stiffness. This equation is the same as Eq. 1 in the main text.

For case 2, the constitutive relation is:

$$\sigma_{xx} = \frac{h_c B_c}{1+\nu}\left(e_{xx} + \frac{\nu}{1-\nu}\left(e_{xx} + e_{yy}\right)\right) - \frac{h_c B_c}{2(1-\nu)}P. \quad \text{[Eq. S8]}$$

$$\sigma_{yy} = \frac{h_c B_c}{1+\nu}\left(e_{yy} + \frac{\nu}{1-\nu}\left(e_{xx} + e_{yy}\right)\right) - \frac{h_c B_c}{2(1-\nu)}P. \quad \text{[Eq. S9]}$$

$$\sigma_{xy} = \frac{h_c B_c}{1+\nu} e_{xy}. \quad \text{[Eq. S10]}$$

For this contractile stripe layer that is translationally invariant along the *y* direction, $e_{yy} = 0$. Additionally, by symmetry, $e_{xy} = 0$. By the definition of the strain tensor, $e_{xx} = \frac{\partial u_x}{\partial x}$, where $u_x$ is the *x*-displacement of the substrate at the cell-substrate contact and – by symmetry – is the only non-zero component of the displacement vector $\vec{u}$ in this case.

If we simply substitute the values of the strain tensor elements into Eqs. S8-S10, we find

$$\sigma_{xx} = \frac{h_c B_c}{(1-\nu^2)} \frac{\partial u_x}{\partial x} - \frac{h_c B_c}{2(1-\nu)} P. \quad \text{[Eq. S11]}$$

$$\sigma_{yy} = \frac{h_c B_c \nu}{(1-\nu^2)} \frac{\partial u_x}{\partial x} - \frac{h_c B_c}{2(1-\nu)} P. \quad \text{[Eq. S12]}$$

$$\sigma_{xy} = 0. \quad \text{[Eq. S13]}$$

To calculate the spatial variation of the in-plane stress, we need to enforce force balance between the cell layer and the substrate:

$$\text{div} \cdot \underline{\underline{\sigma}} = kN\vec{u} = kNu_x\hat{x}. \quad \text{[Eq. S14]}$$

$k\vec{u}$ is the traction force on an individual post. Since $N$ is the planar number density of posts, $kN\vec{u}$ is the traction force per unit area of the substrate and is typically of order $\frac{\text{nN}}{\mu\text{m}^2}$.

To reduce Eq. S14 into an equation for $\sigma_{xx}$, we first note that $(\text{div} \cdot \underline{\underline{\sigma}}) \cdot \hat{x} = \frac{\partial \sigma_{xx}}{\partial x}$; then, we compute the partial derivative with respect to *x* of both sides:

$$\frac{\partial}{\partial x}\left(\left(\text{div} \cdot \underline{\underline{\sigma}}\right) \cdot \hat{x}\right) = \frac{\partial^2}{\partial x^2}\sigma_{xx} = kN\frac{\partial u_x}{\partial x}. \quad \text{[Eq. S15]}$$

Note that based on Eq. S11, we remove the explicit dependence on $u_x$ in Eq. S15 via:

$$kN\frac{\partial u_x}{\partial x} = kN\frac{(1-\nu^2)}{h_c B_c}\sigma_{xx} + kN\frac{(1+\nu)}{2}P. \quad \text{[Eq. S16]}$$

Based on substitution of Eq. S16 into Eq. S15, Eq. S15 becomes:

$$\frac{\partial^2}{\partial x^2}\sigma_{xx} = kN\frac{(1-\nu^2)}{h_c B_c}\sigma_{xx} + kN\frac{(1+\nu)}{2}P. \quad \text{[Eq. S17]}$$

The factor in front of $\sigma_{xx}$, $kN\frac{(1-\nu^2)}{h_c B_c}$, has units of inverse length squared and is proportional to the ratio of substrate stiffness and cell layer stiffness. Based on this quantity, we define a localization length $l_l$:

$$l_l = \sqrt{\frac{h_c B_c}{kN(1-\nu^2)}}. \quad \text{[Eq. S18]}$$

Thus, Eq. S17 becomes:

$$\frac{\partial^2}{\partial x^2}\sigma_{xx} - \frac{1}{l_l^2}\sigma_{xx} - kN\frac{(1+\nu)}{2}P = 0. \quad \text{[Eq. S19]}$$

We must additionally define the boundary condition: at the colony boundary, $\sigma_{xx}$, the component of the stress normal to the boundary, equals zero. (Thus, $\frac{\partial u_x}{\partial x} - \frac{(1+\nu)}{2}P = 0$.)

Given $\sigma_{xx}$ – calculated from Eq. S19 – and $P$, the trace of the stress is:

$$\text{tr}\left(\underline{\underline{\sigma}}\right) = (1+\nu)\sigma_{xx} - \frac{h_c B_c}{2}P = (1+\nu)\sigma_{xx} - \frac{kNl_l^2(1-\nu^2)}{2}P. \quad \text{[Eq. S20]}$$

This is precisely the quantity that feeds back into the evolution of fate in Eqs. S1, S2.

For case 3, the constitutive relation is:

$$\sigma_{rr} = \frac{h_c B_c}{1+\nu}\left(e_{rr} + \frac{\nu}{1-\nu}(e_{rr} + e_{\theta\theta})\right) - \frac{h_c B_c}{2(1-\nu)} P. \quad \text{[Eq. S21]}$$

$$\sigma_{\theta\theta} = \frac{h_c B_c}{1+\nu}\left(e_{\theta\theta} + \frac{\nu}{1-\nu}(e_{rr} + e_{\theta\theta})\right) - \frac{h_c B_c}{2(1-\nu)} P. \quad \text{[Eq. S22]}$$

$$\sigma_{r\theta} = \frac{h_c B_c}{1+\nu} e_{r\theta}. \quad \text{[Eq. S23]}$$

where $r$ and $\theta$ indicate respectively the radial and polar components of a tensor. For this contractile disc that is axisymmetric, $e_{\theta\theta} = \frac{u_r}{r}$. Additionally, by symmetry, $e_{r\theta} = 0$. By the definition of the strain tensor, $e_{rr} = \frac{\partial u_r}{\partial r}$, where $u_r$ is the $r$-displacement of the substrate at the cell-substrate contact and – by symmetry – is the only non-zero component of the displacement vector $\vec{u}$ in this case.

If we simply substitute the values of the strain tensor elements into Eqs. S21-S23,

$$\sigma_{rr} = \frac{h_c B_c}{(1-\nu^2)}\left(\frac{\partial u_r}{\partial r} + \nu \frac{u_r}{r}\right) - \frac{h_c B_c}{2(1-\nu)} P. \quad \text{[Eq. S24]}$$

$$\sigma_{\theta\theta} = \frac{h_c B_c}{(1-\nu^2)}\left(\nu \frac{\partial u_r}{\partial r} + \frac{u_r}{r}\right) - \frac{h_c B_c}{2(1-\nu)} P. \quad \text{[Eq. S25]}$$

$$\sigma_{r\theta} = 0. \quad \text{[Eq. S26]}$$

We need to enforce force balance between the cell layer and the substrate:

$$\text{div} \cdot \underline{\underline{\sigma}} = kN\vec{u} = kNu_r\hat{r}. \quad \text{[Eq. S27]}$$

where $kN\vec{u}$ is the traction force per unit area of the substrate and is typically of order $\frac{\text{nN}}{\mu\text{m}^2}$. Note that $\left(\text{div} \cdot \underline{\underline{\sigma}}\right) \cdot \hat{r} = \frac{\partial \sigma_{rr}}{\partial r} + \frac{1}{r}(\sigma_{rr} - \sigma_{\theta\theta})$ where $(\sigma_{rr} - \sigma_{\theta\theta}) = \frac{h_c B_c}{1+\nu}\left(\frac{\partial u_r}{\partial r} - \frac{u_r}{r}\right)$ and $\frac{\partial \sigma_{rr}}{\partial r} = \frac{h_c B_c}{(1-\nu^2)}\left(\frac{\partial^2 u_r}{\partial r^2} + \frac{\nu}{r}\frac{\partial u_r}{\partial r} - \frac{\nu}{r^2} u_r\right) - \frac{h_c B_c}{2(1-\nu)}\frac{\partial P}{\partial r}$. Thus, Eq. S27 becomes Eq. S28, further simplified in Eq. S29:

$$\frac{\partial \sigma_{rr}}{\partial r} + \frac{1}{r}(\sigma_{rr} - \sigma_{\theta\theta}) = kNu_r. \quad \text{[Eq. S28]}$$

$$r^2 \frac{\partial^2 u_r}{\partial r^2} + r\frac{\partial u_r}{\partial r} - \left(1 + \frac{r^2}{l_l^2}\right) u_r = \frac{1}{2}(1+\nu) r^2 \frac{\partial P}{\partial r}. \quad \text{[Eq. S29]}$$

The boundary condition for Eq. S29 is that $\sigma_{rr}$ at the boundary equals zero. We note that though the boundary condition is in terms of $\sigma_{rr}$, it is not obvious how to express Eq. S28 as an equation in terms of $\sigma_{rr}$ alone. Thus, we phrase the boundary condition in terms of $u_r$ and $\frac{\partial u_r}{\partial r}$ as:

$$\frac{\partial u_r}{\partial r} + \nu \frac{u_r}{r} = \frac{1+\nu}{2} P. \quad \text{[Eq. S30]}$$

In terms of $u_r$, $\frac{\partial u_r}{\partial r}$, and $P$, the trace of the stress, which linearly biases the fate, is:

$$\mathrm{tr}\left(\underline{\underline{\sigma}}\right) = \frac{h_c B_c}{(1-\nu)}\left(\frac{\partial u_r}{\partial r} + \frac{u_r}{r} - P\right). \quad \text{[Eq. S31]}$$

In Section 5, we derive a more convenient differential equation, which is valuable for gaining analytic insights, for the trace of the in-plane stress $\mathrm{tr}\left(\underline{\underline{\sigma}}\right)$ rather than for $u_r$. Because it is easier to impose boundary conditions in terms of $u_r$, however, in practice we numerically solve Eqs. S29, S30 rather than working in terms of $\mathrm{tr}\left(\underline{\underline{\sigma}}\right)$.

For case 4, the constitutive relation in one dimension is:

$$\sigma(x) = h_c B_c \left(\frac{\partial u}{\partial x} - P\right). \quad \text{[Eq. S32]}$$

To calculate the spatial variation of the in-plane stress, we need to enforce force balance between the cell layer and its substrate; in this case, the divergence of the in-plane stress is not simply proportional to the displacement of the underlying substrate. Unlike for cases 1-3, in which the force on an individual micropost is not communicated to other microposts via the substrate, a force applied to the surface of a continuous substrate can propagate through the substrate.

For the sake of clarity, we reproduce here the derivation of an integro-differential equation for $\sigma(x)$ from [22]. In keeping with [22], we include adhesions, which act as linear springs — between the cell layer and its continuous substrate — with effective strength $Y_a$ (with units of spring constant per area). For cases 1-3, we neglected the existence of cell-substrate adhesions, which have some finite stiffness. For cases 1-3, explicit inclusion of adhesions does not change the mathematical structure of the equations because if the adhesions act as Hookean springs, the adhesion "spring" is in series with the micropost "spring"; thus, adhesions simply change the effective spring constant that the cell layer experiences. For case 4, the explicit inclusion of adhesions allows for both local elasticity, due to the adhesions, and non-local elasticity, due to propagation of stress through the continuous substrate.

Because of the elasticity of the cell-substrate adhesions, the displacement of the substrate, $u^s(x)$, is in general not equal to the displacement of the cell layer, $u(x)$. The force balance between the substrate and the adhesions depends on the imbalance between those two displacements:

$$Y_a[u(x) - u^s(x)] = \frac{\partial \sigma(x)}{\partial x}. \quad \text{[Eq. S33]}$$

Eq. S33 is analogous to the force balance in Eq. S6. The introduction of nonlocal elasticity arises via the expression for $u^s(x)$. As previously mentioned, $u^s(x)$ depends on the forces applied everywhere on the surface of the substrate, where the Green's function $G(x - x')$ propagates the effect of a force applied at $x'$ on the substrate's surface to point $x$:

$$u^s(x) = \int dx' G(x - x') \frac{\partial \sigma(x')}{\partial x'}. \quad \text{[Eq. S34]}$$

By taking the derivative of both sides of Eq. S33 with respect to $x$, we find:

$$Y_a \left[ \frac{\partial}{\partial x} u(x) - \frac{\partial}{\partial x} u^s(x) \right] = \frac{\partial^2 \sigma(x)}{\partial x^2}. \quad \text{[Eq. S35]}$$

By substituting $\frac{\partial}{\partial x} u(x)$ from Eq. S32 and $u^s(x)$ from Eq. S34 into Eq. S35, we find:

$$Y_a \left[ \frac{1}{h_c B_c} \sigma(x) + P(x) - L \frac{\partial}{\partial x} \int_0^L dx' G(x - x') \frac{\partial \sigma(x')}{\partial x'} \right] = \frac{\partial^2 \sigma(x)}{\partial x^2}. \quad \text{[Eq. S36]}$$

where $L$ is the extent of the cell layer in the $x$-direction.

We define $l_a = \sqrt{h_c B_c / Y_a}$, a length scale that arises from the interplay between adhesion stiffness and cell layer stiffness and introduce it into Eq. S36.

$$\left[ \sigma(x) + h_c B_c P(x) - h_c B_c L \frac{\partial}{\partial x} \int_0^L dx' G(x - x') \frac{\partial \sigma(x')}{\partial x'} \right] = l_a{}^2 \frac{\partial^2 \sigma(x)}{\partial x^2}. \quad \text{[Eq. S37]}$$

We discuss how to solve Eq. S37 in Section 4 and specify the approximation for $G(x - x')$ that we employ [22].

Because of the four different cases for the cell-substrate mechanical equation, for the sake of clarity, we note that:

- Case 1 is only used for Eq. 1 of the main text.
- Case 2 is only for the purpose of analytical calculations in Sections 5 and 6.
- Model results in Fig. 2(c) of the main text correspond to case 3 of this section.
- Model results in Fig. 3(e, f) of the main text correspond to case 3 of this section.
- Model results in Fig. 4(a, c) of the main text correspond to case 4 of this section.

In this supplemental text, we also make clear in each section whether we are focusing on a case with only local elasticity or one in which stress can propagate through the substrate.

#### 2. Estimating in-plane stress by integrating over cell-substrate forces

In the main text, we propose that the mechanical stresses in the cell layer could guide the formation of a pattern of neural plate (NP) and neural plate border (NPB) cell fates. In this section and the next, we consider data from cell layers grown on an array of microposts, for which the traction force between the cells and their substrate can be measured by imaging micropost deflection [34,35]. In Section 2, we use these traction forces to determine the magnitude of in-plane stresses within the cell layer *in vitro*; we do this to verify that observed magnitude of in-plane stresses could plausibly affect cell fate and behavior (compared to previous studies) and to inform the construction of mathematical models of this system (in Sections 4-6). In Section 3, we then extend the analysis to extract estimates of material properties of the cell layer (like cell layer contractility).

##### *A. An approximate expression for cell layer stress*

First, we consider a rectangular cell monolayer sitting in the *x-y* plane that is much longer along one axis (*y*-axis) than along the other (*x*-axis). One can estimate the in-plane stress via [51]:

$$\langle \sigma_{xx}(x) \rangle = \frac{1}{L_y} \int_0^x \int_0^{L_y} T_x(x', y') dy' \, dx'. \qquad \text{[Eq. S38]}$$

where $T_x$ is the *x*-component of the traction force per cell area. $L_y$ is the length of the cell layer (parallel to the free boundary) over which one averages. $x = 0$ corresponds to the free boundary of the cell layer (where $\langle \sigma_{xx}(x = 0) \rangle = 0$). $\sigma_{xx}$, the *xx* component of the two-dimensional tensor $\underline{\underline{\sigma}}$, has units of force per length because it is a stress integrated over the thickness $h_c$ of the cell layer in the *z* direction.

In the remaining discussion, we use the formalism developed in [21,23]; we predominantly adopt the notation from [23]. Since we are considering an array of posts that behave like Hookean springs,

$$T_x(x', y') = kNu_x(x', y'). \qquad \text{[Eq. S39]}$$

Here, $u_x$ is the *x*-displacement of the substrate caused by the contraction of the cell layer. $k$ is the effective spring constant of the posts. $N$ is the planar density of posts.

Our actual measurements of traction forces are for cell layers in a disc geometry rather than in this rectangular stripe geometry. To develop an estimate of in-plane stress for a cell layer in a disc geometry analogous to Eq. S38, we first provide the force-balance equation in polar coordinates [21,23] in Eqs. S40-S42:

$$\text{div} \cdot \underline{\underline{\sigma}} = kN\vec{u}. \qquad \text{[Eq. S40]}$$

$$\text{div} \cdot \underline{\underline{\sigma}} = \left( \frac{\partial \sigma_{rr}}{\partial r} + \frac{1}{r}\frac{\partial \sigma_{r\theta}}{\partial \theta} + \frac{1}{r}(\sigma_{rr} - \sigma_{\theta\theta}) \right) \hat{r} + \left( \frac{\partial \sigma_{r\theta}}{\partial r} + \frac{1}{r}\frac{\partial \sigma_{\theta\theta}}{\partial \theta} + \frac{2}{r}(\sigma_{r\theta}) \right) \hat{\theta}. \qquad \text{[Eq. S41]}$$

$$\left( \text{div} \cdot \underline{\underline{\sigma}} \right) \cdot \hat{r} = \frac{\partial \sigma_{rr}}{\partial r} + \frac{1}{r}\frac{\partial \sigma_{r\theta}}{\partial \theta} + \frac{1}{r}(\sigma_{rr} - \sigma_{\theta\theta}) = kNu_r. \qquad \text{[Eq. S42]}$$

We then average over the $\theta$-direction to obtain Eq. S43. We use the notation $\frac{1}{2\pi}\int_0^{2\pi}(\dots)d\theta \equiv \langle\dots\rangle$,

$$\langle\left(\text{div}\cdot\underline{\underline{\sigma}}\right)\cdot\hat{r}\rangle = \langle\frac{\partial\sigma_{rr}}{\partial r} + \frac{1}{r}\frac{\partial\sigma_{r\theta}}{\partial\theta} + \frac{1}{r}(\sigma_{rr} - \sigma_{\theta\theta})\rangle = kN\langle u_r\rangle. \quad \text{[Eq. S43]}$$

Since $\int_0^{2\pi}\left(\frac{\partial\sigma_{r\theta}}{\partial\theta}\right)d\theta = 0$, this equation simplifies to:

$$\langle\left(\text{div}\cdot\underline{\underline{\sigma}}\right)\cdot\hat{r}\rangle = \langle\frac{\partial\sigma_{rr}}{\partial r} + \frac{1}{r}(\sigma_{rr} - \sigma_{\theta\theta})\rangle = kN\langle u_r\rangle. \quad \text{[Eq. S44]}$$

Based on the constitutive relations in Eqs. S21-S22, we find that:

$$\sigma_{rr} - \sigma_{\theta\theta} = kN(1-\nu){l_l}^2(e_{rr} - e_{\theta\theta}) = kN(1-\nu){l_l}^2\left(\frac{\partial u_r}{\partial r} - \frac{1}{r}\frac{\partial u_\theta}{\partial\theta} - \frac{u_r}{r}\right). \quad \text{[Eq. S45]}$$

where $e_{rr}$ and $e_{\theta\theta}$ are elements of the strain tensor. $\nu$ is the Poisson's ratio of the cell layer. As defined in Eq. S18, $l_l$ is the localization length, which depends on the ratio of the cell stiffness to the substrate stiffness [21,23].

Since $\int_0^{2\pi}\left(\frac{\partial u_\theta}{\partial\theta}\right)d\theta = 0$,

$$\langle\frac{1}{r}(\sigma_{rr} - \sigma_{\theta\theta})\rangle = kN(1-\nu){l_l}^2\langle\frac{1}{r}\left(\frac{\partial u_r}{\partial r} - \frac{u_r}{r}\right)\rangle. \quad \text{[Eq. S46]}$$

By plugging Eq. S46 into Eq. S44, we conclude that:

$$\langle\left(\text{div}\cdot\underline{\underline{\sigma}}\right)\cdot\hat{r}\rangle = \langle\frac{\partial\sigma_{rr}}{\partial r}\rangle + \frac{kN(1-\nu){l_l}^2}{r}\langle\left(\frac{\partial u_r}{\partial r} - \frac{u_r}{r}\right)\rangle = kN\langle u_r\rangle. \quad \text{[Eq. S47]}$$

Rearranging terms in Eq. S47, we find Eqs. S48 and S49:

$$\frac{\partial}{\partial r}\langle\sigma_{rr}\rangle = \left(-\frac{kN(1-\nu){l_l}^2}{r}\langle\frac{\partial u_r}{\partial r} - \frac{u_r}{r}\rangle + kN\langle u_r\rangle\right). \quad \text{[Eq. S48]}$$

$$\frac{\partial}{\partial r}\langle\sigma_{rr}\rangle = \left(-\frac{kN(1-\nu){l_l}^2}{r}\left(\frac{\partial\langle u_r\rangle}{\partial r} - \frac{\langle u_r\rangle}{r}\right) + kN\langle u_r\rangle\right). \quad \text{[Eq. S49]}$$

Integrating from the boundary of the cell layer (at $r' = r_0$) to $r' = r < r_0$ returns Eqs. S50-S52:

$$\langle\sigma_{rr}(r)\rangle = kN\left(\xi_1(r) + \xi_2(r)\right). \quad \text{[Eq. S50]}$$

$$\xi_1(r) \equiv \int_{r_0}^{r}\langle u_r(r')\rangle\, dr'. \quad \text{[Eq. S51]}$$

$$\xi_2(r) \equiv -(1-\nu){l_l}^2\int_{r_0}^{r}\frac{1}{r'}\left(\frac{\partial\langle u_r(r')\rangle}{\partial r'} - \frac{\langle u_r(r')\rangle}{r'}\right)dr'. \quad \text{[Eq. S52]}$$

As we explain in the paragraph below, we expect that $\xi_2(r)$ is typically much smaller in magnitude than $\xi_1(r)$, so Eq. S50 reduces to the quasi-1D case (see Eq. S38):

$$\langle\sigma_{rr}(r)\rangle \approx kN\xi_1(r) \equiv kN\int_{r_0}^{r}\frac{1}{2\pi}\int_0^{2\pi}u_r(r',\theta')\, d\theta'\, dr'. \quad \text{[Eq. S53]}$$

*B. Comparison of magnitude of cell layer stresses to those observed in previous studies*

To justify the approximation in Eq. S53, we now perform a back-of-the-envelope calculation to check whether $\xi_2(r) \ll \xi_1(r)$. In our experiments, the maximum value of $|\langle u_r \rangle|$ is of order 0.25 μm. The colony radius is 150 μm. The maximum value of $\left|\langle \frac{\partial u_r}{\partial r} \rangle\right|$ is of order 0.1. $\nu$ is expected to be approximately 0.5, and ${l_l}^2$ is typically of order 100 μm$^2$ [22,23]. Near the colony boundary ($r \approx r_0$), $\frac{(1-\nu){l_l}^2}{r}\left(\frac{\partial u_r}{\partial r} - \frac{u_r}{r}\right)$ is of order 0.01 μm, which is significantly smaller than 0.25 μm, the typical maximum of $|\langle u_r \rangle|$. Therefore, we do expect that $\xi_2(r) \ll \xi_1(r)$ for $r \approx r_0$. To support this back-of-the-envelope calculation, in Fig. S1 we explicitly compute $\frac{\xi_2(r)}{(1-\nu){l_l}^2}$ for each cell colony to check that $\xi_2(r) \ll \xi_1(r)$ for $r \approx r_0$.

Since we expect this approximation (Eq. S53) to break down for $r$ small (or approximately equal to $l_l$), we use Eq. S53 to approximate $\langle \sigma_{rr}(r) \rangle$ only for $r \gtrapprox \frac{r_0}{2}$, where $r_0$ is the colony radius. After concluding from the forty-two colonies in Fig. S1A-C that $\xi_2(r) \ll \xi_1(r)$, we provide estimates of $\langle \sigma_{rr}(r) \rangle$ in Fig. S1D. The magnitude of $\langle \sigma_{rr}(r) \rangle$ near the peak is approximately $2\,\frac{\text{nN}}{\mu\text{m}}$, consistent with estimates of cell layer stress in other contexts [48,51-55]. For example, assuming a cell layer height of 5 μm, Trepat *et al.* estimate that the magnitude of $\langle \sigma_{xx}(x) \rangle$ in an expanding cell colony is on the order of $1\,\frac{\text{nN}}{\mu\text{m}}$ (see Fig. 3 in [51]). In measuring how traction forces scale with the size of cohesive cell colonies, Mertz *et al.* estimate that the active stress (integrated over the cell layer height) is approximately $1\,\frac{\text{nN}}{\mu\text{m}}$ [52].

Though the two previously mentioned studies establish that our measurements of cell layer stresses are comparable to previously published estimates, we must show that the magnitude of cell layer stresses in our experiments could plausibly affect cell behavior and fate choices. In the context of mechanotransduction in collective cell migration, Das *et al.* study a key mechanotransducer called merlin and how its localization in the cell changes in response to cell-cell pulling forces. Das *et al.* find that mechanical stresses of magnitude $1\,\frac{\text{nN}}{\mu\text{m}}$ are sufficient to elicit a mechanical response (see Fig. 3 in [48]); thus, we conclude that our measured values for the magnitude of cell layer stresses are large enough to affect cell behavior by activating mechanosensitive pathways.

#### 3. Estimating cellular material properties via fits to post displacement profiles

In the preceding section, we verified that in our experiments, cell layer stresses are not only of comparable magnitude to cell layer stresses measured in previous studies, but also are large enough to affect cell behavior. In estimating cell layer stresses (Eq. S53), we avoided assuming a specific value for the stiffness of the cell layer, which determines $l_l$ (Eq. S18). In this section, we ask whether we can extract estimates for cellular material properties like contractility from post displacement data.

To extract estimates for the mechanical properties of cell layers, several previous investigators have compared post displacement data to mechanical models of thin contractile sheets. For example, to model traction forces from single cells, one typically treats the single cell as a uniformly contractile medium with a uniform stiffness [24,56]. Even for colonies of multiple cells, previous investigators have assumed that all cells have the same mechanical properties, including both stiffness and contractility [20,25,52]. In our system, the cells in the colony are differentiating in a spatially patterned manner; thus, we expect that a model of a uniform contractile medium will not fit our traction force measurements well.

##### *A. Defining three distinct models for fitting traction force data*

In this section, we fit radial post displacements versus $r$ to three increasingly complicated models of the pattern of cell fates and contractilities, with the goal of determining how many mechanically distinct regions are present in the colonies. To fit the post displacements to some functional form, we assume that the cell layer is axisymmetric. We assume the cell layer is composed of distinct (concentric) domains with distinct cell stiffnesses and distinct contractilities. For each model, we use the formalism developed in [23] to determine the displacements given the spatial pattern of the target strain $P(r)$. As discussed in the supplementary note in [23], we assume that at the boundary between two domains, $u_r$ and $\sigma_{rr}$ are continuous.

Before introducing each of the three models, we would like to point out a few choices that we made to reduce model complexity. The three choices that we make to reduce model complexity are as follows:

1. Instead of fitting post displacements all the way to the nominal colony radius (150 μm), we only fit a functional form to post displacements from $r = 0$ μm to $r = 140$ μm.

2. For models with multiple domains with distinct mechanical properties (models 2 and 3 below), we assume that the innermost domain is very stiff relative to the substrate.
3. We assume a specific value for the Poisson's ratio of the cell layer.

Below we explain why we made those choices and how those choices should affect the fits.

Relative to our first simplifying choice, one might expect that the maximum $|\langle u_r(r)\rangle|$ would occur at the free boundary ($r = 150$ μm) of the micropattern [21,23,24] and that $|\langle u_r(r)\rangle|$ would abruptly collapse to zero for $r > 150$ μm. However, consistent with a recently published study [57], in our experiments the maximal $|\langle u_r(r)\rangle|$ occurs near $r = 140$ μm, and $|\langle u_r(r)\rangle|$ gradually transitions to zero from $r = 140$ μm to $r = 150$ μm (Figs. S2-S3). We could account for this effect in the model by introducing a region near the edge with different material properties, but that would unnecessarily increase the number of fit parameters. Instead, we incorporate the existence of the region from $r = 140$ μm to $r = 150$ μm by imposing a non-zero stress at $r = 140$ μm. We calculate this non-zero stress along $r$ (*i.e.*, $\sigma_{rr}$) at $r = 140$ μm via the method in Section 2A; in the model descriptions below, we denote by $\tilde{\sigma}$ this $\sigma_{rr}$ value at $r = 140$ μm. Thus, the boundary condition for our fit at $r = 140$ μm is *not* a free boundary and enforces force balance, so we expect that this method should have a minimal effect on the quality of our fits.

Relative to our second simplifying choice, we consistently find that the post displacement profiles near the center of the colony are very nearly linear. Such a linear profile is expected to occur when the localization length (Eq. S18) is equal to or greater than the radius of the interior tissue domain (Figs. S2-S3). To incorporate this observation into our fits (for models 2 and 3 below) and to limit the number of free parameters, we first analytically solve the model from [23] with multiple tissue domains and then take the limit in which the localization length of the innermost tissue domain goes to infinity. By taking this limit, we are assuming that the innermost tissue domain is stiff relative to the underlying substrate and that the cell layer's observed strain along $r$ (*i.e.*, $e_{rr}$) is equal to its target strain along $r$ (*i.e.*, $\frac{P}{2}$ as in Eq. S24). This assumption of a large localization length in the interior tissue domain is reflective of the observed data and allows us to focus our attention on mechanical interaction between cells in the vicinity of the fate boundary ($r \gtrapprox \frac{r_0}{2}$).

The last simplifying assumption relates to Poisson's ratio for the cell layer. For a 1D colony (e.g., a cell layer in a stripe geometry [23]), it is impossible to extract independent estimates of the target strain $P$ and Poisson's ratio $\nu$ from the post displacements. These two values appear in the differential equation for stress (Eq. S19) and in its boundary condition as the product $(1+\nu)P$ [21,23]. (Note that the post displacements are proportional to the derivative of the stress in the 1D case as in Eq. S14.) We argued in Section 2 that our colonies are quasi-1D—*i.e.* that the curvature of the domains can be neglected (see Fig. S1)—so we expect that it will be difficult to extract independent estimates of $P$ and $\nu$. As in other studies [24,25,48,53], we choose a value for Poisson's ratio for the cell layer within a biologically relevant range. We choose $\nu = 0.43$ [24]. We expect that tuning $\nu$ will not qualitatively affect the fit and that such tuning would just change all measured target strain values $P$ by the same constant multiplicative factor.

We detail below our three competing models.

Model 1: The cell layer is uniform in both contractility and stiffness. The cell layer extends to $r = r' = 140$ μm where $\langle \sigma_{rr}(r = r') \rangle = \tilde{\sigma}$. The cell layer has localization length $l_l{}'$ and target strain $P'$. By solving Eqs. S29 and S30 and simplifying notation by replacing $u_r$ with $u$,

$$u(r) = A I_1\left(\frac{r}{l_l{}'}\right); 0 \le r \le r'. \quad \text{[Eq. S54]}$$

$$A \equiv \frac{\left(\frac{1+\nu}{2}P' + \frac{1}{kN\left(l_l{}'\right)^2}\tilde{\sigma}\right)}{\left(\frac{I_0\left(\frac{r'}{l_l{}'}\right)}{l_l{}'} + \frac{(\nu-1)I_1\left(\frac{r'}{l_l{}'}\right)}{r'}\right)}. \quad \text{[Eq. S55]}$$

The free parameters are $l_l{}'$ and $P'$.

Model 2: The cell layer is composed of two distinct domains. One domain extends from $r = 0$ to $r = r'$; this domain has localization length $l_l{}'$ ($l_l{}' \to \infty$) and target strain $P'$. The other domain extends from $r = r'$ to $r = r''$; this domain has localization length $l_l{}''$ and target strain $P''$. The cell layer extends to $r = r'' = 140$ μm where $\langle \sigma_{rr}(r = r'') \rangle = \tilde{\sigma}$.

$$u(r) = \begin{cases} \frac{P'r}{2}; 0 \le r \le r' \\ B I_1\left(\frac{r}{l_l{}''}\right) + C K_1\left(\frac{r}{l_l{}''}\right); r' \le r \le r'' \end{cases}. \quad \text{[Eq. S56]}$$

$$B \equiv \frac{\left(kN\, l_l{}'' P' r' r'' K_0\left(\frac{r''}{l_l{}''}\right) + r''\left(kN\left(l_l{}''\right)^2(1+\nu)P'' + 2\tilde{\sigma}\right)K_1\left(\frac{r'}{l_l{}''}\right) - kN\left(l_l{}''\right)^2(-1+\nu)P' r' K_1\left(\frac{r''}{l_l{}''}\right)\right)}{2kN\, l_l{}''\left(\left(r'' I_0\left(\frac{r''}{l_l{}''}\right) + l_l{}''(-1+\nu)I_1\left(\frac{r''}{l_l{}''}\right)\right)K_1\left(\frac{r'}{l_l{}''}\right) + I_1\left(\frac{r'}{l_l{}''}\right)\left(r'' K_0\left(\frac{r''}{l_l{}''}\right) - l_l{}''(-1+\nu)K_1\left(\frac{r''}{l_l{}''}\right)\right)\right)}. \quad \text{[Eq. S57]}$$

$$C \equiv \frac{\left(kN\, l_l'' P' r' r'' I_0\left(\frac{r''}{l_l''}\right) - r''\left(kN(l_l'')^2(1+\nu)P'' + 2\tilde{\sigma}\right) I_1\left(\frac{r'}{l_l''}\right) + kN(l_l'')^2(-1+\nu)P' r' I_1\left(\frac{r''}{l_l''}\right)\right)}{2kN\, l_l''\left(\left(r'' I_0\left(\frac{r''}{l_l''}\right) + l_l''(-1+\nu) I_1\left(\frac{r''}{l_l''}\right)\right) K_1\left(\frac{r'}{l_l''}\right) + I_1\left(\frac{r'}{l_l''}\right)\left(r'' K_0\left(\frac{r''}{l_l''}\right) - l_l''(-1+\nu) K_1\left(\frac{r''}{l_l''}\right)\right)\right)}. \quad \text{[Eq. S58]}$$

The free parameters are $l_l''$, $r'$, $P'$ and $P''$.

Model 3: The cell layer is composed of three distinct domains. One domain extends from $r = 0$ to $r = r'$; this domain has localization length $l_l'$ ($l_l' \to \infty$) and target strain $P'$. Another domain extends from $r = r'$ to $r = r''$; this domain has localization length $l_l''$ and target strain $P''$. The last domain extends from $r = r''$ to $r = r'''$; this domain has localization length $l_l'''$ and target strain $P'''$. The cell layer extends to $r = r''' = 140$ µm where $\langle \sigma_{rr}(r = r''') \rangle = \tilde{\sigma}$.

For the sake of simplicity (to reduce the number of free parameters), we assume $l_l'' = l_l'''$.

$$u(r) = \begin{cases} \frac{P' r}{2}; 0 \le r \le r' \\ B I_1\left(\frac{r}{l_l''}\right) + C K_1\left(\frac{r}{l_l''}\right); r' \le r \le r'' \\ D I_1\left(\frac{r}{l_l''}\right) + E K_1\left(\frac{r}{l_l''}\right); r'' \le r \le r''' \end{cases}. \quad \text{[Eq. S59]}$$

The free parameters are $l_l''$, $r'$, $r''$, $P'$, $P''$ and $P'''$. The analytical expressions for $B, C, D,$ and $E$ in terms of the fit parameters are rather cumbersome. We do not explicitly state them here, but we do provide a MATLAB function, which contains the definitions of $B, C, D,$ and $E$.

(In reviewing models 1-3, note that although these fits inform our choice of model parameters in Sections 4-6 of this supplementary text and in the main text, we use a distinct notation. For example, although $P''$ and $P'''$ in model 3 could be used to compute $\tilde{P}_1$ and $\tilde{P}_2$ in Eq. 5 in the main text, we intentionally use a different notation for these fits to data and our subsequent modeling of fate patterning.)

##### *B. Comparing the goodness-of-fit of three distinct models for traction force data*

Given a specific model, we used the curve fitting tool in MATLAB to find the parameters which correspond to the best fit (cftool in MATLAB 2016B, Mathworks, Natick, MA). For a specific sample – *i.e.*, a single colony – we have three distinct fits, one for each of the three competing models. We would like to compare models 1-3 to see which is most correct based on goodness-of-fit to the data. Because these post displacement data are from early differentiation, we expect significant sample-to-sample variability. Thus, it is possible that one model might perform best for some samples, while another model performs best for other samples. Additionally, because each fit is nonlinear and because there are potentially meaningful parameter differences between samples, we avoid averaging parameters (or goodness-of-fit metrics) over our set of samples. Instead, we report the best-fit parameters for each sample and subsequently compare the models on a sample-to-sample basis.

To compare the models, we would like to know the probability $Pr(M_j|D)$ that model $M_j$ ($j = 1, 2, 3$) is correct for the data $D$. For the sake of clarity, we reproduce some of the discussion from [58,59] below. The probability of the data $D$ given the model $M_j$ is

$$Pr\left(D\middle|M_j\right) = \int Pr\left(D\middle|\vec{\vartheta}_j, M_j\right)Pr\left(\vec{\vartheta}_j\middle|M_j\right)d\vec{\vartheta}_j. \qquad \text{[Eq. S60]}$$

where $Pr\left(D\middle|\vec{\vartheta}_j, M_j\right)$ is the likelihood of $M_j$ with parameters $\vec{\vartheta}_j$. $Pr\left(\vec{\vartheta}_j\middle|M_j\right)$ is the prior probability of parameters $\vec{\vartheta}_j$ for the model $M_j$. One can compare two models (for example, $M_1$ and $M_2$) via:

$$\frac{Pr(M_2|D)}{Pr(M_1|D)} = \left[\frac{Pr(D|M_2)}{Pr(D|M_1)}\right]\left[\frac{Pr(M_2)}{Pr(M_1)}\right] = \text{(ratio of likelihoods of models for data D)(prior ratio)}. \qquad \text{[Eq. S61]}$$

If our prior is that both models are equally likely, then the ratios of the probabilities of the two models $M_1$ and $M_2$, given the data, is simply equal to the ratio $\left[\frac{Pr(D|M_2)}{Pr(D|M_1)}\right]$.

To compute $Pr\left(D\middle|M_j\right)$, we must compute $Pr\left(D\middle|\vec{\vartheta}_j, M_j\right)$. Suppose we have $n$ distinct values of the independent variable $(r_1, r_2, \ldots, r_{n-1}, r_n)$; in this case, this variable is the radial coordinate. For each value of the radial coordinate $r_i$, we have a measured value of the dependent variable $y_i$, in our case the concentrically averaged radial post displacement, and the standard deviation $\sigma_i$ of its random error. (For each radial position $r_i$, we estimate $\sigma_i$ straightforwardly by calculating the standard error of the mean radial post displacement.) For each value $r_i$, we also have a predicted value of the dependent variable $f_i(\vec{\vartheta}_j, M_j)$; the predicted value of the dependent variable depends on the value of fit parameters ($\vec{\vartheta}_j$) and the model $M_j$. The likelihood of observing dependent variables $y_i$, which we assume to normally distributed with standard error $\sigma_i$, given the predicted values $f_i(\vec{\vartheta}_j, M_j)$ from model $M_j$ is:

$$Pr\left(D\middle|\vec{\vartheta}_j, M_j\right) = \prod_{i=1}^{n}\left[\frac{1}{\sqrt{2\pi}\sigma_i}\exp\left(-\frac{1}{2}\left(\frac{y_i - f_i(\vec{\vartheta}_j, M_j)}{\sigma_i}\right)^2\right)\right]. \qquad \text{[Eq. S62]}$$

$$\log\left(Pr\left(D\middle|\vec{\vartheta}_j, M_j\right)\right) = \sum_{i=1}^{n}\left[-\frac{1}{2}\log(2\pi) - \log(\sigma_i) - \frac{1}{2}\frac{1}{(\sigma_i)^2}\left(y_i - f_i\left(\vec{\vartheta}_j, M_j\right)\right)^2\right].$$

[Eq. S63]

To compare the likelihood values of models, we use the Bayesian information criterion (BIC), an approximation to the integral in Eq. S60 [58-60]. The assumptions behind the BIC are described in Section 4 of [58]. For a choice of "noninformative" prior distribution $Pr(\vec{\vartheta}_j|M_j)$ described in [58],

$$\log\left(Pr(D|M_j)\right) = -\frac{1}{2}BIC(M_j) + O\left(n^{-\frac{1}{2}}\right). \qquad \text{[Eq. S64]}$$

$$BIC(M_j) = -2\log\left(Pr\left(D\middle|\hat{\vec{\vartheta}}_j, M_j\right)\right) + d_j\log(n). \qquad \text{[Eq. S65]}$$

where $d_j$ is the number of parameters in the model $M_j$ and $\hat{\vec{\vartheta}}_j$ is the maximum-likelihood estimate of fit parameters $\vec{\vartheta}_j$ for model $M_j$ ($j = 1, 2, 3$). The error $O\left(n^{-\frac{1}{2}}\right)$ is such that $\lim_{n\to\infty} n^{\frac{1}{2}}O\left(n^{-\frac{1}{2}}\right) =$ constant.

To compare two models (for example, $M_1$ and $M_2$), we compute the following ratio:

$$BF(M_2, M_1) = \frac{Pr(M_2|D)}{Pr(M_1|D)} = \exp\left(\frac{log\left(Pr(D|M_2)\right)}{log\left(Pr(D|M_1)\right)}\right) \approx \exp\left(\frac{\Delta BIC_{12}}{2}\right). \qquad \text{[Eq. S66]}$$

where $\Delta BIC_{12} = BIC(M_1) - BIC(M_2)$.

In Tables S1-S3, for each sample, we summarize the best-fit parameters for models 1-3, respectively. These tables include estimates of the localization length (Eq. S18) as well as the target strain, which is proportional to active stress. We conclude by discussing which model performs best (see Table S4). We can categorically reject model 1; based on the BIC, model 1 performs worse than model 2 and model 3 for each of the more than forty samples. Comparing the performance of model 2 and model 3 is more difficult. For some samples, model 2 outperforms model 3 ($\frac{Pr(M_2|D)}{Pr(M_3|D)} > 20$; Blue in Table S4). For some samples, model 3 outperforms model 2 ($\frac{Pr(M_3|D)}{Pr(M_2|D)} > 20$; Maize in Table S4). Those which are neither maize nor blue in Table S4 are those for which $0.05 < \frac{Pr(M_3|D)}{Pr(M_2|D)} < 20$.

Why might it be difficult to distinguish between model 2 and model 3 for these data? As motivated by Fig. 3D,D' and formalized in Eq. 5 of the main text, we attribute the outermost domain in model 3 to the ring of cells (a single-cell wide) at the colony boundary and the intermediate domain in model 3 to a ring of differentiating NPB cells in the colony interior. We expect it might be difficult to distinguish model 2 from model 3 for some samples for the following reasons:

1. Because these traction force measurements are for cell layers at day 4 (two days after initiation of neural induction [11]), the size of the NPB domain might not have reached

steady-state; thus, the intermediate domain (of NPB cells) might be small for some samples, making model 2 and model 3 difficult to distinguish from each other.

2. We have assumed that the fate domains are axisymmetric. If this assumption is broken, we expect that it might be difficult to detect three distinct domains in the concentrically averaged micropost displacements because the two exterior domains would be blended into each other in the average.

Based on these data, which model should we choose? Which ideas should motivate our subsequent mathematical modelling? We choose model 3 for a few reasons. First, we have evidence that the cells at the boundary are morphologically distinct from all other cells in the colony (Fig. 3D,D'). Second, based on previous studies [29,30,38-40], we expect that the cells which lack adherens junctions on one side, because they are at the colony boundary, would have distinct mechanical properties. Also, for many samples, model 3 does, indeed, clearly outperform model 2 (Table S4). In Section 6, we discuss how the existence of a ring of cells at the colony boundary that is more contractile than the cells in the colony interior renders the position of the fate boundary stable to fluctuations in position; thus, in the context of our model, the existence of this ring of cells (a single-cell wide) at the colony boundary is essential for generating an NPB domain that extends into the colony interior.

#### 4. Simulations of tissue patterning on substrate of finite thickness

For the key experimental tests of our fate patterning model (in Fig. 4), the cell layer is bound to a continuous substrate with a thickness that is of the same order as the cell layer's radius (compare 200 μm radius to entries of Table S5). The mechanical model that we detail in Eqs. 1, 4 of the main text [21,23] does not include any effects of substrate thickness. To model the mechanical effect of both the substrate's thickness and the substrate's Young's modulus on the size of the PAX3+ domain (*i.e.*, neural plate border), we employ a formalism developed in a previous study [22]. See case 4 of Section 1 for a quick summary of this formalism.

Unfortunately, the non-local elasticity renders the case of finite-thickness substrates analytically intractable, which prevents us from developing a physical intuition for the predictions of Fig. 4. In Sections 5 and 6, we develop some analytical tools for the case of a cell layer coupled to a substrate of microposts. These analytical tools allow us to develop a physical intuition for the non-monotonic dependence of the NPB domain size on the stiffness of the substrate and for the weak dependence of NPB domain size on colony diameter.

#### A. Technique for evaluating stresses for cell layer bound to finite-thickness substrate

We use the formalism developed in [22] to calculate the mechanical stress in a thin, contractile cell layer bound to a finite-thickness substrate. Specifically, we consider a one-dimensional cell layer (see case 4 of Section 1) of length $L$ ($L = 400$ μm for most of our colonies) lying parallel to the *x* axis and sitting on top of an elastic substrate occupying the region $0 \leq z \leq h_s$ in the *x*-*z* plane. The mechanical stress $\sigma(x)$ within the cell layer has both a passive, elastic contribution and an active contribution: $\sigma(x) = h_c B_c \partial_x u + \sigma_a$, where $u(x)$ is the displacement field of the cellular medium, $h_c$ is the thickness of the cell layer, $B_c$ is an elastic modulus of the cell layer, and $\sigma_a = -h_c B_c P$ is an active, contractile stress (as in Eq. 1 of the main text). Note that $\sigma(x)$ as defined here thus differs by a factor of $h_c$ from the convention used in [22].

We take the substrate to be a linear elastic medium with Poisson's ratio $\nu_s$ and Young's modulus $E_s$; the displacement field of the substrate is $u^s(x)$. In contrast to the case of a cell layer on a micropost array (but in keeping with [22]), here we also allow for focal adhesions of finite stiffness linking the cell layer and the substrate (see case 4 of Section 1 for additional discussion). Force balance on the cell layer then takes the form $Y_a[u(x) - u^s(x)] = \partial_x \sigma(x)$, where $Y_a$ is an effective focal adhesion strength. If we differentiate this equation with respect to *x* and rewrite $u^s(x)$ in terms of the Green's function $G(x)$ giving the response of the substrate to a point force at its surface, we finally arrive at the analog, for a finite thickness substrate, of Eq. 1 of the main text:

$$l_a{}^2 \frac{\partial^2 \sigma}{\partial x^2} - \sigma + \sigma_a = -B_c L h_c \int_0^L dx' \frac{\partial G(|x-x'|)}{\partial x} \frac{\partial \sigma(x')}{\partial x'}. \qquad \text{[Eq. S67]}$$

Here $l_a = \sqrt{h_c B_c / Y_a}$ is a length scale that arises from the interplay between adhesion stiffness and cell layer stiffness; we assume that the adhesions are relatively stiff such that this length scale is approximately 2 μm.

In [22], the authors derive the approximate expression for the Green's function

$$G(x) \approx \frac{2}{\pi L E_s} K_0 \left[\frac{a+|x|}{h_s(1+\nu_s)}\right]. \qquad \text{[Eq. S68]}$$

which we will use throughout this section. Here, $a$ is a short distance cutoff that makes $\lim_{x \to 0} G(x)$ finite. We choose $a \approx 0.5$ μm, the approximate size of an adhesion complex [35,61].

We now describe our numerical methods for solving Eqs. S67, S68. For the sake of completeness, we first reproduce the discussion in [22] on how to solve for in-plane stress $\sigma(x)$ in a cell layer bound to an infinitely thick elastic substrate. For an infinitely thick substrate, the Green's function (Eq. S68) becomes:

$$G(x) = \left(-\frac{2}{\pi L E_s}\right)\left(\gamma + \log\left(\frac{a+|x|}{L}\right)\right). \qquad \text{[Eq. S69]}$$

where $\gamma$ is the Euler-Mascheroni constant. We compute $\frac{\partial G(|x-x'|)}{\partial x}$

$$\frac{\partial G(|x-x'|)}{\partial x} = \left(-\frac{2}{\pi L E_s}\right)\frac{(x-x')}{|x-x'|(a+|x-x'|)}. \qquad \text{[Eq. S70]}$$

As discussed in the supplemental material of [22], we expand $\sigma(x)$ in a Fourier series. Taking note of the boundary conditions $\sigma(x) = 0$ at $x = 0, L$ and $\frac{d\sigma}{dx} = 0$ at $x = \frac{L}{2}$, we define $\sigma_n \equiv \frac{2}{L}\int_0^L dx\left(\sin\left(n\pi\left(\frac{x}{L}\right)\right)\sigma(x)\right)$, and similarly for $\sigma_{a,n}$ and $\sigma_a(x)$. Eq. S67 becomes:

$$\sigma_a(x) = \sum_{n=1,3,5,..} \sigma_n \sin\left(n\pi\left(\frac{x}{L}\right)\right)\left[(n\pi)^2\left(\frac{l_a}{L}\right)^2 + 1\right] -$$

$$B_c L h_c \int_0^L dx' \frac{\partial G(|x-x'|)}{\partial x}\sum_{m=1,3,5,...} \sigma_m \cos\left(m\pi\left(\frac{x'}{L}\right)\right)\left(\frac{m\pi}{L}\right). \qquad \text{[Eq. S71]}$$

Integrating both sides of Eq. S71 by $\frac{2}{L}\int_0^L dx \sin\left(n\pi\left(\frac{x}{L}\right)\right)$,

$$\sigma_{a,n} = \left[(n\pi)^2\left(\frac{l_a}{L}\right)^2 + 1\right]\sigma_n -$$

$$B_c L h_c\left(\frac{2}{L}\right)\sum_{m=1,3,5,...}\sigma_m\left(\frac{m\pi}{L}\right)\int_0^L dx\int_0^L dx' \sin\left(n\pi\left(\frac{x}{L}\right)\right)\frac{\partial G(|x-x'|)}{\partial x}\cos\left(m\pi\left(\frac{x'}{L}\right)\right). \qquad \text{[Eq. S72]}$$

Plugging in Eq. S70,

$$\sigma_{a,n} = \left[(n\pi)^2\left(\frac{l_a}{L}\right)^2 + 1\right]\sigma_n +$$

$$\frac{4B_c h_c}{\pi L E_s}\sum_{m=1,3,5,...}\sigma_m\left(\frac{m\pi}{L}\right)\int_0^L dx\int_0^L dx' \sin\left(n\pi\left(\frac{x}{L}\right)\right)\frac{(x-x')}{|x-x'|(a+|x-x'|)}\cos\left(m\pi\left(\frac{x'}{L}\right)\right). \qquad \text{[Eq. S73]}$$

Defining ${l_{s\infty}}^2 \equiv \frac{4B_c h_c L}{\pi E_s}$,

$$\sigma_{a,n} = \left[(n\pi)^2\left(\frac{l_a}{L}\right)^2 + 1\right]\sigma_n + \left(\frac{l_{s\infty}}{L}\right)^2\sum_{m=1,3,5,...}\sigma_m H_{mn}. \qquad \text{[Eq. S74]}$$

where $H_{mn} = m\pi\int_0^1 dx\int_0^1 dx' \sin(n\pi x)\frac{(x-x')}{|x-x'|(a/L+|x-x'|)}\cos(m\pi x')$. [Eq. S75]

where $n = 1,3,5, ...$

If instead we consider a cell layer bound to an elastic substrate of finite thickness $h_s$,

$$\frac{\partial G(|x-x'|)}{\partial x} = \left(-\frac{2}{\pi L E_s h_s(1+\nu_s)}\right)\frac{(x-x')K_1\left(\frac{a+|x-x'|}{h_s(1+\nu_s)}\right)}{|x-x'|}. \qquad \text{[Eq. S76]}$$

Plugging Eq. S76 into Eq. S72,

$$\sigma_{a,n} = \left[(n\pi)^2 \left(\frac{l_a}{L}\right)^2 + 1\right]\sigma_n -$$

$$B_c L h_c \left(\frac{2}{L}\right)\left(-\frac{2}{\pi L E_s h_s (1+\nu_s)}\right)\sum_{m=1,3,5,\dots} \sigma_m \left(\frac{m\pi}{L}\right)\int_0^L dx \int_0^L dx' \sin\left(n\pi\left(\frac{x}{L}\right)\right)\frac{(x-x')K_1\left(\frac{a+|x-x'|}{h_s(1+\nu_s)}\right)}{|x-x'|}\cos\left(m\pi\left(\frac{x'}{L}\right)\right). \quad \text{[Eq. S77]}$$

$$\sigma_{a,n} = \left[(n\pi)^2 \left(\frac{l_a}{L}\right)^2 + 1\right]\sigma_n + \left(\frac{l_{s\infty}}{L}\right)^2 \sum_{m=1,3,5,\dots} \sigma_m H_{mn}. \quad \text{[Eq. S78]}$$

where $H_{mn} = \left(\frac{1}{(1+\nu_s)}\right)\frac{1}{L h_s} m\pi \int_0^L dx \int_0^L dx' \sin\left(n\pi\left(\frac{x}{L}\right)\right)\frac{(x-x')K_1\left(\frac{a+|x-x'|}{h_s(1+\nu_s)}\right)}{|x-x'|}\cos\left(m\pi\left(\frac{x'}{L}\right)\right).$ [Eq. S79]

For both the cases of infinitely thick substrates and of finitely thick substrates, the linear relationship between the Fourier coefficients can be written in the form:

$$\left(\sigma_{a,1}\ \sigma_{a,3}\ \sigma_{a,5}\ \dots\right) = \left(\sigma_1\ \sigma_3\ \sigma_5\ \dots\right) \times \left[\begin{pmatrix} 1 & 0 & 0 & \\ 0 & 1 & 0 & \\ 0 & 0 & 1 & \\ & & & \ddots \end{pmatrix} + \begin{pmatrix} (\pi)^2 & 0 & 0 & \\ 0 & 9(\pi)^2 & 0 & \\ 0 & 0 & 25(\pi)^2 & \\ & & & \ddots \end{pmatrix}\left(\frac{l_a}{L}\right)^2 + \begin{pmatrix} H_{11} & H_{13} & H_{15} & \\ H_{31} & H_{33} & H_{35} & \\ H_{51} & H_{53} & H_{55} & \\ & & & \ddots \end{pmatrix}\left(\frac{l_{s\infty}}{L}\right)^2\right]. \quad \text{[Eq. S80]}$$

To compute the elements $H_{mn}$, we use the fact that:

$$H_{mn} = 2\left(\frac{1}{(1+\nu_s)}\right)\frac{1}{L h_s} m\pi \int_0^L dx \int_0^x dx' \sin\left(n\pi\left(\frac{x}{L}\right)\right)\frac{(x-x')K_1\left(\frac{a+|x-x'|}{h_s(1+\nu_s)}\right)}{|x-x'|}\cos\left(m\pi\left(\frac{x'}{L}\right)\right). \quad \text{[Eq. S81]}$$

We evaluate $H_{mn}$ via integral2 in MATLAB (MATLAB 2016B, Mathworks, Natick, MA). We solve the linear equation via linsolve in MATLAB (MATLAB 2016B, Mathworks, Natick, MA). We truncate at $n = 400$, such that our discretization is on the order of our short distance cutoff $a \approx 0.5$ µm, the approximate size of an adhesion complex [35,61].

##### *B. Results of simulations of tissue patterning on substrate of finite thickness*

Given the framework detailed above (in Section 4A), we would like to explain how we generate the model predictions detailed in Fig. 4. We couple the fate and the stress as detailed in Eqs. 2, 3, 5 of the main text. Because we are solving for stress in a cell layer bound to a substrate of finite thickness, Eqs. S67, S68 replace Eq. 4 of the main text. For each of the substrate conditions (Table S5), we use the experimental estimates of the Young's modulus and height of the substrate. Table S6 details the remaining parameters for the simulations in Fig. 4. We find, in agreement with experimental results, that the NPB domain size depends weakly on colony diameter and non-monotonically on substrate stiffness.

To verify that the results of simulations in Fig. 4 are representative, we scan over values of both cell layer stiffness $B_c$ and $\sigma^*$. As shown in Fig. S4, the NPB domain size, in general, depends non-monotonically on substrate stiffness and is, in general, approximately independent of colony diameter. Because the equation for mechanical stress (Eq. S67) is analytically intractable, developing a physical intuition for the predictions of Fig. 4 based on Eq. S67 is difficult. To develop an analytically motivated physical intuition for our simulation results Figs. 4, S4, we proceed to consider the limiting case of a cell layer bound to a substrate for which the elasticity is entirely local. That limiting case is the subject of Sections 5 and 6.

#### 5. Differential equation for mechanical pressure in cell layer bound to micropost array

In this section, to arrive at the simpler form given in Eq. 4 of the main text, we rewrite Eqs. S21–S31 describing case 3 of Section 1 in terms of the trace of the stress tensor. Specifically, we consider the case of a cell layer bound to a substrate for which the elasticity is local (either a substrate of microposts or a substrate that is thin relative to the cell layer's in-plane extent) [21,23]. Let us, further, assume that cell layer is two-dimensional. Since we are only considering the lowest-order, rotationally invariant feedback of stress onto fate, we do not need to know the full structure of $\underline{\underline{\sigma}}$; we only need to know the trace of $\underline{\underline{\sigma}}$ (or, equivalently, the mechanical pressure). Therefore, we would like to reduce the full equation for the mechanical stress tensor $\underline{\underline{\sigma}}$ to an equation for the trace of $\underline{\underline{\sigma}}$. The form of the equation that results from this section will be particularly useful for the analytical approximation of NPB domain size developed in Section 6.

We start from the force balance and constitutive equations for case 3 of Section 1, expressed in rotationally invariant form using the Einstein summation convention:

$$\partial_i \sigma_{ij} = kNu_j. \qquad \text{[Eq. S82]}$$

$$\sigma_{ij} = \frac{h_c B_c}{1+\nu}\left(e_{ij} + \frac{\nu}{1-\nu} e_{kk}\delta_{ij} - \frac{1+\nu}{2(1-\nu)} P\delta_{ij}\right). \qquad \text{[Eq. S83]}$$

where $e_{ij} = \frac{1}{2}\left(\partial_i u_j + \partial_j u_i\right)$ is the strain tensor, $u_i$ is the vectorial displacement field, and $\partial_i = \partial/\partial x_i$. We can split both the stress and the strain tensors into their traces and their deviatoric parts:

$$\sigma_{ij} = \frac{\bar{\sigma}}{2}\delta_{ij} + \tilde{\sigma}_{ij}. \qquad \text{[Eq. S84]}$$

$$e_{ij} = \frac{\bar{e}}{2}\delta_{ij} + \tilde{e}_{ij}. \qquad \text{[Eq. S85]}$$

Substituting these expressions into the constitutive relation, Eq. S83, one immediately concludes that

$$\tilde{\sigma}_{ij} = \frac{h_c B_c}{1+\nu} \tilde{e}_{ij}. \qquad \text{[Eq. S86]}$$

$$\bar{\sigma} = \frac{h_c B_c}{1-\nu} (\bar{e} - P). \qquad \text{[Eq. S87]}$$

Taking the divergence of the force balance equation, Eq. S82, and noting that $\partial_i u_i = \bar{e}$, we then have

$$kN\bar{e} = \partial_i \partial_j \sigma_{ij} = \frac{h_c B_c}{1+\nu} \left( \partial_i \partial_j e_{ij} + \frac{\nu}{1-\nu} \partial_i^2 \bar{e} - \frac{1+\nu}{2(1-\nu)} \partial_i^2 P \right). \qquad \text{[Eq. S88]}$$

Further, we can replace $e_{ij}$ by an expression involving derivatives of $u_i$ to arrive at $\partial_i \partial_j e_{ij} = \partial_i^2 \partial_j u_j = \partial_i^2 \bar{e}$. Finally, we rewrite $\bar{e}$ in terms of $\bar{\sigma}$ and $P$ and rearrange to obtain

$$\partial_i^2 \left( \bar{\sigma} + \frac{h_c B_c}{2} P \right) = \frac{1}{l_l^2} \left( \bar{\sigma} + \frac{h_c B_c}{2} P \right) + \frac{kN(1+\nu)^2}{2} P. \qquad \text{[Eq. S89]}$$

With the identifications

$$\sigma_{smooth} = \bar{\sigma} + \frac{h_c B_c}{2} P \qquad \text{[Eq. S90]}$$

$$\bar{\sigma} = \mathrm{tr}(\underline{\underline{\sigma}}) \qquad \text{[Eq. S91]}$$

$$l_l^2 = \frac{h_c B_c}{kN(1-\nu^2)}, \qquad \text{[Eq. S92]}$$

this equation is identical to Eq. 4 of the main text. In the next section, we comment on the physical significance of the quantity $\sigma_{smooth}$.

#### 6. Domain-wall approximation

In Section 5, we derived a differential equation (Eq. S89, equivalent to Eq. 4 of the main text) for the in-plane mechanical pressure, which is the quantity that feeds back onto cell fate in our model. In this section, we use this equation as the starting point to develop analytic expressions for the size of the NPB domain in the limit of a sharp boundary between the NP and NPB domains. These results will give us some physical understanding of the numerical results shown in Fig. 4 of the main text.

##### *A. Position of the fate boundary in the domain-wall limit*

To make analytic progress, we work in the limit that there is a sharp domain wall separating regions of NP ($w \approx 0$) and NPB ($w \approx 1$) fate. Many well-established methods exist to treat pattern-forming systems in this limit [41-43,62,63]. In their simplest formulation, they require a separation of length scales, with a bistable field (in our case, the fate variable $w$) varying on much shorter scales than the other variables (in our case, some stress variable) in the problem. We thus begin by putting our Eq. S89 in the form of a screened diffusion equation with a decay length that is long relative to the length scale of the fate variable.

It is straightforward to verify that the trace of the stress itself does not obey a screened diffusion equation with a single length scale. To see this fact, we return to case 2 of Section 1, which is the case of a contractile stripe which is infinite along one direction (along the $y$ axis). Suppose there is a discontinuous jump in the target strain $P$; for example, such a discontinuous jump would occur if there is a sharp domain wall separating regions of NP ($w \approx 0$) and NPB ($w \approx 1$) fate. In the vicinity of this sharp jump, $\sigma_{xx}$ must be continuous because it is the stress normal to the domain wall. If $\sigma_{xx}$ were not continuous, force balance would be violated. Via Eq. S20, the trace of the stress is then not continuous because it is just a linear sum of a discontinuous term (proportional to $P$) and a continuous term (proportional to $\sigma_{xx}$). This reflects the fact that, unlike $\sigma_{xx}$, $\sigma_{yy}$ can be discontinuous across the domain wall without violating force balance at the wall. More generally, if the scale $\sqrt{D\tau_w}$ on which $w$, and thus $P$, varies is much less than the mechanical length scale $l_l$, then we expect that $\sigma_{xx}$ varies smoothly on the scale $l_l$, whereas $\mathrm{tr}(\underline{\underline{\sigma}})$ also has contributions that vary on the much shorter scale $\sqrt{D\tau_w}$. In order to apply a perturbative approach based on a separation of length scales $\sqrt{D\tau_w} \ll l_l$, we should thus take $w$ and $\sigma_{xx}$ (not $\mathrm{tr}(\underline{\underline{\sigma}})$) as our basic variables. In order to use the same perturbative approach in more than one dimension, we first need to identify the appropriate rotationally invariant generalization of $\sigma_{xx}$.

From Eqs. S89, S90, it is obvious that the rotationally invariant scalar that varies only on the slow scale $l_l$ that we seek is simply $\sigma_{smooth} = \left(\mathrm{tr}(\underline{\underline{\sigma}}) + \frac{h_c B_c}{2} P\right)$, which satisfies a screened diffusion equation with decay length $l_l$. For the sake of completeness, we reproduce Eqs. S1, S2, S89 below since we will use these extensively in our calculations in the domain-wall limit.

$$\tau_w \frac{\partial w}{\partial t} = f(w, \sigma_{smooth}) + D\tau_w \nabla^2 w. \qquad \text{[Eq. S93]}$$

$$f(w, \sigma_{smooth}) \equiv w(w - w_{mid})(1 - w) + \alpha\left(\sigma_{smooth} - \frac{h_c B_c}{2}\tilde{P}w - \sigma^*\right). \qquad \text{[Eq. S94]}$$

$$l_l^2 \nabla^2 \sigma_{smooth} + g(w, \sigma_{smooth}) = 0. \qquad \text{[Eq. S95]}$$

$$g(w, \sigma_{smooth}) \equiv -\sigma_{smooth} - \frac{1+\nu}{1-\nu}\frac{h_c B_c}{2}\tilde{P}w. \qquad \text{[Eq. S96]}$$

$$\sigma_{smooth} \equiv \left(\mathrm{tr}(\underline{\underline{\sigma}}) + \frac{h_c B_c}{2}\tilde{P}w\right). \qquad \text{[Eq. S97]}$$

Having rewritten the governing equations in terms of the fate field $w$ that varies on a scale $\sqrt{D\tau_w}$ and the auxiliary stress field $\sigma_{smooth}$ that varies on the scale $l_l$, we proceed to apply standard methods to the limit $\sqrt{D\tau_w} \ll l_l$ in which sharp domain walls are expected to appear [41-43]. In this limit, away from the domain wall, both fields vary on the scale $l_l$, and the diffusion term $D\tau_w\nabla^2 w$ in Eq. S93 can be dropped to leading order; $w$ is then slaved to $\sigma_{smooth}$ in steady state. On the other hand, within the domain wall, $w$ jumps from one stable state to the other over a distance of order $\sqrt{D\tau_w}$; on this short scale, spatial variation in $\sigma_{smooth}$ can be neglected, so that $\sigma_{smooth}$ takes on the constant value $\sigma_{smooth}^{FB}$ within the domain wall (where "FB" stands for "Fate Boundary").

For a quasi-1D domain wall (*i.e.*, not only $\sqrt{D\tau_w} \ll l_l$, but also $\sqrt{D\tau_w} \ll R$, where $R$ is the radius of curvature of the domain wall) at steady state, Eq. S93 then reduces to

$$0 \approx f(w, \sigma_{smooth}^{FB}) + D\tau_w \frac{d^2 w}{dx^2} \qquad \text{[Eq. S98]}$$

within the domain wall, where $x$ is the coordinate orthogonal to the domain wall. For the domain wall approximation to be consistent, this equation must have a solution that joins $w^{high}$ on one side of the domain wall to $w^{low}$ on the other side, where $w^{high}$ and $w^{low}$ are the maximal and minimal roots of $f(w, \sigma_{smooth}^{FB}) = 0$, which correspond respectively to the NPB and NP fates. On the scale $\sqrt{D\tau_w}$ over which $w$ varies, we can take the edges of the domain wall to extend to infinity, so that we want a solution that approaches $w^{high}$ and $w^{low}$, respectively, as $x \to \pm\infty$. Such a solution turns out to exist only for one particular value of $\sigma_{smooth}^{FB}$ [41-43,62,63]. Indeed, multiplying Eq. S98 by $\frac{dw}{dx}$ and integrating with respect to $x$ from $-\infty$ to $\infty$ we obtain,

$$-\int_{-\infty}^{\infty} f(w, \sigma_{smooth}^{FB}) \frac{dw}{dx} dx = D\tau_w \int_{-\infty}^{\infty} \frac{d^2 w}{dx^2}\frac{dw}{dx} dx. \qquad \text{[Eq. S99]}$$

Via integration by parts and with no-flux boundary conditions, one can show that the right-hand side of Eq. S99 is equal to zero. We then change the variable for the integral on the left-hand side from $x$ to $w$, obtaining

$$\int_{w^{low}}^{w^{high}} f(w, \sigma_{smooth}^{FB}) dw = 0. \qquad \text{[Eq. S100]}$$

Given a specific set of model parameter values, Eq. S100 determines a unique value for $\sigma_{smooth}^{FB}$ that the smooth stress must take at the fate boundary.

The process for determining the position of the fate boundary for a given set of parameters is hence as follows:

1. Use Eq. S100 to find the value of $\sigma_{smooth}^{FB}$ at which a stationary boundary can exist; this value, which we call $\overline{\sigma_{smooth}^{FB}}$, will depend on the model parameter values but not on the colony size or on position within the colony. Note in particular that $\overline{\sigma_{smooth}^{FB}}$ varies linearly with $\sigma^*$ so that the parameter $\sigma^*$ can be used to vary the position of the fate boundary.
2. After solving for $\overline{\sigma_{smooth}^{FB}}$, calculate the corresponding fate boundary position. One can do this by first calculating the smooth stress at the fate boundary as a function of fate boundary position and then inverting that function to find the boundary position for which $\sigma_{smooth}^{FB} = \overline{\sigma_{smooth}^{FB}}$.

To calculate the smooth stress at the fate boundary as a function of fate boundary position, we make the additional, common approximation that, away from the fate boundary, $w$ equals either 0 (NP) or 1 (NPB) [41,43,62,63]. Then, for each possible position of the fate boundary, we determine the corresponding profile of the target strain, including the effect of the boundary cell and recalling that the fate variable determines the target strain through $P = \tilde{P}w$ (Fig S5). We then solve div $\cdot \underline{\underline{\sigma}} = kN\vec{u}$ (as described in Section 1 and Materials and methods) to find $\sigma(x)$, assuming that that $\vec{u} \cdot \hat{n}$ and $\underline{\underline{\sigma}} \cdot \hat{n}$ are continuous at the fate boundary, as discussed in the supplemental material in [23], where $\hat{n}$ is the unit vector normal to the fate boundary. Finally, we set *x* equal to the fate boundary position to obtain $\sigma_{smooth}^{FB}$.

Below we report the smooth stress at the fate boundary as a function of fate boundary position for a stripe geometry and a disc geometry. By "stripe geometry", we mean case 2 of Section 1, *i.e.* a cell layer that is infinite in one direction (the *y* direction) and finite in the other direction (the *x* direction). In this configuration, the strain along the *y*-axis is zero.

For the stripe geometry, the smooth stress at the fate boundary $\sigma_{smooth}^{FB}$ as a function of fate boundary position $x_1$ is

$$\frac{\sigma_{smooth}^{FB}(x_1)}{\frac{h_C B_C}{1-\nu^2}} = \left\{-\frac{1}{2}(1+\nu)^2\left(-\tilde{P}_2 + \left(-\tilde{P}_1 + \tilde{P}_2\right) * \cosh\left(\frac{x_2 - x_3}{l_l}\right)\right) * \frac{\cosh\left(\frac{x_1}{l_l}\right)}{\cosh\left(\frac{x_3}{l_l}\right)}\right\} + \left\{-\frac{1}{2}(1+\nu)^2\left(\tilde{P}_1 * \cosh\left(\frac{x_1 - x_3}{l_l}\right)\right) * \frac{\cosh\left(\frac{x_1}{l_l}\right)}{\cosh\left(\frac{x_3}{l_l}\right)}\right\}. \quad \text{[Eq. S101]}$$

where for future reference we have written the righthand side as a sum of two terms in curly braces. Here $x_3$ is the distance from the colony center (which we place at $x = 0$) to the colony boundary, and $x_2 = x_3 - \Delta x$ is the distance from the colony center to the boundary cell, where $\Delta x \approx 25\ \mu$m is the width of the boundary cell. (The quantities $x_3$ and $x_2$ correspond respectively to $r_0$ and $r_0 - \Delta r$ in Eq. 5 of the main text for a radially symmetric domain.) Also, note that we have non-dimensionalized the stress on the lefthand side of Eq. S101. The black curve in Fig. S6A is a representative example of Eq. S101.

Far from the colony center and far from the boundary cell (where "far" is relative to the localization length $l_l$), the smooth stress at the fate boundary does not depend on the position of the fate boundary. Near the colony center, the smooth stress at the fate boundary does depend on the fate boundary position; the second term on the righthand side of Eq. S101 largely determines this dependence (see red curve in Fig. S6B). Likewise, near the boundary cell, the smooth stress at the fate boundary depends on the fate boundary position; that dependence is set primarily by the first term on the right-hand side of Eq. S101 (see blue curve in Fig. S6B).

Having calculated $\sigma_{smooth}^{FB}(x_1)$, we can find the fate boundary position by solving $\sigma_{smooth}^{FB}(x_1) = \overline{\sigma_{smooth}^{FB}}$ for $x_1$. Fig. S6A illustrates a graphical solution of this equation, with the dashed horizontal line marking $\overline{\sigma_{smooth}^{FB}}$. We show in Section 6B below that a fate pattern with NP at the center and NPB at the colony edge is stable to shifts in fate boundary position when $\frac{\partial \sigma_{smooth}^{FB}(x_1)}{\partial x_1} > 0$ an unstable otherwise. (We emphasize that this is a condition on $\frac{\partial \sigma_{smooth}^{FB}(x_1)}{\partial x_1}$, that is on the change in the stress at the fate boundary as the fate boundary is moved; this is different from the change $\frac{\partial \sigma_{smooth}}{\partial x}$ in the stress as a function of the coordinate $x$ in the colony at fixed fate boundary position.) Thus, in Fig. S6A, only the outermost intersection corresponds to a stable fate pattern. The NPB domain for the example of Fig. S6A would extend from the colony boundary (at $x_3 = 200$ µm) to the intersection of the gray line and the black curve (at $x_1 \approx 160$ µm, marked by the arrow).

Now that we have treated the case of a cell layer in a stripe geometry, we repeat the analysis for a cell layer in a disc geometry (case 3 of Section 1). For the disc geometry, the smooth stress at the fate boundary $\sigma_{smooth}^{FB}$ as a function of fate boundary position $r_1$ is:

$$\frac{\sigma_{smooth}^{FB}(r_1)}{\frac{h_c B_c}{1-\nu^2}} =$$

$$\left\{\frac{\left((1+\nu)^2\left(l_l*\widetilde{P_2}*r_3-\left(r_3 I_0\left(\frac{r_3}{l_l}\right)+l_l(-1+\nu)*I_1\left(\frac{r_3}{l_l}\right)\right)*\left((-\widetilde{P_1}+\widetilde{P_2})*r_2*K_1\left(\frac{r_2}{l_l}\right)\right)+(\widetilde{P_1}-\widetilde{P_2})*r_2*I_1\left(\frac{r_2}{l_l}\right)*\left(r_3 K_0\left(\frac{r_3}{l_l}\right)-l_l*(-1+\nu)*K_1\left(\frac{r_3}{l_l}\right)\right)\right)\right)}{2\left(r_3 I_0\left(\frac{r_3}{l_l}\right)+l_l*(-1+\nu)I_1\left(\frac{r_3}{l_l}\right)\right)} I_0\left(\frac{r_1}{l_l}\right)\frac{1}{l_l}\right\} +$$

$$\left\{\frac{\left((1+\nu)^2\left(-\left(r_3 I_0\left(\frac{r_3}{l_l}\right)+l_l*(-1+\nu)I_1\left(\frac{r_3}{l_l}\right)\right)*\left(\widetilde{P_1}*r_1*K_1\left(\frac{r_1}{l_l}\right)\right)+\widetilde{P_1}r_1 I_1\left(\frac{r_1}{l_l}\right)*\left(-r_3*K_0\left(\frac{r_3}{l_l}\right)+l_l*(-1+\nu)K_1\left(\frac{r_3}{l_l}\right)\right)\right)\right)}{2\left(r_3 I_0\left(\frac{r_3}{l_l}\right)+l_l(-1+\nu)I_1\left(\frac{r_3}{l_l}\right)\right)} * I_0\left(\frac{r_1}{l_l}\right)\frac{1}{l_l}\right\}.$$

[Eq. S102]

where we have written the righthand side as a sum of two terms in curly braces for future reference. Here $r_3$ is the distance from the colony center to the colony boundary, $r_2 = r_3 - \Delta r$ is the distance from the colony center to the boundary cell, and $\Delta r \approx 25\ \mu$m is the width of the boundary cell. (The colony radius $r_3$ in this equation corresponds to $r_0$ in Eq. 5 of the main text.) The black curve in Fig. S6C is a representative example of Eq. S102. The first term on the righthand side of Eq. S102 depends on $r_2$ and represents the effect of the highly contractile boundary cell. The second term on the right-hand side of Eq. S102 is independent of $r_2$ and dominates the $r_1$ dependence of the fate-boundary stress when the fate boundary is near (in terms of $l_l$) the colony center. The blue and red curves in Fig. S6D are examples of the first and second curly-braced terms, respectively, in Eq. S102.

Fig. S6C illustrates a graphical solution of the equation $\sigma_{smooth}^{FB}(r_1) = \overline{\sigma_{smooth}^{FB}}$ to find $r_1$ at the fate boundary. Just as in the stripe geometry, for the stability of a fate pattern with NP at center and NPB at the colony edge, $\frac{\partial \sigma_{smooth}^{FB}(r_1)}{\partial r_1}$ must be greater than zero at the intersection point (see Section 6B below and [43] for physical argument). In Fig. S6C, only the outermost intersection corresponds to a stable fate pattern. The NPB domain for the example of Fig. S6C would extend from the colony boundary (at $r_3 = 200$ µm) to the intersection of the gray line and the black curve (at $r_1 \approx 160$ µm).

Though the two cases are not qualitatively different, the equation for a cell layer in a stripe geometry is much simpler than that for a cell layer in a disc geometry (compare Eq. S101 to Eq. S102). We thus focus primarily on the stripe case for the remainder of this section. In Section 6B, we briefly provide a physical justification for the stability of a fate boundary to shifts in fate boundary position and a physical confirmation that only one of the intersections in Fig. S6A corresponds to a stable fate pattern. In Section 6C, we use Eq. S101 to demonstrate that NPB domain size is approximately independent of colony diameter. Finally, in Section 6D, we explain analytically why the NPB domain size depends non-monotonically on the substrate stiffness.

##### *B. Stability of fate boundary to shifts in fate boundary position*

We have seen that, within the domain wall approximation, one can determine the position of the fate boundary by finding the position with the appropriate smooth stress at the fate boundary (see Fig. S6A, C). It remains, however, to determine which of the solutions of $\sigma_{smooth}^{FB}(x_1) = \overline{\sigma_{smooth}^{FB}}$ is stable. In this section, we sketch a simple, physically-motivated argument suggesting that one of the two solutions is generically unstable. The essential point is that when $\sigma_{smooth}^{FB}(x_1) \neq \overline{\sigma_{smooth}^{FB}}$, the domain wall separating the two fate domains tends to move in a direction set by the sign of $\sigma_{smooth}^{FB}(x_1) - \overline{\sigma_{smooth}^{FB}}$; a fixed point can thus be expected to be stable or unstable depending on the slope of $\sigma_{smooth}^{FB}(x_1)$ at the fixed point.

In some textbook examples of domain patterns, for example phase-field models, it is possible to reduce the system to an effective interface dynamics and thus to systematically study pattern stability in the limit $\varepsilon \equiv \frac{\sqrt{D\tau_w}}{l_l} \ll 1$ [43]. We make no claims to a similar level of rigor in the present treatment. Indeed, in those standard cases the equivalent of our fate variable $w$ not only varies on a short length scale of order $\varepsilon$, but also relaxes much faster than the variable corresponding to our $\sigma_{smooth}$. In contrast, in our model, $\sigma_{smooth}$ is instead assumed to relax infinitely fast compared to $w$; as a result, the general problem of the linear stability of patterns cannot be reduced to a question of domain wall motion. The analytic results of this section thus do not directly address whether our patterns are stable against arbitrary perturbations. They do, however show that some fixed points are *un*stable against a perturbation corresponding to a small shift of the fate boundary position. This is sufficient to let us rule out one of the two fixed points seen in Fig. S6A, C; our numerical results (Fig. 3(e)-(f) ) can then be taken to confirm the stability of the other fixed point.

For simplicity, we give the argument only for the stripe geometry. Suppose that the position of the fate boundary is displaced a small distance $\delta x_1$ from a solution of $\sigma_{smooth}^{FB}(x_1) = \overline{\sigma_{smooth}^{FB}}$. We focus on the inner region of width of order $\varepsilon$ centered on the fate boundary over which $w$ changes roughly from 0 to 1, and we assume that for small enough $\varepsilon$ the stress is constant to a good approximation across this region. We further assume (as in the calculation of $\sigma_{smooth}^{FB}(x_1)$ above) that outside of this region the fate variable only takes on the values $w \approx 0$ or $w \approx 1$; we are thus only considering stability against perturbations that correspond to shifts in the boundary, not to perturbations that vary $w$ away from the domain wall separating the two fates. With these assumptions, the stress at the fate boundary is $\sigma_{smooth}^{FB} \approx \overline{\sigma_{smooth}^{FB}} + \delta\sigma$, where $\delta\sigma = \left(\frac{\partial \sigma_{smooth}^{FB}(x_1)}{\partial x_1}\right)\delta x_1$. Finally, we assume that within the inner region, $w$ has relaxed to the spatial profile corresponding to a front translating with (constant) velocity $v$; for our ordering of timescales, this assumption cannot be justified *a priori* and again amounts to limiting our analysis to perturbations of a particular shape. Then, it is a standard result [43] that, to leading order in $\delta x$, the domain wall has velocity

$$v = -\delta\sigma \frac{\int_{-\infty}^{\infty} d\xi \left(\frac{dw}{d\xi}\right) \frac{\partial f(w,\sigma_{smooth})}{\partial \sigma_{smooth}}}{\int_{-\infty}^{\infty} d\xi \left(\frac{dw}{d\xi}\right)^2}, \qquad \text{[Eq. S103]}$$

where $\xi = x/\varepsilon$ is a stretched spatial coordinate corresponding to the inner length scale, and the partial derivative of $f$ with respect to $\sigma_{smooth}$ is to be evaluated at $\sigma_{smooth} = \overline{\sigma_{smooth}^{FB}}$.

Depending on whether $v$ has the same sign as $\delta x$ or the opposite sign, the fate boundary location will be either unstable or stable. To determine the sign of $v$, we need to know the signs of $\frac{dw}{d\xi}$, $\frac{\partial f(w,\sigma_{smooth})}{\partial \sigma_{smooth}}$ evaluated at $\sigma_{smooth} = \overline{\sigma_{smooth}^{FB}}$, and $\delta\sigma$.

It is straightforward to check that:

1. For the patterns of interest to us, with the NPB domain at the outside of the colony, $\frac{dw}{d\xi} > 0$.
2. For $\alpha > 0$, as in our system, $\frac{\partial f(w,\sigma_{smooth})}{\partial \sigma_{smooth}} > 0$.

Thus, both integrals in Eq. S103 are positive, and $v$ and $\delta\sigma$ have opposite signs. Finally, because $\delta\sigma = \left(\frac{\partial \sigma_{smooth}^{FB}(x_1)}{\partial x_1}\right)\delta x_1$, we can conclude that when $\frac{\partial \sigma_{smooth}^{FB}(x_1)}{\partial x_1} > 0$ at the fate boundary, $v$ and $\delta x_1$ also have opposite signs, and the fate boundary position is stable (at least against the specific perturbations considered here). On the other hand, when $\frac{\partial \sigma_{smooth}^{FB}(x_1)}{\partial x_1} < 0$, $v$ and $\delta x_1$ have the same sign, and the domain pattern is unstable.

The fact that $\frac{\partial \sigma_{smooth}{}^{FB}(x_1)}{\partial x_1}$ must be greater than zero at the intersection point in order for the fate pattern to be stable also explains why the presence of a mechanically distinct boundary cell is necessary. The heightened contractility of the boundary cell relative to cells in the colony interior (Fig. S5B) allows for $\frac{\partial \sigma_{smooth}{}^{FB}(x_1)}{\partial x_1}$ to be positive near $x_1 \approx x_2$. One can easily see this if one sets $\widetilde{P_1}$ equal to $\widetilde{P_2}$ in Eq. S101, which becomes

$$\frac{\sigma_{smooth}^{FB}(x_1)}{\frac{h_c B_c}{1-\nu^2}} = \left\{-\frac{1}{2}(1+\nu)^2\left(-\tilde{P}_1\right) * \frac{\cosh\left(\frac{x_1}{l_l}\right)}{\cosh\left(\frac{x_3}{l_l}\right)}\right\} + \left\{-\frac{1}{2}(1+\nu)^2 \left(\tilde{P}_1 * \cosh\left(\frac{x_1-x_3}{l_l}\right)\right) * \frac{\cosh\left(\frac{x_1}{l_l}\right)}{\cosh\left(\frac{x_3}{l_l}\right)}\right\}. \quad \text{[Eq. S104]}$$

To show that $\frac{\partial \sigma_{smooth}{}^{FB}(x_1)}{\partial x_1}$ is negative everywhere if $\widetilde{P_1}$ is equal to $\widetilde{P_2}$, we take the derivative with respect to $x_1$ of Eq. S104.

$$\frac{1}{\frac{h_c B_c}{1-\nu^2}} \frac{\partial \sigma_{smooth}^{FB}(x_1)}{\partial x_1} = (1+\nu)^2 \widetilde{P_1} * \frac{\operatorname{sech}\left(\frac{x_3}{l_l}\right)\left(\sinh\left(\frac{x_1}{l_l}\right) - \sinh\left(\frac{2x_1-x_3}{l_l}\right)\right)}{2l_l}. \quad \text{[Eq. S105]}$$

To see that $\frac{\partial \sigma_{smooth}{}^{FB}(x_1)}{\partial x_1}$ is negative everywhere if $\widetilde{P_1}$ is equal to $\widetilde{P_2}$, it is important to note that:

(1) $\widetilde{P_1} < 0$.

(2) $\operatorname{sech}\left(\frac{x_3}{l_l}\right)$ is positive.

(3) $\sinh(x)$ is strictly monotone.

(4) $\frac{x_1}{l_l} \geq \frac{2x_1-x_3}{l_l}$.

(5) Points 3 and 4 imply that $\sinh\left(\frac{x_1}{l_l}\right) - \sinh\left(\frac{2x_1-x_3}{l_l}\right) \geq 0$.

Thus, $\frac{\partial \sigma_{smooth}{}^{FB}(x_1)}{\partial x_1}$ is negative everywhere (except for at $x_1 = x_3$).

If on the other hand $\alpha$ in Eq. S94 (see Fig. 3B) were of opposite sign, then the presence of a mechanically distinct boundary cell would not be necessary to stabilize the pattern. Our sign of the coupling from stress to fate means that a stable fate pattern with a domain wall cannot be formed in the absence of mechanical heterogeneity at the boundary.

C. *NPB domain size is approximately independent of colony diameter, but is slightly larger for small colonies.*

One can rather easily demonstrate that the NPB domain size is approximately independent of colony size. Suppose that the colony diameter is much greater than $l_l$. Then, the position of the fate position is primarily determined by the spatial dependence due to the blue curve (Fig. S6B). The blue profile (see first curly-braced term of Eq. S101) does not depend on colony size; it only depends on the size of the boundary cell $(x_2 - x_3)$ and the position of the fate boundary relative to the boundary $(x_1 - x_3)$ since:

$$\lim_{x_3/l_l \to \infty} \frac{\cosh\left(\frac{x_1}{l_l}\right)}{\cosh\left(\frac{x_3}{l_l}\right)} = \lim_{x_3/l_l \to \infty} \frac{\cosh\left(\frac{(x_1 - x_3) + x_3}{l_l}\right)}{\cosh\left(\frac{x_3}{l_l}\right)} = \exp\left(\frac{(x_1 - x_3)}{l_l}\right). \qquad \text{[Eq. S106]}$$

Based on this logic, if the colony diameter is much greater than $l_l$, the NPB domain size is independent of domain size.

What happens if the colony diameter and $l_l$ are of comparable magnitude or if the colony diameter is small relative to $l_l$? In that case, the tails of the two exponential curves (Fig. S6B) overlap. As the two curves overlap more and more, the size of the NPB domain (from the colony boundary to the intersection of the gray line and the black curve) gets larger. That is, the intersection point moves inward relative to the colony boundary as the colony diameter goes down. If the tails overlap sufficiently, the black curve in Fig. S6A (or in Fig. S6C) may no longer intersect the horizontal line (corresponding to the ${\sigma_{smooth}}^{FB} = \overline{\sigma_{smooth}^{FB}}$); in that case, no stable fate pattern with a domain wall exists because a saddle-node bifurcation has occurred.

This bifurcation hints at a notable feature of this model: $\overline{\sigma_{smooth}^{FB}}$ (which is model-parameter dependent; see Eq. S100) must fall within a certain range in order for a stable fate pattern with a domain wall to exist. For example, in Fig. S6A, the horizontal gray dashed line must be greater than the minimum of the black curve and must be less than the value of the black curve at $x_1 = x_2$. (Physically, this means that the cell at the very boundary is generating enough stress to stabilize some fate boundary away from itself. The outer cell's influence propagates inward toward the colony center.) For the size of the NPB (outer) fate domain to be as large as $5l_l$ (for example), the horizontal gray dashed line must be approximately equal to, but greater than the minimum of the black curve.

This requirement that $\overline{\sigma_{smooth}^{FB}}$ be constrained to a certain range could be problematic; in general, models of living systems should not require fine-tuning of parameters. However, we know that for our *in vitro* experimental model system (as discussed in [11]), the NP/NPB fate pattern exists specifically because the level of BMP inhibition in the spatially uniform medium has been tuned (see Fig. 5 of [11]). We, thus, speculate that the constraint on $\overline{\sigma_{smooth}^{FB}}$ within the model might reflect the tuning of experimental conditions (specifically BMP inhibition) *in vitro*. This experimental fine-tuning allowed us to study the role of mechanical interactions between cells in NP/NPB pattern formation in a simplified *in vitro* system where the cells experience a uniform chemical medium. In an actual developing embryo, we would expect the existence of a non-uniform gradients of chemical morphogens like BMP (see introduction of main text) to interact with mechanical stresses to confer an appropriate robustness of the NP/NPB fate pattern against variations in parameter values.

##### *D. NPB domain size depends non-monotonically on substrate stiffness*

To demonstrate that the NPB domain size depends non-monotonically on the substrate stiffness, we consider the same limit as in Section 6C: that the colony diameter is much greater than $l_l$. Then, the position of the fate boundary is primarily determined by the spatial dependence of the blue curve (Fig. S6B and S6D). Physically, this means that the stress generated by the outermost contractile band is providing the information to position the domain wall. Based on Eqs. S101 and S106, the stress at the fate boundary, in this limit, is

$$\frac{\sigma_{smooth}^{FB}(x_1)}{\frac{h_C B_C}{1-\nu^2}} + \frac{1}{2}(1+\nu)^2\left(\tilde{P}_1 * \frac{1}{2}\right) \approx -\frac{1}{2}(1+\nu)^2\left(-\tilde{P}_2 + \left(-\tilde{P}_1 + \tilde{P}_2\right) * \cosh\left(\frac{x_2 - x_3}{l_l}\right)\right) * \exp\left(\frac{(x_1 - x_3)}{l_l}\right). \quad \text{[Eq. S107]}$$

We would like to know how $\frac{\sigma_{smooth}^{FB}(x_1)}{\frac{h_C B_C}{1-\nu^2}}$ behaves as the stiffness of the substrate changes. For decreasing substrate stiffness $k$, $l_l$ increases (*i.e.* $l_l$ is monotonic in $k$; Eq. S18). Since $l_l$ is monotonic in $k$, all we need to know is how $\frac{\sigma_{smooth}^{FB}(x_1)}{\frac{h_C B_C}{1-\nu^2}}$ changes as $l_l$ varies. We will break down our analysis of $\frac{\sigma_{smooth}^{FB}(x_1)}{\frac{h_C B_C}{1-\nu^2}}$ into how $\exp\left(\frac{(x_1 - x_3)}{l_l}\right)$ changes with $l_l$ as well as how $\left(-\tilde{P}_2 + \left(-\tilde{P}_1 + \tilde{P}_2\right) * \cosh\left(\frac{x_2 - x_3}{l_l}\right)\right)$ changes with $l_l$. By analyzing how the exponential itself as well as its prefactor changes with $l_l$, we hope to gain a physical intuition for the non-monotonic dependence of NPB domain width on substrate stiffness.

The partial derivative of the prefactor of the exponential in Eq. S107 is

$$\frac{\partial}{\partial l_l}\left[-\frac{1}{2}(1+\nu)^2\left(-\tilde{P}_2+\left(-\tilde{P}_1+\tilde{P}_2\right)*\cosh\left(\frac{x_2-x_3}{l_l}\right)\right)\right]=-\frac{1}{2}(1+\nu)^2\left(\left(-\tilde{P}_1+\tilde{P}_2\right)*\frac{\partial}{\partial l_l}\cosh\left(\frac{x_2-x_3}{l_l}\right)\right).$$

[Eq. S108]

$$\frac{\partial}{\partial l_l}\left[-\frac{1}{2}(1+\nu)^2\left(-\tilde{P}_2+\left(-\tilde{P}_1+\tilde{P}_2\right)*\cosh\left(\frac{x_2-x_3}{l_l}\right)\right)\right]=(1+\nu)^2*\left(-\tilde{P}_1+\tilde{P}_2\right)*\left(\frac{x_2-x_3}{2{l_l}^2}\right)*\sinh\left(\frac{x_2-x_3}{l_l}\right).$$

[Eq. S109]

We note that:

(1) $\left(-\tilde{P}_1+\tilde{P}_2\right)<0.$

(2) $(x_2-x_3)*\sinh\left(\frac{x_2-x_3}{l_l}\right)>0.$

Thus, the prefactor of the exponential in Eq. S107 monotonically decreases as $l_l$ increases.

The way that the exponential itself behaves as $l_l$ changes is more obvious.

$$\frac{\partial}{\partial l_l}\left[\exp\left(\frac{-(x_3-x_1)}{l_l}\right)\right]=\frac{(x_3-x_1)}{{l_l}^2}\exp\left(\frac{-(x_3-x_1)}{l_l}\right). \quad \text{[Eq. S110]}$$

As $l_l$ increases, the exponential itself increases. Thus, as discussed in the main text, the stress at the fate boundary (which results from the outermost contractile cell) is an exponential function with a prefactor which increases with increasing substrate stiffness, while the decay length of the exponential itself decreases with increasing substrate stiffness. So if you cut Eq. S101 at some fixed ${\sigma_{smooth}}^{FB}=\overline{\sigma_{smooth}^{FB}}$ as in Fig. S6A and S6C, the competition between these two effects gives rise to the observed non-monotonicity of the NPB domain size with respect to substrate stiffness.

To formalize this argument slightly more, suppose that $\sigma_{smooth}^{FB}=\overline{\sigma_{smooth}^{FB}}$. Then, it follows that

$$\frac{\frac{\overline{\sigma_{smooth}^{FB}}}{\frac{h_C B_C}{1-\nu^2}}+\frac{1}{2}(1+\nu)^2\left(\tilde{P}_1*\frac{1}{2}\right)}{-\frac{1}{2}(1+\nu)^2\left(-\tilde{P}_2+(-\tilde{P}_1+\tilde{P}_2)*\cosh\left(\frac{x_2-x_3}{l_l}\right)\right)}=\exp\left(\frac{(x_1-x_3)}{l_l}\right). \quad \text{[Eq. S111]}$$

$$l_l*\ln\left(\frac{\frac{\overline{\sigma_{smooth}^{FB}}}{\frac{h_C B_C}{1-\nu^2}}+\frac{1}{2}(1+\nu)^2\left(\tilde{P}_1*\frac{1}{2}\right)}{-\frac{1}{2}(1+\nu)^2\left(-\tilde{P}_2+(-\tilde{P}_1+\tilde{P}_2)*\cosh\left(\frac{x_2-x_3}{l_l}\right)\right)}\right)=(x_1-x_3). \quad \text{[Eq. S112]}$$

$$-1 * l_l * \ln\left(\frac{\left(\frac{\overline{\sigma_{smooth}^{FB}}}{\frac{h_c B_c}{1-\nu^2}}+\frac{1}{2}(1+\nu)^2\left(\tilde{P}_1 * \frac{1}{2}\right)\right)}{\frac{-\frac{1}{2}(1+\nu)^2(-\tilde{P}_2)}{\left(1+\frac{(-\tilde{P}_1+\tilde{P}_2)}{-\tilde{P}_2} * \cosh\left(\frac{x_2-x_3}{l_l}\right)\right)}}\right) = (x_3 - x_1). \quad \text{[Eq. S113]}$$

We have assumed that $\left(-\tilde{P}_1 + \tilde{P}_2\right) < 0$ (see Eq. 5 and Fig. S5B). Given this assumption, one can easily show that for the left-hand side of Eq. S113 to be real, the following constraint must hold:

$$\frac{\overline{\sigma_{smooth}^{FB}}}{\frac{h_c B_c}{1-\nu^2}} > -\frac{1}{2}(1+\nu)^2\left(\tilde{P}_1 * \frac{1}{2}\right). \quad \text{[Eq. S114]}$$

For the example of Fig. S6A, Eq. S114 simply means that the horizontal gray dashed line must be greater than the minimum of the black curve

To demonstrate the non-monotonicity of the NPB domain size $(x_3 - x_1)$ with respect to substrate stiffness, take the partial derivative of Eq. S113 with respect to $l_l$. (Since $l_l$ is monotonic in $k$, we can prove non-monotonicity of $(x_3 - x_1)$ as a function of substrate stiffness by taking $\frac{\partial}{\partial l_l}$ of Eq. S113 instead of taking $\frac{\partial}{\partial k}$.)

$$\frac{\partial}{\partial l_l}(x_3 - x_1) = -\ln\left(\frac{\left(\frac{\overline{\sigma_{smooth}^{FB}}}{\frac{h_c B_c}{1-\nu^2}}+\frac{1}{2}(1+\nu)^2\left(\tilde{P}_1 * \frac{1}{2}\right)\right)}{\frac{-\frac{1}{2}(1+\nu)^2(-\tilde{P}_2)}{\left(1+\frac{(-\tilde{P}_1+\tilde{P}_2)}{-\tilde{P}_2} * \cosh\left(\frac{x_2-x_3}{l_l}\right)\right)}}\right) - \frac{\left(\frac{(-\tilde{P}_1+\tilde{P}_2)}{-\tilde{P}_2}\frac{x_2-x_3}{l_l}\sinh\left(\frac{x_2-x_3}{l_l}\right)\right)}{\left(1+\frac{(-\tilde{P}_1+\tilde{P}_2)}{-\tilde{P}_2} * \cosh\left(\frac{x_2-x_3}{l_l}\right)\right)}. \quad \text{[Eq. S115]}$$

For a stable fate pattern with a domain wall to exist, the first term on the right-hand side of Eq. S115 must be positive because the first term on the right-hand side of Eq. S115 is simply Eq. S113 divided by $l_l$. The second term on the right-hand side of Eq. S115 is strictly negative. The first term on the right-hand side of Eq. S115 corresponds to the change in NPB domain size due to changes in the decay length of the exponential; the second term corresponds to the change in NPB domain size due to changes in the magnitude of the exponential.

### Figure S1

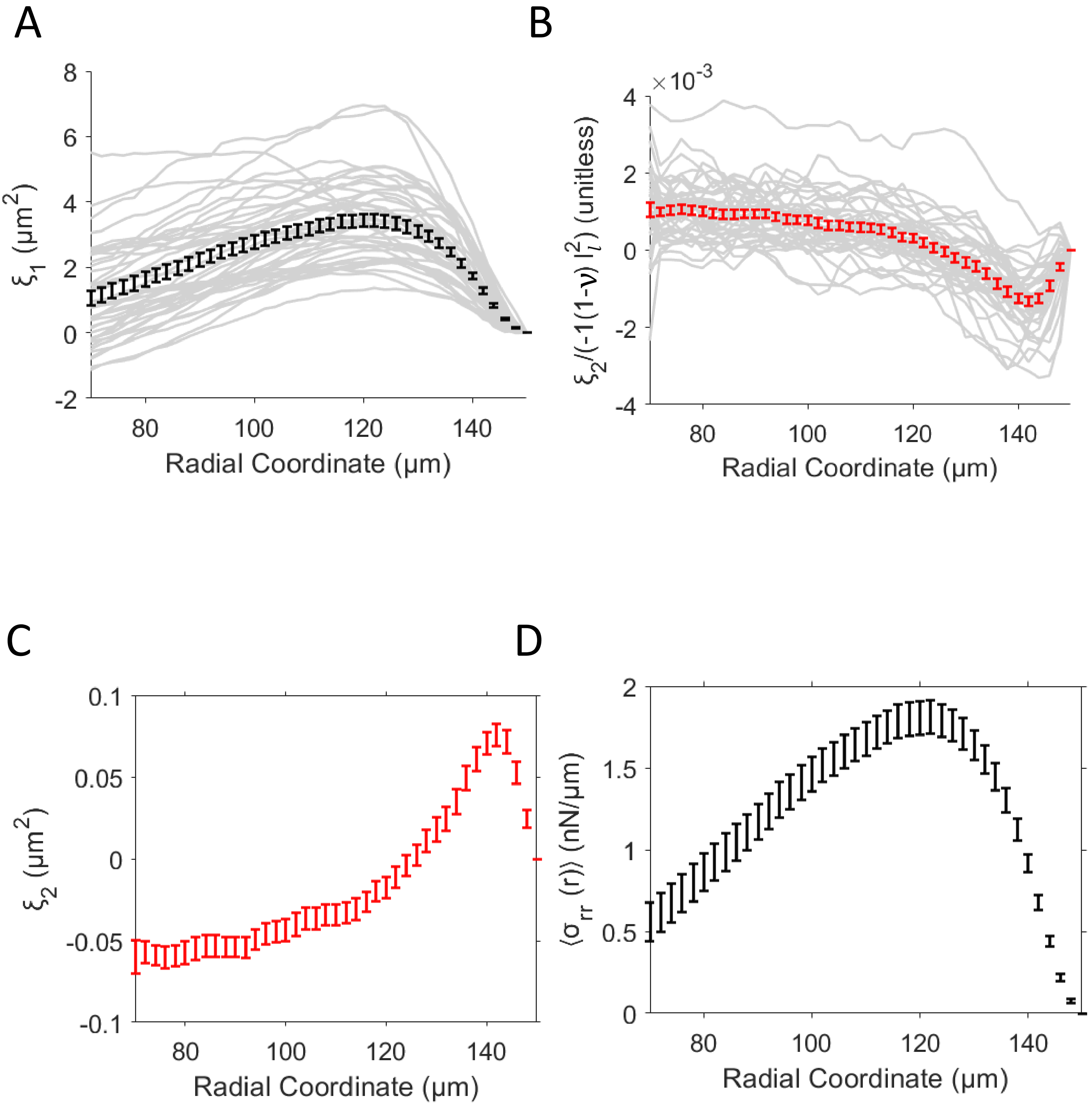
A
B
C
D
8
6
4
2
0
-2
80
100
120
140
$\xi_1$ ($\mu m^2$)
Radial Coordinate (μm)
×10^-3
4
2
0
-2
-4
$\xi_2/(-1(1-\nu)\, l_l^2)$ (unitless)
0.1
0.05
0
-0.05
-0.1
$\xi_2$ ($\mu m^2$)
2
1.5
1
0.5
0
$\langle\sigma_{rr}(r)\rangle$ (nN/μm)

### Figure S2

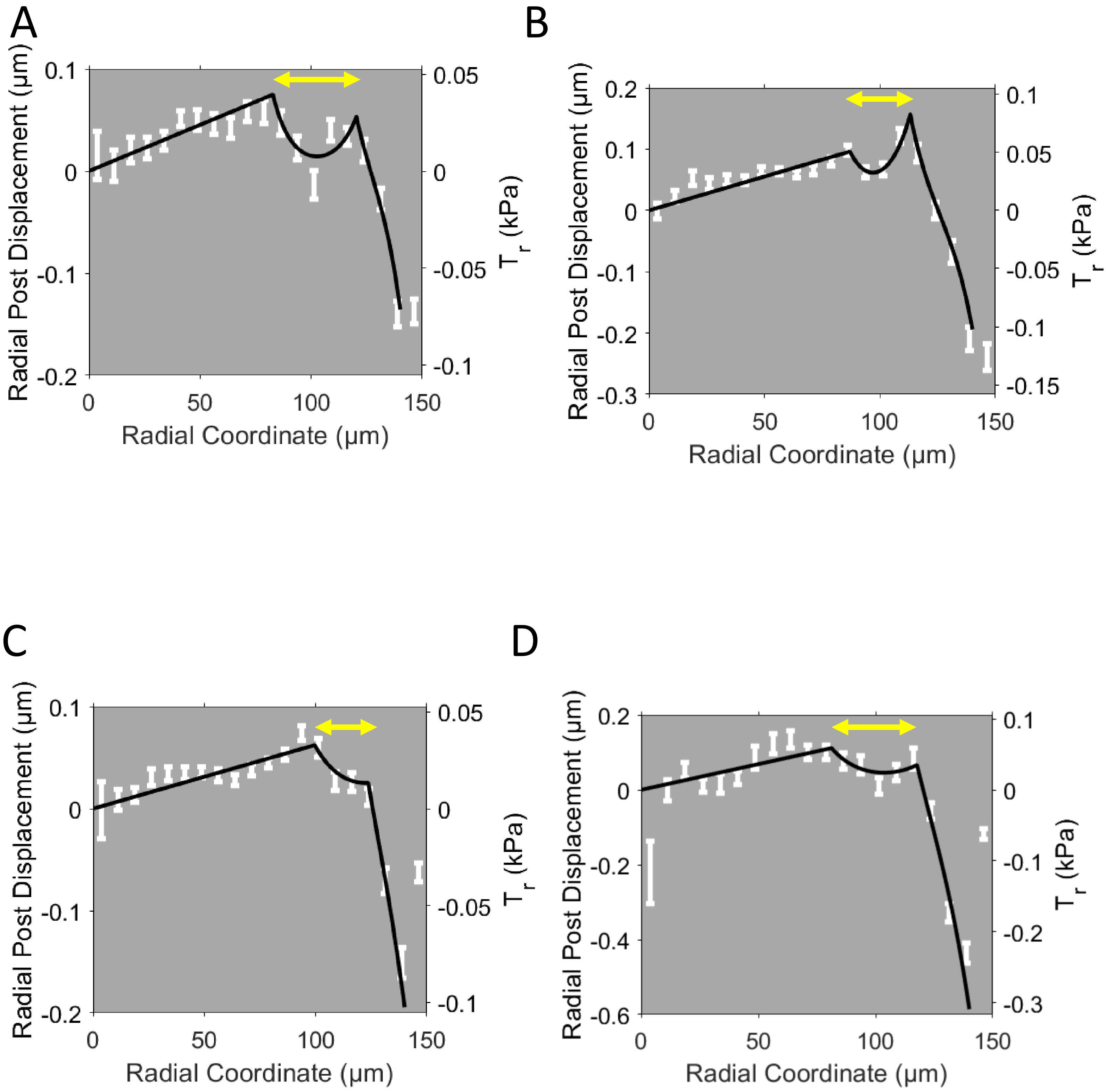
A
Radial Post Displacement (μm)
Radial Coordinate (μm)
$T_r$ (kPa)
B
Radial Post Displacement (μm)
Radial Coordinate (μm)
$T_r$ (kPa)
C
Radial Post Displacement (μm)
Radial Coordinate (μm)
$T_r$ (kPa)
D
Radial Post Displacement (μm)
Radial Coordinate (μm)
$T_r$ (kPa)

Figure S3

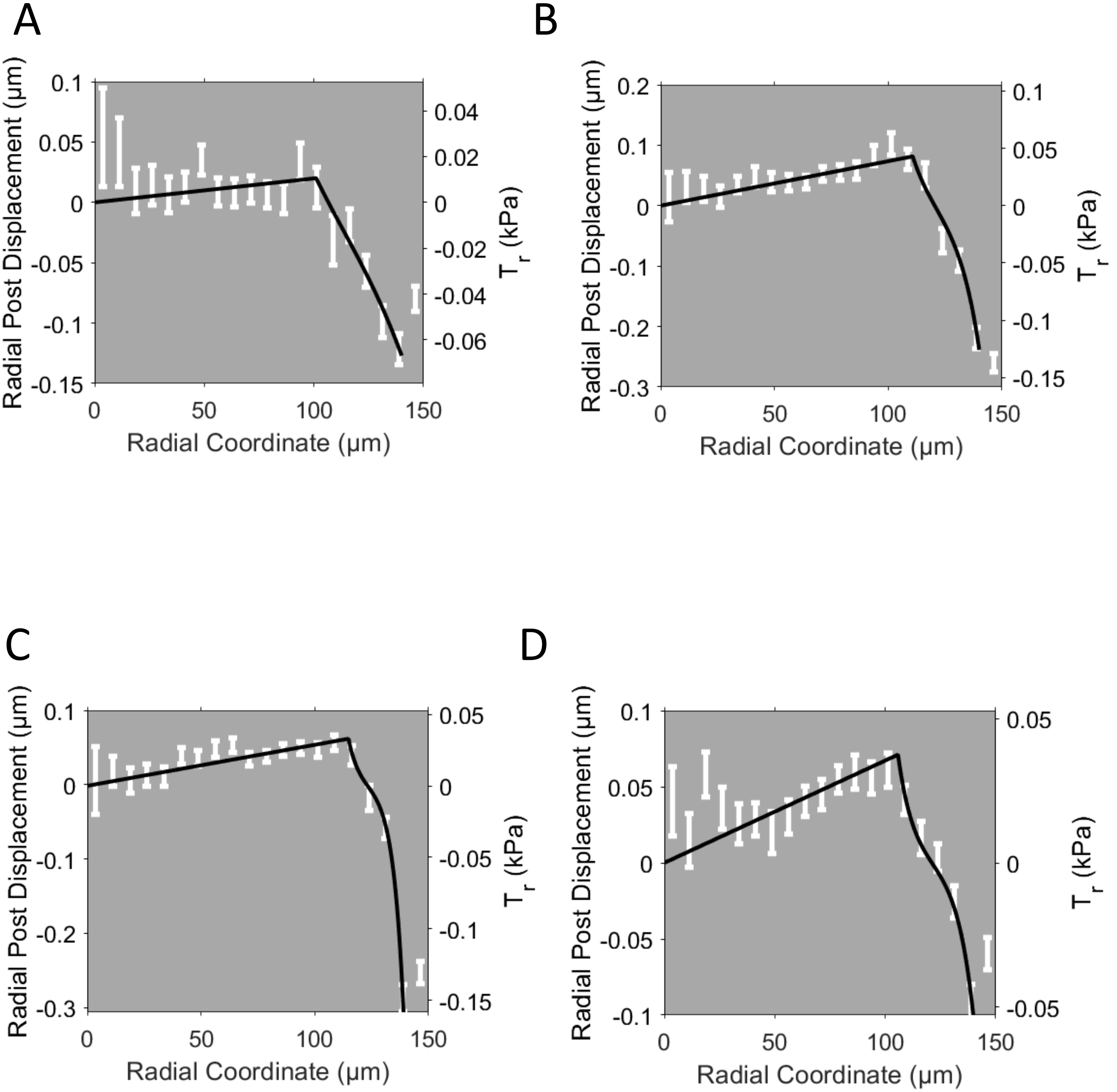
A
B
C
D
Radial Post Displacement (μm)
Radial Coordinate (μm)
$T_r$ (kPa)
0.1
0.05
0
-0.05
-0.1
-0.15
0.04
0.02
-0.02
-0.04
-0.06
0.2
-0.2
-0.3
-0.15
50
100
150

### Figure S4

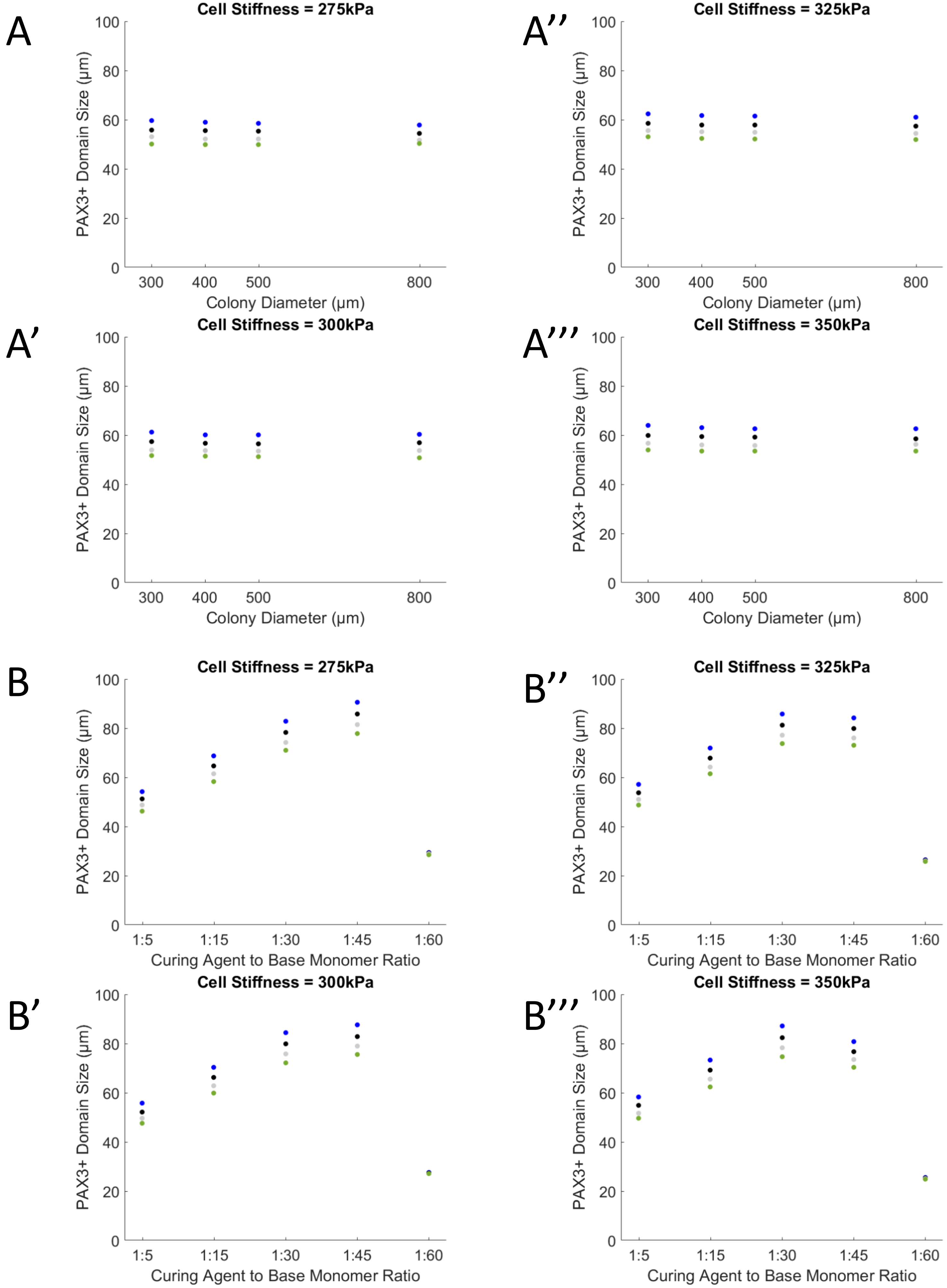
A
Cell Stiffness = 275kPa
PAX3+ Domain Size (μm)
Colony Diameter (μm)
A''
Cell Stiffness = 325kPa
PAX3+ Domain Size (μm)
Colony Diameter (μm)
A'
Cell Stiffness = 300kPa
PAX3+ Domain Size (μm)
Colony Diameter (μm)
A'''
Cell Stiffness = 350kPa
PAX3+ Domain Size (μm)
Colony Diameter (μm)
B
Cell Stiffness = 275kPa
PAX3+ Domain Size (μm)
Curing Agent to Base Monomer Ratio
B''
Cell Stiffness = 325kPa
PAX3+ Domain Size (μm)
Curing Agent to Base Monomer Ratio
B'
Cell Stiffness = 300kPa
PAX3+ Domain Size (μm)
Curing Agent to Base Monomer Ratio
B'''
Cell Stiffness = 350kPa
PAX3+ Domain Size (μm)
Curing Agent to Base Monomer Ratio

### Figure S5

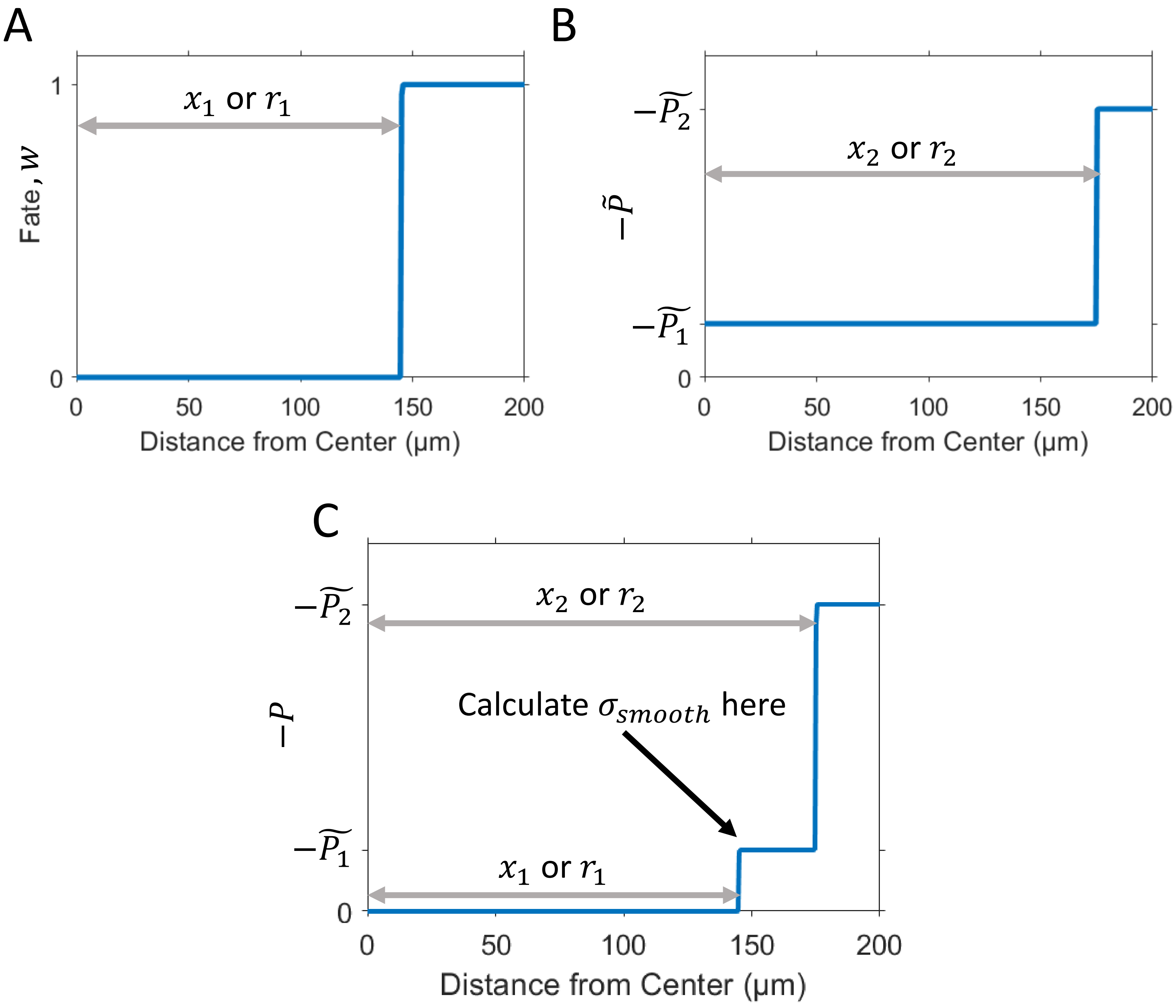

A
B
C
Fate, w
1
0
$x_1$ or $r_1$
Distance from Center (μm)
0
50
100
150
200
$-\tilde{P}$
$-\widetilde{P_2}$
$-\widetilde{P_1}$
$x_2$ or $r_2$
$-P$
Calculate $\sigma_{smooth}$ here

### Figure S6

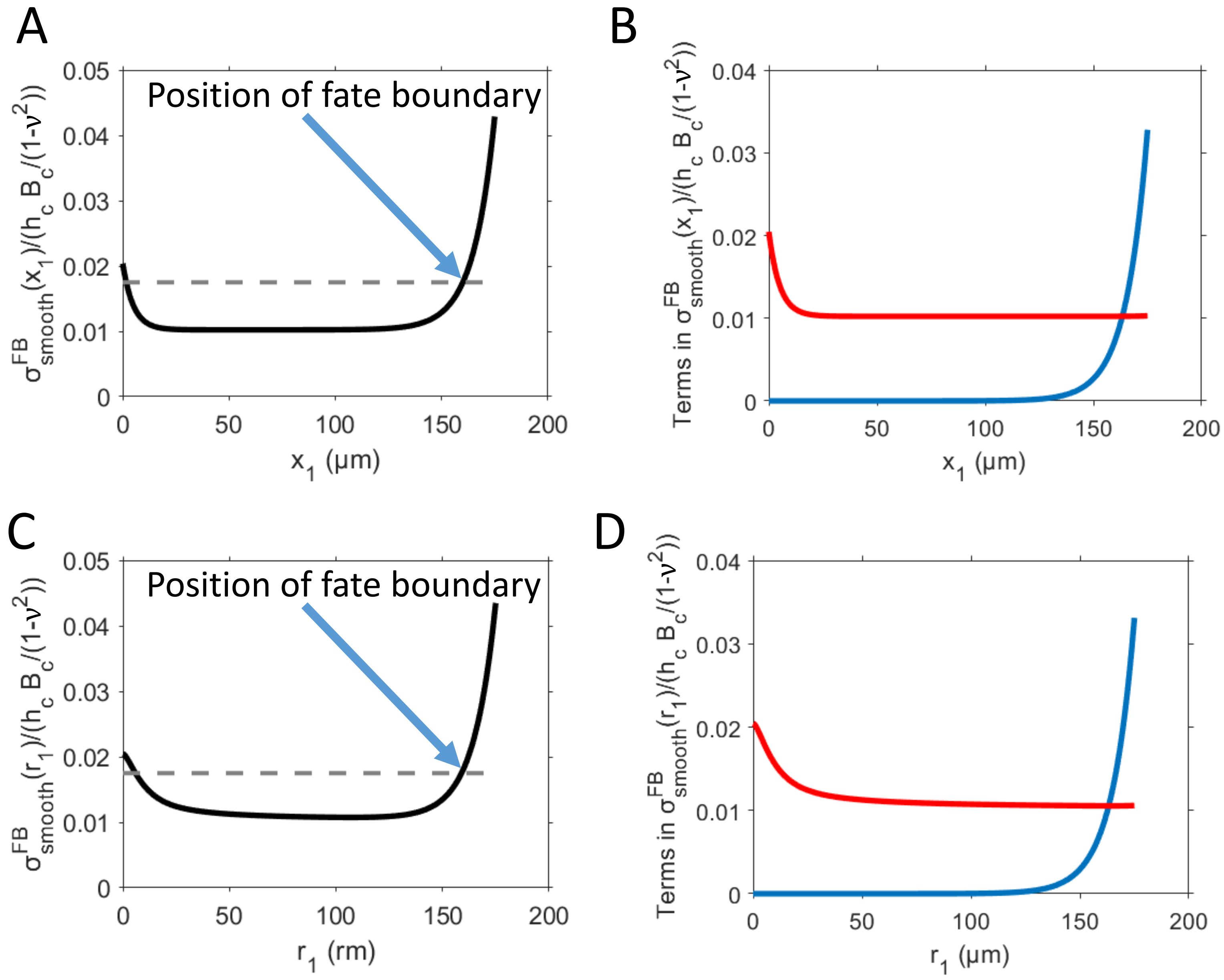
A
B
C
D
Position of fate boundary
Position of fate boundary
x1 (µm)
r1 (rm)
r1 (µm)
Terms in σ FB smooth(x1)/(hc Bc/(1-ν2))
Terms in σ FB smooth(r1)/(hc Bc/(1-ν2))

**Table S1**

| Number | $\tilde{\sigma}\left(\frac{\text{nN}}{\mu\text{m}}\right)$ | $l_l{}'(\mu\text{m})$ | $P'$ |
|---|---|---|---|
| 1 | 0.40 | 24.9 | -0.01 |
| 2 | 1.78 | 6.3 | -0.23 |
| 3 | 0.84 | 4.4 | -0.20 |
| 4 | 1.25 | 4.1 | -0.37 |
| 5 | 1.42 | 2.4 | -0.82 |
| 6 | 0.78 | 5.0 | -0.14 |
| 7 | 0.68 | 5.9 | -0.11 |
| 8 | 0.92 | 7.7 | -0.16 |
| 9 | 0.81 | 9.2 | -0.07 |
| 10 | 0.86 | 6.2 | -0.12 |
| 11 | 0.83 | 4.9 | -0.15 |
| 12 | 1.34 | 6.4 | -0.17 |
| 13 | 0.84 | 6.1 | -0.14 |
| 14 | 0.62 | 4.4 | -0.15 |
| 15 | 0.36 | 4.7 | -0.08 |
| 16 | 1.21 | 4.2 | -0.27 |
| 17 | 1.36 | 3.7 | -0.38 |
| 18 | 0.94 | 5.0 | -0.16 |
| 19 | 1.29 | 4.7 | -0.30 |
| 20 | 0.81 | 6.3 | -0.13 |
| 21 | 0.58 | 5.2 | -0.10 |
| 22 | 0.68 | 6.5 | -0.06 |
| 23 | 0.81 | 3.5 | -0.26 |
| 24 | 0.55 | 4.8 | -0.13 |
| 25 | 1.12 | 4.8 | -0.20 |
| 26 | 0.64 | 6.6 | -0.07 |
| 27 | 1.41 | 5.0 | -0.26 |

| | | | |
|---|---|---|---|
| 28 | 1.08 | 6.6 | -0.13 |
| 29 | 0.76 | 9.2 | -0.06 |
| 30 | 1.52 | 4.2 | -0.37 |
| 31 | 0.94 | 9.7 | -0.05 |
| 32 | 1.43 | 5.9 | -0.17 |
| 33 | 0.90 | 9.1 | -0.07 |
| 34 | 0.97 | 6.4 | -0.12 |
| 35 | 0.86 | 9.2 | -0.08 |
| 36 | 0.42 | 7.3 | -0.05 |
| 37 | 0.41 | 4.9 | -0.11 |
| 38 | 0.48 | 12.9 | -0.02 |
| 39 | 0.82 | 5.2 | -0.14 |
| 40 | 0.80 | 7.9 | -0.06 |
| 41 | 0.41 | 7.8 | -0.04 |
| 42 | 1.52 | 3.2 | -0.52 |
| Median | 0.84 | 5.9 | -0.14 |

**Table S2**

| Number | $\tilde{\sigma}\left(\frac{\text{nN}}{\mu\text{m}}\right)$ | $l_l''(\mu\text{m})$ | $r'(\mu\text{m})$ | $P'$ | $P''$ |
|---|---|---|---|---|---|
| 1 | 0.40 | >100 | 87 | 0.0002 | -0.01 |
| 2 | 1.78 | 8.2 | 105 | 0.0022 | -0.15 |
| 3 | 0.84 | 5.3 | 101 | 0.0015 | -0.13 |
| 4 | 1.25 | 5.5 | 118 | 0.0019 | -0.22 |
| 5 | 1.42 | >100 | 127 | 0.0016 | -0.03 |
| 6 | 0.78 | 5.4 | 94 | 0.0014 | -0.12 |
| 7 | 0.68 | 6.4 | 98 | 0.0022 | -0.10 |
| 8 | 0.92 | 8.2 | 84 | 0.0028 | -0.15 |
| 9 | 0.81 | 12.2 | 87 | 0.0023 | -0.04 |
| 10 | 0.86 | 6.6 | 90. | 0.0023 | -0.10 |
| 11 | 0.83 | 6.7 | 113 | 0.0011 | -0.09 |

| | | | | | |
|---|---|---|---|---|---|
| 12 | 1.34 | 7.1 | 92 | 0.0026 | -0.14 |
| 13 | 0.84 | 9.3 | 109 | 0.0016 | -0.07 |
| 14 | 0.62 | 5.6 | 106 | 0.0018 | -0.10 |
| 15 | 0.36 | 7.2 | 106 | 0.0013 | -0.04 |
| 16 | 1.21 | 6.1 | 102 | 0.0018 | -0.14 |
| 17 | 1.36 | 4.4 | 117 | 0.0020 | -0.28 |
| 18 | 0.94 | 5.0 | 99 | 0.0017 | -0.17 |
| 19 | 1.29 | 4.9 | 101 | 0.0020 | -0.28 |
| 20 | 0.81 | 9.9 | 102 | 0.0025 | -0.06 |
| 21 | 0.58 | 7.8 | 104 | 0.0017 | -0.05 |
| 22 | 0.68 | >100 | 111 | 0.0009 | -0.01 |
| 23 | 0.81 | 3.6 | 90. | 0.0017 | -0.24 |
| 24 | 0.55 | 5.6 | 109 | 0.0008 | -0.10 |
| 25 | 1.12 | 4.9 | 97 | 0.0013 | -0.19 |
| 26 | 0.64 | 9.5 | 107 | 0.0008 | -0.04 |
| 27 | 1.41 | 8.2 | 117 | 0.0012 | -0.12 |
| 28 | 1.08 | >100 | 119 | 0.0010 | -0.02 |
| 29 | 0.76 | 15.1 | 95 | 0.0012 | -0.03 |
| 30 | 1.52 | 4.3 | 115 | 0.0011 | -0.31 |
| 31 | 0.94 | 52.6 | 105 | 0.0011 | -0.01 |
| 32 | 1.43 | 9.3 | 111 | 0.0015 | -0.08 |
| 33 | 0.90 | 9.5 | 77 | 0.0014 | -0.07 |
| 34 | 0.97 | >100 | 121 | 0.0006 | -0.02 |
| 35 | 0.86 | 13.1 | 103 | 0.0013 | -0.05 |
| 36 | 0.42 | 10.8 | 96 | 0.0014 | -0.03 |
| 37 | 0.41 | 5.7 | 104 | 0.0013 | -0.09 |
| 38 | 0.48 | 28.6 | 101 | 0.0004 | -0.01 |
| 39 | 0.82 | 6.0 | 112 | 0.0008 | -0.11 |
| 40 | 0.80 | >100 | 109 | 0.0013 | -0.01 |
| 41 | 0.41 | >100 | 103 | 0.0012 | -0.01 |

| | | | | | |
|---|---|---|---|---|---|
| 42 | 1.52 | >100 | 122 | 0.0014 | -0.02 |
| Median | 0.84 | 8.2 | 104 | 0.0014 | -0.08 |

**Table S3**

| Number | $\tilde{\sigma}\left(\frac{\text{nN}}{\mu\text{m}}\right)$ | $l_l''(\mu\text{m})$ | $r'(\mu\text{m})$ | $r''(\mu\text{m})$ | $P'$ | $P''$ | $P'''$ |
|---|---|---|---|---|---|---|---|
| 1 | 0.40 | 86.1 | 98 | 108 | 0.0002 | -0.02 | 0.00 |
| 2 | 1.78 | >100 | 101 | 123 | 0.0022 | 0.00 | -0.03 |
| 3 | 0.84 | 46.2 | 87 | 124 | 0.0016 | 0.00 | -0.02 |
| 4 | 1.25 | 7.7 | 117 | 127 | 0.0019 | -0.11 | -0.14 |
| 5 | 1.42 | 20.0 | 92 | 129 | 0.0022 | -0.01 | -0.05 |
| 6 | 0.78 | 7.2 | 93 | 116 | 0.0014 | -0.05 | -0.07 |
| 7 | 0.68 | 9.8 | 91 | 115 | 0.0024 | -0.04 | -0.05 |
| 8 | 0.92 | 15.3 | 81 | 118 | 0.0027 | -0.03 | -0.07 |
| 9 | 0.81 | 10.4 | 56 | 80. | 0.0022 | -0.01 | -0.05 |
| 10 | 0.86 | 6.2 | 90. | 108 | 0.0023 | -0.10 | -0.12 |
| 11 | 0.83 | 7.4 | 77 | 110. | 0.0018 | -0.03 | -0.07 |
| 12 | 1.34 | 10.9 | 89 | 114 | 0.0026 | -0.05 | -0.07 |
| 13 | 0.84 | 10.3 | 83 | 111 | 0.0023 | -0.04 | -0.06 |
| 14 | 0.62 | 12.6 | 99 | 121 | 0.0018 | -0.02 | -0.03 |
| 15 | 0.36 | >100 | 89 | 116 | 0.0014 | 0.00 | -0.01 |
| 16 | 1.21 | 4.6 | 101 | 114 | 0.0018 | -0.18 | -0.23 |
| 17 | 1.36 | 10.0 | 87 | 113 | 0.0022 | -0.02 | -0.07 |
| 18 | 0.94 | 8.0 | 98 | 129 | 0.0017 | -0.06 | -0.08 |
| 19 | 1.29 | 5.5 | 101 | 125 | 0.0020 | -0.21 | -0.22 |
| 20 | 0.81 | 12.0 | 81 | 104 | 0.0029 | -0.02 | -0.04 |
| 21 | 0.58 | 8.2 | 80. | 102 | 0.0019 | -0.01 | -0.05 |
| 22 | 0.68 | 16.0 | 77 | 110. | 0.0013 | -0.01 | -0.02 |
| 23 | 0.81 | 8.8 | 83 | 120. | 0.0018 | -0.03 | -0.05 |
| 24 | 0.55 | >100 | 81 | 126 | 0.0001 | 0.00 | -0.02 |
| 25 | 1.12 | 6.8 | 94 | 121 | 0.0014 | -0.09 | -0.11 |
| 26 | 0.64 | 52.3 | 72 | 121 | 0.0010 | 0.00 | -0.01 |
| 27 | 1.41 | >100 | 55 | 117 | 0.0006 | 0.00 | -0.02 |
| 28 | 1.08 | 32.9 | 64 | 119 | 0.0014 | 0.00 | -0.02 |
| 29 | 0.76 | 27.6 | 91 | 115 | 0.0012 | -0.01 | -0.01 |
| 30 | 1.52 | 5.4 | 97 | 112 | 0.0011 | -0.18 | -0.22 |
| 31 | 0.94 | 19.8 | 67 | 109 | 0.0020 | -0.01 | -0.02 |
| 32 | 1.43 | 14.7 | 83 | 107 | 0.0014 | -0.01 | -0.04 |
| 33 | 0.90 | 16.8 | 75 | 121 | 0.0014 | -0.02 | -0.03 |
| 34 | 0.97 | 16.2 | 86 | 121 | 0.0009 | -0.01 | -0.04 |
| 35 | 0.86 | 17.0 | 87 | 111 | 0.0014 | -0.02 | -0.04 |
| 36 | 0.42 | 45.2 | 91 | 125 | 0.0015 | 0.00 | -0.01 |
| 37 | 0.41 | 15.2 | 100. | 124 | 0.0013 | -0.01 | -0.03 |
| 38 | 0.48 | 22.9 | 60. | 101 | 0.0006 | -0.01 | -0.01 |

| | | | | | | | |
|---|---|---|---|---|---|---|---|
| 39 | 0.82 | >100 | 99 | 131 | 0.0009 | 0.00 | -0.03 |
| 40 | 0.80 | 40.8 | 99 | 109 | 0.0013 | 0.00 | -0.01 |
| 41 | 0.41 | 36.2 | 93 | 105 | 0.0013 | 0.00 | -0.01 |
| 42 | 1.52 | 42.1 | 64 | 120. | 0.0002 | 0.00 | -0.02 |
| Median | 0.84 | 15.6 | 88 | 116 | 0.0014 | -0.01 | -0.04 |

**Table S4**

| Number | $\tilde{\sigma}\left(\frac{\text{nN}}{\mu\text{m}}\right)$ | $BIC(M_1)$ | $BIC(M_2)$ | $BIC(M_3)$ | $BF(M_3, M_2)$ |
|---|---|---|---|---|---|
| 1 | 0.40 | -235. | -256. | -271. | 1150 |
| 2 | 1.78 | 1. | -237. | -244. | 24.2 |
| 3 | 0.84 | -13 | -323. | -323. | 1.16 |
| 4 | 1.25 | 379. | -314. | -309. | 0.088 |
| 5 | 1.42 | 213. | -262. | -307. | $6.0 \times 10^9$ |
| 6 | 0.78 | -132. | -263. | -276. | 744 |
| 7 | 0.68 | 139. | -297. | -289. | 0.017 |
| 8 | 0.92 | -7. | -138. | -164. | 497000 |
| 9 | 0.81 | -61. | -234. | -240. | 23.4 |
| 10 | 0.86 | -7. | -303. | -300. | 0.16 |
| 11 | 0.83 | -101. | -225. | -241. | 4040 |
| 12 | 1.34 | 8. | -223. | -247. | 173000 |
| 13 | 0.84 | -71. | -257. | -278. | 42000 |
| 14 | 0.62 | 617. | -350. | -355. | 11.0 |
| 15 | 0.36 | -68. | -307. | -300. | 0.025 |
| 16 | 1.21 | 2. | -293. | -308. | 2000 |
| 17 | 1.36 | 811. | -291. | -311. | 32700 |
| 18 | 0.94 | 37. | -281. | -277. | 0.12 |
| 19 | 1.29 | -11. | -250. | -242. | 0.022 |
| 20 | 0.81 | 188. | -264. | -272. | 54.8 |
| 21 | 0.58 | 124. | -311. | -318. | 31.3 |
| 22 | 0.68 | -96. | -298. | -299. | 1.97 |
| 23 | 0.81 | -28. | -278. | -310. | $6.5 \times 10^6$ |
| 24 | 0.55 | -175. | -342. | -351. | 73.8 |

| | | | | | |
|---|---|---|---|---|---|
| 25 | 1.12 | -84. | -294. | -294. | 0.92 |
| 26 | 0.64 | -238. | -357 | -365. | 43.6 |
| 27 | 1.41 | -35. | -314. | -324. | 139. |
| 28 | 1.08 | -164. | -292. | -291. | 0.49 |
| 29 | 0.76 | -97. | -270. | -275. | 12.8 |
| 30 | 1.52 | -60. | -302. | -291. | 0.004 |
| 31 | 0.94 | -186. | -256. | -270. | 1350 |
| 32 | 1.43 | -113. | -282. | -274. | 0.02 |
| 33 | 0.90 | -144. | -206. | -203. | 0.21 |
| 34 | 0.97 | -220. | -266. | -263. | 0.16 |
| 35 | 0.86 | -155. | -221. | -220. | 0.62 |
| 36 | 0.42 | -44. | -341. | -338. | 0.19 |
| 37 | 0.41 | 247. | -336. | -353. | 5040 |
| 38 | 0.48 | -261. | -272. | -265. | 0.03 |
| 39 | 0.82 | -150. | -309. | -305. | 0.12 |
| 40 | 0.80 | 138. | -294. | -285. | 0.010 |
| 41 | 0.41 | -11. | -347. | -339. | 0.019 |
| 42 | 1.52 | 691. | -377. | -372. | 0.08 |

**Table S5**

| Curing agent : base monomer | Thickness $h_s$ (µm) | Young's Modulus $E_s$ (kPa) |
|---|---|---|
| 1 : 5 | 40 | 3,000 |
| 1 : 15 | 81 | 1,200 |
| 1 : 30 | 101 | 560 |
| 1 : 45 | 120 | 120 |
| 1 : 60 | 134 | 30 |

**Table S6**

| | |
|---|---|
| Fig. 3E-F | $\alpha * \tilde{P}_1 * k * N * l_l^2 * (1+\nu) = -0.05; \nu = 0.43; w_{mid} = \frac{1}{2}; l_l^2 = (20\ \mu\text{m})^2;$ $\tilde{P}_1 = -0.01; \tilde{P}_2 = -0.05; \Delta r = (150 - 129)\ \mu\text{m};\ \sigma^* = 1.865 \frac{\text{nN}}{\mu\text{m}}; D\tau_w = 0.0125\ \mu\text{m}^2$ |

| | |
|---|---|
| Fig. 4A | $h_c = 5.0\ \mu m; \nu_s = 0.4; Y_a = \frac{(5\ \mu m)(300\ kPa)}{4\ \mu m^2}; w_{mid} = \frac{1}{2}; \Delta r = 25\ \mu m; \tilde{P}_1 = -0.02;$ <br> $\tilde{P}_2 = -0.1;$ <br> $B_c = 350\ kPa; \alpha = \frac{0.05}{h_c B_c * -\tilde{P}_1}; D\tau_w = 0.0125\ \mu m^2; h_s = 81\ \mu m; E_s = 2100\ kPa;$ <br> $\sigma^* = 0.53 * h_c * B_c * (-1) * \tilde{P}_1$ |
| Fig. 4C | $h_c = 5.0\ \mu m; \nu_s = 0.4; Y_a = \frac{(5\ \mu m)(300\ kPa)}{4\ \mu m^2}; w_{mid} = \frac{1}{2}; \Delta r = 25\ \mu m; \tilde{P}_1 = -0.02;$ <br> $\tilde{P}_2 = -0.1;$ <br> $B_c = 350\ kPa; \alpha = \frac{0.05}{h_c B_c * -\tilde{P}_1}; D\tau_w = 0.0125\ \mu m^2;$ $h_s$, $E_s$ from experiments; <br> $\sigma^* = 0.53 * h_c * B_c * (-1) * \tilde{P}_1$ |
| Fig. S4A-A''' | $h_c = 5.0\ \mu m; \nu_s = 0.4; Y_a = \frac{(5\ \mu m)(300\ kPa)}{4\ \mu m^2}; w_{mid} = \frac{1}{2}; \Delta r = 25\ \mu m; \tilde{P}_1 = -0.02;$ <br> $\tilde{P}_2 = -0.1;$ <br> $B_c = 350\ kPa;\ \alpha = \frac{0.05}{h_c B_c * -\tilde{P}_1}; D\tau_w = 0.0125\ \mu m^2; h_s = 81\ \mu m; E_s = 2100\ kPa$ <br> $\sigma^*(blue) = 0.52 * h_c * B_c * (-1) * \tilde{P}_1; \sigma^*(black) = 0.525 * h_c * B_c * (-1) * \tilde{P}_1;$ <br> $\sigma^*(gray) = 0.53 * h_c * B_c * (-1) * \tilde{P}_1; \sigma^*(green) = 0.535 * h_c * B_c * (-1) * \tilde{P}_1$ |
| Fig. S4B-B''' | $h_c = 5.0\ \mu m; \nu_s = 0.4; Y_a = \frac{(5\ \mu m)(300\ kPa)}{4\ \mu m^2}; w_{mid} = \frac{1}{2}; \Delta r = 25\ \mu m; \tilde{P}_1 = -0.02;$ <br> $\tilde{P}_2 = -0.1;$ <br> $B_c = 350\ kPa; \alpha = \frac{0.05}{h_c B_c * -\tilde{P}_1}; D\tau_w = 0.0125\ \mu m^2;$ $h_s$, $E_s$ from experiments <br> $\sigma^*(blue) = 0.52 * h_c * B_c * (-1) * \tilde{P}_1; \sigma^*(black) = 0.525 * h_c * B_c * (-1) * \tilde{P}_1;$ <br> $\sigma^*(gray) = 0.53 * h_c * B_c * (-1) * \tilde{P}_1; \sigma^*(green) = 0.535 * h_c * B_c * (-1) * \tilde{P}_1$ |
| Fig. S6 | $x_3\ or\ r_3 = 200\ \mu m; \nu = 0.43; kN = 0.5271 \frac{nN}{\mu m^3}; l_l = 10\ \mu m; \widetilde{P_1}$ <br> $= 0.02;\ \widetilde{P_2} = 0.1$ <br> (Grey dashed line): $\frac{\overline{\sigma^{FB}_{smooth}}}{h_c B_c (1-\nu^2)} = 0.0175$ |

**Table S7**

| Protein | Vendor | Catalog number | Dilution |
|---|---|---|---|

| | | | |
|---|---|---|---|
| **PAX6** | Abcam | ab78545 | 1:200 |
| **PAX3** | Novus | NBP1-32944 | 1:400 |
| **PAX3** | R&D systems | MAB2457-SP | 1:200 |
| **E-cadherin** | BD-Biosciences | 610181 | 1:100 |

**Table S8**

| | 300 µm | 400 µm | 500 µm | 800 µm |
|---|---|---|---|---|
| 300 µm | | 0.003 | 0.002 | 0.002 |
| 400 µm | 0.003 | | 0.16 | 0.34 |
| 500 µm | 0.002 | 0.16 | | 0.64 |
| 800 µm | 0.002 | 0.34 | 0.64 | |

**Table S9**

| | glass | 1:5 | 1:15 | 1:30 | 1:45 | 1:60 |
|---|---|---|---|---|---|---|
| glass | | 0.006 | 0.003 | 0.0004 | 0.00005 | 0.0001 |
| 1:5 | 0.006 | | 0.73 | 0.00006 | 0.0002 | 0.07 |
| 1:15 | 0.003 | 0.73 | | 0.0002 | 0.0008 | 0.13 |
| 1:30 | 0.0004 | 0.00006 | 0.0002 | | 0.18 | 0.0008 |
| 1:45 | 0.00005 | 0.0002 | 0.0008 | 0.18 | | |
| 1:60 | 0.0001 | 0.07 | 0.13 | 0.0008 | | |

#### Figure Captions (SI)

S1: **Estimates of $\sigma_{rr}$ from colonies on micropost substrates at day 4. A)** $\xi_1$ (see Eqs. S50, S51) versus radial coordinate. Gray lines: individual cell colonies. Black: mean $\pm$ SEM. **B)** $\frac{\xi_2}{-(1-\nu)l_l^2}$ (Eqs. S50, S52) versus radial coordinate. Gray lines: individual cell colonies. Red: mean $\pm$ SEM. **C)** $\xi_2$ (mean $\pm$ SEM). To compare the typical magnitudes of $\xi_1$ and $\xi_2$, we need to know the typical magnitude of $(1-\nu)l_l^2$. Based on Tables S2, S3, we assume that $l_l^2 \approx 100\ \mu\text{m}^2$. We additionally assume that the cell layer is nearly incompressible in 2D (*i.e.*, $\nu \approx 0.4$). Based on this rough estimate for $(1-\nu)l_l^2$, we find that $\xi_2(r) \ll \xi_1(r)$. **D)** Approximation of$\langle\sigma_{rr}(r)\rangle$ (mean $\pm$ SEM) based on $\xi_1(r)$ alone.

S2: **Examples of concentrically averaged radial post displacement profiles with three clearly identifiable domains. A-D)** Examples of colonies for which Model 3 is more likely than Model 2. Yellow double-headed arrow: extent of intermediate domain **A)** Colony 23; **B)** Colony 17; **C)** Colony 37; **D)** Colony 8. (See SI Section 3, Tables S1-S4)

S3: **Examples of concentrically averaged radial post displacement profiles with two domains. A-D)** Examples of colonies for which Model 3 is less likely than Model 2. **A)** Colony 38; **B)** Colony 32; **C)** Colony 30; **D)** Colony 15. (See SI Section 3, Tables S1-S4)

S4: **Predicted dependence of NPB domain width on colony diameter and substrate stiffness.** To fit the model to experimental data, we scan over two fit parameters: cell layer stiffness $B_c$ and $\sigma^*$. Each plot is at fixed cell layer stiffness. The distinct colors correspond to different values of $\sigma^*$. (See Table S6) **A-A''')** Model predicts that NPB domain size is approximately independent of colony diameter. **B-B''')** Model predicts that NPB domain size depends non-monotonically on substrate stiffness.

S5: **Procedure for calculating the smooth stress at the fate boundary if given fate boundary position.** If you repeat this procedure for all possible positions of the fate boundary, then you know the smooth stress at the fate boundary as a function of fate boundary position. This function can then be inverted to find the position of the fate boundary as a function of smooth stress at the fate boundary (see Section 6). **A)** Set the fate variable $w = 0$ for NP (distance from center less than $x_1$). Set the fate variable $w = 1$ for NPB (distance from center greater than $x_1$). **B)** Specify the size of the boundary cell, which determines the spatial profile of $\tilde{P}$**.** The boundary cell spans from ($x_2$ for stripe and $r_2$ for disk) to the colony boundary. **C)** Calculate the target strain $P$, which is the product of the functions in A, B. Calculate the smooth stress at the fate boundary ($x_1$ for stripe and $r_1$ for disk) based on this $P$.

**S6: Smooth stress at the fate boundary as a function of the fate boundary position.** For each of these plots, we only plot the smooth stress away from the outermost – because we are focusing on the mechanical effect of a highly contractile cell at the colony boundary. That is why the plots only extent to 175 μm from the center even though the colony half-length is 200 μm. **A)** Representative example of (non-dimensionalized) smooth stress at the fate boundary as a function of the fate boundary's position for a cell layer in a stripe geometry, from Eq. S101; see Table S6 for parameter values. In this case, the colony half-length is 200 μm, and the boundary cell extends 25 μm inward from the boundary. The gray dashed line represents a potential value of $\overline{\sigma_{smooth}^{FB}}$, the value of the smooth stress at the fate boundary as determined from Eq. S100. For this value of $\overline{\sigma_{smooth}^{FB}}$, the position of the fate boundary is at the outermost intersection between the gray line and the black curve (see Section 6). **B)** We can decompose the function in panel A into two contributions. The red contribution encodes the non-uniformity near the colony center; the blue contribution encodes the non-uniformity due to the boundary cell. See curly-braced terms in Eq. S101. **C-D)** Same plots as in panels A, B except that the cell layer is a disc rather than a stripe.

#### Table Captions (SI)

**S1** **Best-fit parameters for Model 1 (see Section 3) for each sample.** This model assumes that the cell layer is uniform in both contractility and stiffness. $\tilde{\sigma}$ is the stress at $r = 140$ µm as determined by methods in Section 2A; this serves as the boundary condition for the fit. $l_l{}'$ is the localization length from the fit. $P'$ is the target strain.

**S2** **Best-fit parameters for Model 2 (see Section 3) for each sample.** This model assumes two concentric domains exist. The interior domain is stiff. $\tilde{\sigma}$ is the stress at $r = 140$ µm as determined by methods in Section 2A; this serves as the boundary condition for the fit. $l_l{}''$ is the localization length for the outer domain. $r'$ is the position of the boundary between the two domains. $P'$ is the target strain of the interior domain, and $P''$ is that of the exterior domain.

**S3** **Best-fit parameters for Model 3 (see Section 3) for each sample.** This model assumes three concentric domains exist. The innermost domain is stiff. $\tilde{\sigma}$ is the stress at $r = 140$ µm as determined by methods in Section 2A; this serves as the boundary condition for the fit. $l_l{}''$ is the localization length for the central and outermost domains. $r'$ is the position of the boundary between the innermost domain and the central domain; $r''$ is the position of the boundary between the central domain and the outermost domain $P'$ is the target strain of the innermost domain; $P''$ is that of the central domain; $P'''$ is that of the outermost domain.

**S4** **Comparing probability of each model via Bayesian Information Criterion for each sample. Color code: Maize $\boldsymbol{BF(M_3, M_2) \geq 20}$; Blue $\boldsymbol{BF(M_2, M_3) \geq 20}$; no color if $\boldsymbol{\frac{1}{20} < BF(M_3, M_2) < 20}$.** See Fig S2 for examples for which the three-domain model is the most probable (**Maize**). See Fig S3 for examples for which the two-domain model is the most probable (**Blue**).

**S5** **Young's modulus and substrate thickness for each value of curing-agent-to-base-monomer ratio.** These modulus values are as determined by a tensile testing machine (see Methods). Thicknesses were measured by a stylus profilometer (see Methods). These substrate properties are inputs for the modeling in Fig. 4C and Fig. S4B-B'''.

**S6** **Parameter values for figures.** For Figs. 3, S4, S6, we list all relevant parameter values. Fig. 3E,F and Fig. S6 are for cell layers on microposts. Fig. S4 is of cell layers on finite-thickness substrates.

**S7** **List of primary antibodies with corresponding vendors and dilution.**

**S8** **Statistical comparison via Mann-Whitney U-Test for Fig. 4A.** Each matrix element is the p-value for a difference in medians between the two groups (for example, between the 400-$\mu m$-colony-diameter samples and the 500-$\mu m$-colony-diameter samples).

**S9** **Statistical comparison via Mann-Whitney U-Test for Fig. 4C.** Each matrix element is the p-value for a difference in medians between the two groups (for example, between the samples on 1:5 substrate and those on 1:15 substrate).
